# IMC-Denoise: a content aware denoising pipeline to enhance Imaging Mass Cytometry

**DOI:** 10.1101/2022.07.21.501021

**Authors:** Peng Lu, Karolyn A. Oetjen, Diane E. Bender, Marianna B. Ruzinova, Daniel A.C. Fisher, Kevin G. Shim, Russell K. Pachynski, W. Nathaniel Brennen, Stephen T. Oh, Daniel C. Link, Daniel L.J. Thorek

**Author notes:** These authors contributed equally to this work (Peng Lu, Karolyn A. Oetjen).

## Abstract

Imaging Mass Cytometry (IMC) is an emerging multiplexed imaging technology for analyzing complex microenvironments that has the ability to detect the spatial distribution of at least 40 cell markers. However, this new modality has unique image data processing requirements, particularly when applying this technology to patient tissue specimens. In these cases, signal-to-noise ratio for particular markers can be low despite optimization of staining conditions, and the presence of pixel intensity artifacts can deteriorate image quality and the subsequent performance of downstream analysis. Here we demonstrate an automated content-aware pipeline, IMC-Denoise, to restore IMC images. Specifically, we deploy a differential intensity map-based restoration (DIMR) algorithm for removing hot pixels and a self-supervised deep learning algorithm for filtering shot noise (DeepSNF). IMC-Denoise outperforms existing methods for adaptive hot pixel removal, and delivers significant image quality improvement and background noise removal to a diverse set of IMC channels and datasets. This includes a unique, technically challenging, human bone marrow IMC dataset; in which we achieve noise level reduction of 87% for a 5.6-fold higher contrast-to-noise ratio, and more accurate background noise removal with approximately two-fold improved F1 score. Our approach remarkably enhances both manual gating and automated phenotyping with cell-scale down-stream analysis on these complex data. We anticipate that IMC-Denoise will provide similar benefits in mass cytometry imaging domains to more deeply characterize the complex and diverse tissue microenvironment.

## 1 INTRODUCTION

Disease states are the result of a complex interplay of many different cell types interacting in close proximity in the context of often heterogeneous tissues. Traditional approaches to study these features at the tissue scale have been limited in the number of specific markers that can be acquired to robustly resolve distinct cell types. Flow cytometry, perhaps the most widely used technique to study cell populations and states in this milieu, requires single cell disaggregation of the tissue resulting in complete loss of spatial context [1, 2]. Highly multiplexed imaging provides a means to assess these events at cellular resolution in *situ*, with extensive protocol development in progress [3], including tissue-based cyclic immunofluorescence (*t*-CyCIF) [4], co-detection by indexing (CODEX) [5], Multiplexed Ion Beam Imaging (MIBI) [6, 7] and Imaging Mass Cytometry (IMC) [8]. In IMC, tissue sections are stained with a panel of metal-conjugated antibodies, and data is acquired by UV-laser raster ablation of the section in 1-micron pixels for cytometry by time-of-flight (CyTOF) mass analyzer. This novel imaging technology allows for the detection of more than 40 antigens simultaneously. It enables single-cell, spatially resolved, highly multiplexed analysis of solid tissues and provides essential information on the distribution of transcripts, proteins, and protein modifications within single cells, microenvironments, and entire tissues [8–17]. The pixel data is processed into an image, thereby allowing the visualization of phenotypes and incorporation of spatial information in subsequent analyses. These properties make it a unique tool for the evaluation of complex biological systems.

Despite the wide applications in pre- and clinical research using this state-of-the-art multiplexed imaging technique, there exist specific technical noise sources in IMC, which include hot pixels, channel spillover and shot noise [8–10, 15, 18, 19]. Hot pixels are concentrated areas of high counts which are uncorrelated with any biological structures. Putatively, these can result from deposition of metal-stained antibody aggregates. In IMC images, single hot pixels are the most common outliers, and small hot clusters with several consecutive pixels may also exist. Channel spillover refers to scenarios where the signal of a source channel contaminates a target channel or is correlated with such contamination. The spillover in IMC can occur from a variety of reasons, such as instrument properties (abundance sensitivity), isotopic impurities and oxidation. Finally, shot noise exists because of ion counting imaging processes, which are pixel-independent, signal-dependent and usually modeled as a Poisson process. Additionally, noise levels are related to multiple other factors, including variations in conjugated metal isotopes, antibody concentration and arrangement.

Together these noise sources deteriorate the image quality and distort downstream analyses of IMC data. Differing from traditional fluorescence-based imaging modalities, there are low background features with highly multiplexed channels and no read-out noises from imaging sensors in IMC. A number of studies have attempted to address the unique imaging data features of IMC. Hot pixels can be corrected by thresholding methods [10, 14, 15, 20]; however, due to the differences between marker channels and tissues, a threshold needs to be pre-set carefully in these methods. An inappropriate threshold may lead to unsatisfactory results. Post-acquisition methods [10,19] and a bead-based compensation workflow [18] have been proposed to correct the channel spillover phenomenon. However, spillover correction may not be necessary if the marker panel employed is well-designed and titrated; and the intensity of channel-overlapping signal is often weak [18]. Therefore, spillover can be neglected when using low concentrations of staining antibodies, which however further lowers signal-to-noise ratio (SNR). To account for the impact of shot noise, MAUI [7, 19] and a semi-automated Ilastik-based method [21] have been used for background noise removal. These approaches require finely tuned parameters or manually annotated background regions, requiring preprocessing expertise. The results may be deteriorated by high noise levels for channels with weak signal. In tissues with low marker signals, highly intermixed cell populations, or difficult immunostaining, such as bone marrow specimens that must be subjected to additional processing for bone decalcification, defining thresholds can be time consuming with high inter-user subjectivity.

In the present work we develop and apply IMC-Denoise, a content aware denoising pipeline to enhance IMC images through an automated process. To account for the two major noise sources in this modality, hot pixels and shot noise, IMC-Denoise invokes novel algorithms for differential intensity map-based restoration (DIMR) and self-supervised deep learning-based shot noise filtering (DeepSNF). We demonstrate the flexibility and effectiveness of the proposed pipeline on publicly available IMC datasets of pancreatic cancer [10], breast cancer [12], a MIBI dataset [19], and a technically challenging unique human bone marrow dataset. We benchmark our approach against existing hot pixel removal methods [10, 14, 15, 20] and other advanced biomedical imaging denoising algorithms, such as non-local means filtering (NLM) [22], batch matching and 3D filtering (BM3D) [23] and Noise2Void (N2V) [24], which is used in IMC here for the first time. We demonstrate that the image formation model derived IMC-Denoise pipeline produces image quality enhancements that are best in class and leads to improved downstream analysis, with limited manual user manipulation. We provide this tool to augment studies that seek to more deeply characterize the complex and diverse tissue microenvironment.

## 2 RESULTS

### 2.1 IMC-Denoise principle

The general principle of IMC-Denoise is schematized in Fig. 1a and Supplementary Notes 1. To account for hot pixels and shot noise, an accurate IMC imaging joint model is built as Eq. (1), by considering ion counting imaging as a Poisson process (Supplementary Notes 1.1).

**Figure 1.**
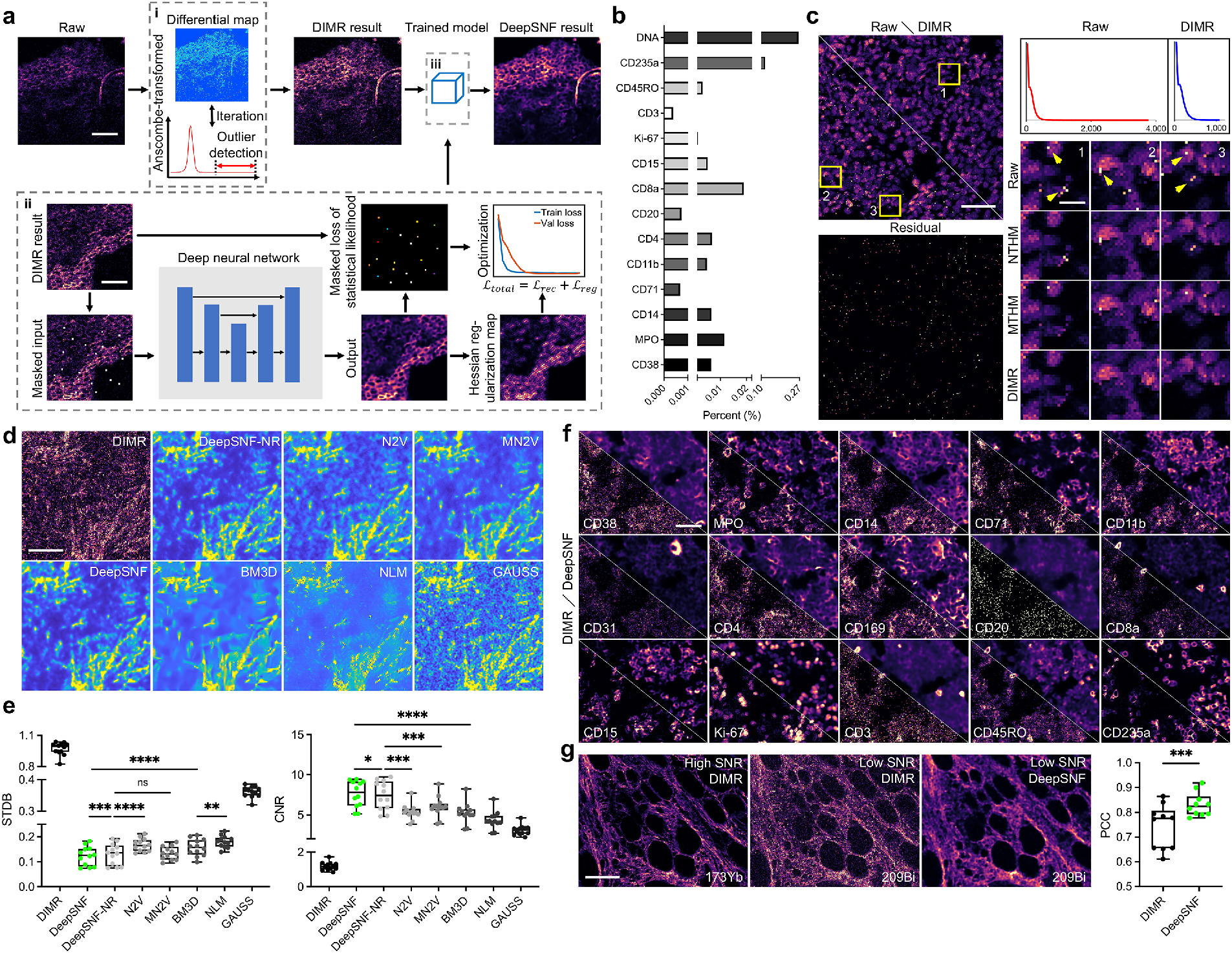
General principle and validation of IMC-Denoise on the human bone marrow IMC dataset. (a) Schematic of IMC-Denoise: (i) The DIMR algorithm: After the Anscombe transformation, the difference maps calculated from the raw image are operated to form a histogram. The outliers are detected based on this histogram and removed by a 3×3 median filter, iteratively. (ii) The training phase of the self-supervised DeepSNF algorithm: In the hot pixel corrected images, several pixels are randomly selected and masked. The hot pixel corrected images before and after the masking are set as the outputs and inputs of a deep neural network, respectively. Statistics-derived I-divergence on the masked pixels combined with the Hessian regularization on all the pixels is set as the loss function to guarantee the optimal denoising performance. (iii) The prediction phase of the DeepSNF algorithm: the hot pixel corrected IMC images are fed into the trained network to account for the shot noise. (b) The fractions of detected hot pixels by DIMR in selected channels. (c) DIMR removes hot pixels in DNA intercalator channel effectively. Left: Comparison of the raw and DIMR-processed images; and the difference between the images, in which Residual corresponds to the detected hot pixels. Upper right: The corresponding histograms of the raw and DIMR-processed images. Lower right: Comparisons between the raw, NTHM, MTHM and DIMR processed images. (d) Visual inspection of DeepSNF and other statistics-based denoising algorithms on a Collagen III-labeled IMC image. (e) DeepSNF performs significantly better than other algorithms (*n*=12) on denoising Collagen III-labeled IMC images in terms of STDB and CNR. (f) Visual inspection of DeepSNF denoised IMC images labeled with other markers. (g) DeepSNF improves the Pearson correlations between Collagen III-labeled IMC images with low and high SNR significantly (*n*=10). Scale bar: (a) Upper: 100 *µ*m, lower: 125 *µ*m. (c) Left: 75 *µ*m, right: 8 *µ*m. (d) 50 *µ*m. (f) 45 *µ*m. (g) 100 *µ*m.

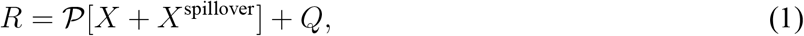

where *R* is the raw image, *X* the “clean” signal, *X*^spillover^ the spillover signals without noise, *P*[*x*] the Poisson noise with mean *x*, and *Q* the hot pixels. Normally, the term *X*^spillover^ in Eq. (1) can be omitted if the spillover is limited.

In IMC-Denoise, the DIMR algorithm (Fig. 1a(i) and Supplementary Notes 1.2) builds differential maps to detect the hot pixels by comparing adjacent pixels in a 3 × 3 sliding window, as hot pixels are local maxima. The Anscombe transformation [25] is applied to the raw image *R* followed by background removal of intensities lower than 4, so that the difference between adjacent pixels, *D*_*i*_, can be feasibly approximated as a generalized Gaussian distribution [26], where *i* is the neighbour index in the sliding window (*i* ∈ {1, 2, …, 8}). Additionally, as with all biomedical imaging acquisition, in IMC datasets the tissue or background pixels should be continuous. Under these conditions, for a specific pixel *p* there must exist several 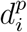 close to the mean *µ*_*i*_ of its corresponding distribution *D*_*i*_, except in the presence of a hot pixel. To unmix outliers from normal pixels, we consequently calculate the distances between 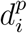 and *µ*_*i*_ as 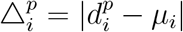 and sort 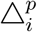 for *i* ∈ {1, 2, …, 8}. Then, the 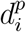 corresponding to the first *n* smallest 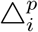 are summed, and the results from all pixels form a new distribution, *T*_*n*_. Compared to those in the distributions *D*_*i*_, the hot pixels move beyond the right tail of *T*_*n*_, while the normal relevant pixels move towards its center (Supplementary Note 1.2.1). To robustly detect the outliers, the kernel density estimation algorithm [27] is applied to *T*_*n*_ afterwards (Supplementary Note 1.2.2). On the fitted curve (*x, ĝ*_*h*_(*x*)), a threshold point *x*_*T*_ is defined so that any points *x > x*_*T*_ are considered as outliers and filtered by a 3 × 3 median filter. Because outliers are located beyond the right tail of *T*_*n*_, it is reasonable to set *x*_*T*_ when 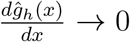, which means the current distribution ends. Likewise, the shape of the distribution should not change from convex to concave on the right tail. Thus, it is also reasonable to set *x*_*T*_ when 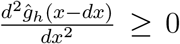 and 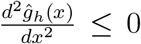. We operate DIMR for multiple iterations to adequately remove hot pixels until no outliers are detected. The hot pixel removed images are transformed to their original scales with the direct algebraic inverse Anscombe transformation [28]. Normally, *n* = 4 and the iteration number is set as 3. In addition, use of the median 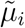 of distribution *D*_*i*_ is a robust estimation of the mean *µ*_*i*_. The DIMR algoirthm is summarized as Supplementary Algorithm 1.

After hot pixel removal, the imaging model is simplified as: *R* = *𝒫* [*X*], for which we have developed DeepSNF (Fig. 1a(ii, iii) and Supplementary Note 1.3) to account for the ion noise in IMC images. By combining Poisson statistics and detection theory, I-divergence [29] is derived as the loss function to enable the maximized likelihood estimation for the denoising task. Unlike traditional imaging methods for which noise-free training label images can be generated, commonly with long exposures, the image formation process in IMC requires laser ablation. Thus, a tissue can only be imaged once in IMC. Autofluorescence artifacts in immunofluorescence (IF) images, and the tedious and potentially interfering processes for consecutive IF and IMC imaging, are further confounds. Therefore, conventional supervised denoising approaches [30–32] or Noise2Noise [33] are not available here.

We overcome these limitations by applying a self-supervised approach inspired by Noise2Void [24] and Noise2Self [34]. This approach randomly masks several pixels in the DIMR-processed hot pixel-removed images by a stratified sampling strategy. Subsequently, the manipulated images are set as the inputs of the network and the hot pixel removed ones are the outputs. For this construct, the self-supervised training is approximately equivalent to a supervised learning process (Supplementary Note 1.3.1). The network follows U-Net [35] structure with Res-Blocks [36] to enable high quality training and prediction (Methods, Supplementary Fig. 11). Notably, the last activation function of the network is set as softplus (log(1 + exp(*x*))) to restrict non-negativity for the images. Nevertheless, the denoising performance is still sub-optimal, due to neglected information of the masked pixels and partially utilized pixels in the self-supervised strategy. To further boost DeepSNF, the Hessian regularization [37–39] is applied in the loss function with the continuity between biological structures *a priori* (Supplementary Note 1.3.2). Overall, the loss function of DeepSNF is summarized as Eq. (2).

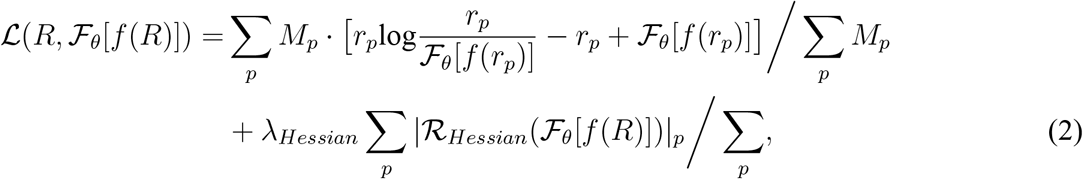

where *ℱ*_*θ*_ represents the learnt weights of the network, *f* demonstrates the random pixel masking approach, *r*_*p*_ the *p*-th pixel of the hot pixel removed training set *R, M* the pixel mask (*M* ∈ {0, 1}), *ℛ*_*Hessian*_ the Hessian operator, *λ*_*Hessian*_ the regularization parameter and *p* the pixel index. Here, the pixel *p* is masked only when *M*_*p*_ = 1. Prior to training and prediction, the images are normalized between 0 and 1 by a percentile-normalization approach (Supplementary Note 1.3.3). As validated by simulation (Supplementary Note 3.4.2), *λ*_*Hessian*_ is empirically set as 3e-6 to balance the data fidelity and regularization. The DeepSNF algorithm is summarized as Supplementary Algorithm 2.

### 2.2 Validation of IMC image quality improvement

We initially tested our DIMR algorithm on selected markers of a human bone marrow dataset. Here, inherently high autofluorescence and tissue features (fragile haematopoietic stroma intermixed with dense cortical bone) excluded other spatial biology methods, even after substantial pre-processing. Fig. 1b enumerates the proportion of hot pixels detected by DIMR for each marker. We then selected DNA intercalator, CD235a and MPO (Fig. 1c, Supplementary Figs. 12 and 13) to evaluate DIMR due to their high hot pixel density. By comparing the images and the corresponding histograms, hot pixels had been effectively eliminated by DIMR.

We further compared DIMR with two recent hot pixel removal methods, neighbour-based threshold hot pixel removal method (NTHM) [14, 15, 20] and median-based threshold hot pixel removal method (MTHM) [10] with default parameters, to benchmark its performance (Supplementary Note 2.1, Supplementary Algorithms 3 and 4). From the results, all three methods could remove spurious signal, but their performances varied from each other. To quantitatively evaluate these methods, we utilized *t*-CyCIF data [40] to generate simulated IMC images (Supplementary Note 3.1) with a range of noise levels and hot pixel densities. The three methods were then applied on the simulated datasets, and root mean squared errors between the hot pixel-free and processed images were set as the metric to evaluate the accuracy of hot pixel removal (Supplementary Note 3.2). Note that in simulations the thresholds of NTHM and MTHM were manually tuned to guarantee their optimal performances, while DIMR was configured automatically. The simulation results indicate DIMR is the best performer among the three methods (Supplementary Note 3.3). In fact, the threshold of NTHM requires contextual adjustment as different tissues and channels may have different scales. Moreover, this method is not efficient at removing consecutive hot pixels. MTHM is not locally adaptive and may overlook hot pixels with similar intensity to that of normal pixels; or erroneously remove normal pixels located at the border between tissues and background. Use of a lower search range or threshold for MTHM may also generate false negatives. In comparison, the outlier detection of DIMR is based on overall image statistics. Therefore, no manual threshold adjustment is required for images with different intensity scales, and a higher detection sensitivity is achieved even for hot pixels with lower intensities. These features along with the simulation data results demonstrate the versatility and accuracy of DIMR. The automated DIMR approach also results in the additional benefit of moderately improved cell segmentation, through robust removal of artifacts caused by hot pixels (Supplementary Fig. 14).

Next, we benchmarked the performance of DeepSNF along with DIMR and other statistics-based denoising methods including a Gaussian filter with standard deviation of 1 (GAUSS), NLM, BM3D, N2V, modified N2V (MN2V) and DeepSNF with no regularization (DeepSNF-NR) (Supplementary Notes 2.2 and 2.3) on the simulated dataset (Supplementary Note 3.4) and on IMC images labeled with Collagen III, CD31, CD34 and CD3 from the human bone marrow dataset (Fig. 1d and Supplementary Fig. 15). First we visually assessed images with different processing approaches for their overall appearance and in particular for retention of fine cell-level details. We found all the algorithms enhanced the DIMR data even though variant performances were achieved. GAUSS lowers the noise level by sacrificing resolution. NLM is effective at background denoising but does not account adequately for the noise components of signal. BM3D improves NLM further by its cooperative denoising procedure. However, we found it tended to over-smooth foreground and distorted cell shapes. N2V always generates artifacts because of inappropriate noise model. DeepSNF-NR performs better than MN2V because the Anscombe transformation in MN2V may generate some bias for extremely low counts; both of which are better than GAUSS, NLM and BM3D. DeepSNF further enhances these results by mitigating the discontinuities in the DeepSNF-NR output, and furthermore retains cell morphology features.

We quantitatively compared the differently processed images across a range of different characterization methods. Assessment of peak SNR (PSNR) and structural similarity (SSIM) [41] (Supplementary Note 3.2) were computed from the simulated data, and the standard deviation of background (STDB) and contrast-to-noise ratio (CNR) (Methods) were utilized for the IMC images labeled with Collagen III. All results indicated DeepSNF enables the optimal denoising performance among these algorithms (Fig. 1e and Supplementary Note 3.4). In particular, the noise level (STDB) decreased by 87% and CNR increased by 5.6-fold after DeepSNF (0.9938 to 0.1254 and 1.1749 to 7.8065, median value).

We further visually inspect the denoising results of IMC-Denoise on multiple datasets including human bone marrow images (Fig. 1f and Supplementary Fig. 16), human breast cancer (Supplementary Fig. 17), human pancreatic cancer (Supplementary Fig. 18) and a MIBI dataset (Supplementary Fig. 19). Image quality improvements that enhance image interpretation are apparent, in particular for low SNR channels. Two orthogonal staining approaches were pursued in order to provide further validation of these image quality improvements. Firstly, the same antibody was conjugated to two different metals and co-stained on the same tissue for detection in high and low sensitivity channels, without spillover. IMC-Denoise was employed on the low signal channel (209Bi) and was able to restore the image quality to match the high sensitivity channel (173Yb) with the Pearson correlation coefficient (PCC) improved as high as 0.16, as shown in representative images (Supplementary Fig. 20 and Fig. 1g). Similar conclusions can also be drawn from other channels with increased PCC by more than 0.48 and 0.35, respectively (Supplementary Fig. 21). Secondly, tissue sections stained with metal-conjugated antibodies (for CD3, CD4, CD61 and CD169) were probed with a fluorophore-conjugated secondary antibody for IF, individually. We then followed IF imaging by ablative-IMC (Supplementary Fig. 22). The additional handling and washing after IF imaging often leads to extremely low remaining metal isotope signal; however, enhancement in image quality can still be observed to restore the image to correlate to the IF. Specifically, the PCC quantitatively verified the image quality improvement of DeepSNF (CD3: 0.5557 to 0.7939, CD4: 0.4975 to 0.7793, CD61: 0.9096 to 0.9492, and CD169: 0.4481 to 0.7726).

### 2.3 IMC-Denoise enables background noise removal of IMC images and enhancement of IMC downstream analysis

We next evaluated the ability of DeepSNF in IMC-Denoise to remove background noise of IMC images. We first applied a single threshold per image, suggested in [21], for the images in Fig. 1g and Supplementary Figs. 17–19. Manual thresholding can pose inherent challenges in many tissues, particularly in bone marrow specimens with highly heterogeneous cell mixing and low marker signal. Visual inspection (Supplementary Figs. 23–26) reveals DeepSNF enhances background noise removal effectively. To fully evaluate the enhancement by DeepSNF, we manually annotated 15 images labeled with CD34 and 12 IMC images labeled with Collagen III (Fig. 2a). The single threshold-based method and semi-automated Ilastik-based method were applied on both DIMR and DeepSNF-processed CD34 and Collagen III images (DIMR_thresh, DeepSNF_thresh, DIMR_Ilastik and DeepSNF_Ilastik, respectively), and MAUI was only applied on DIMR images (MAUI). The results were compared with the manually annotated ground truths (Fig. 2b), and F1 score was set as the accuracy metric to quantitatively access the results (Fig. 2c). Overlaid masks and F1 scores for both markers indicated DeepSNF_Ilastik achieves the highest accuracy while DIMR_thresh is the weakest performer (CD34: 0.9143 to 0.3204, and Collagen III: 0.9434 to 0.5378, median value). Surprisingly, DeepSNF_thresh is a better method for background noise removal than the semi-automated DIMR_Ilastik (CD34: 0.9040 to 0.8716, and Collagen III: 0.9345 to 0.9108, median value), and its F1 score was improved by approximately twofold compared to DIMR_thresh. We infer that DeepSNF is capable of unmixing the signal and background, while the shot noise in DIMR images hinders the performances of the Ilastik-based method. MAUI was able to account for the background noise at the cost of false negative generation (CD34: 0.7824 and Collagen III: 0.7305, median value). Through a wide range of tested parameters we found that the accuracy of MAUI is demonstrably lower than DeepSNF_thresh (Supplementary Fig. 27). Furthermore, on MIBI data, an alternative highly multiplexed imaging technology, DeepSNF achieves similar denoising performance with MAUI (Supplementary Figs. 19 and 26). Overall, these findings indicate DeepSNF achieves robust background noise removal, which may enable tedious semi-automated approaches to be replaced by automated DeepSNF. Indeed, even the need for background thresholding can be discarded because the signal has been unmixed from background through DeepSNF.

**Figure 2.**
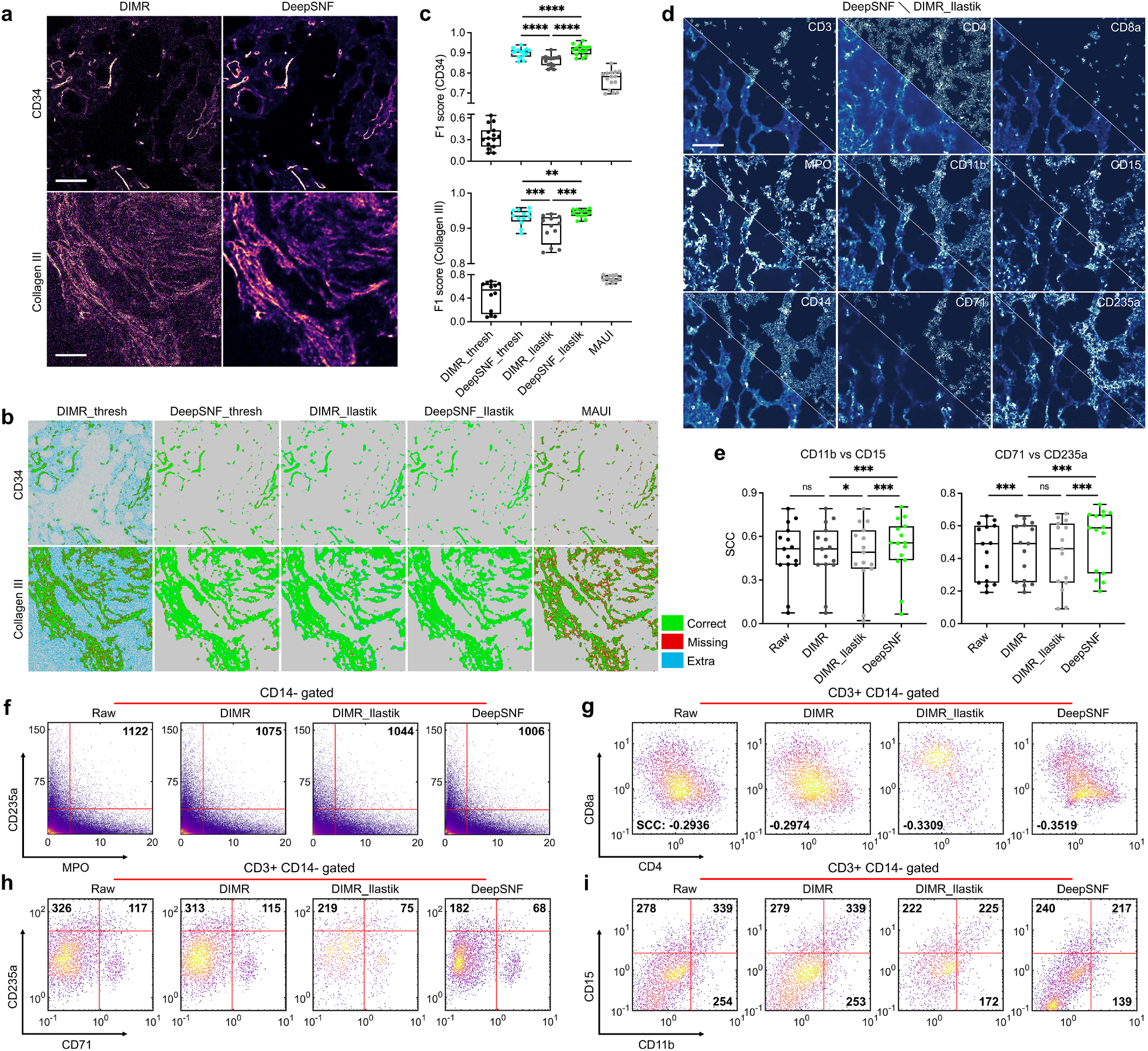
IMC-Denoise enables background noise removal and enhances downstream analysis of the human bone marrow IMC dataset. (a) Examples of DIMR and DeepSNF-processed IMC images labeled with CD34 and Collagen III. (b) Visual inspection of background removal results of DIMR and DeepSNF-processed images, in which DIMR_thresh and DeepSNF_thresh are binarized with the threshold value 1 directly, DIMR_Ilastik and DeepSNF_Ilastik are segmented by the Ilastik software package, and MAUI results are the DIMR images processed by the MAUI software package, respectively. Manual annotated images are served as ground truths. (c) After DeepSNF denoising, the background removal accuracy improves significantly in terms of F1 score, for both CD34 (*n*=15) and Collagen III-labeled images (*n*=12). Notably, DeepSNF_Ilastik achieves the highest accuracy, while DeepSNF_thresh performs better than all the background removal results from DIMR images. (d) Visual inspection of DeepSNF and DIMR_Ilastik-based denoising results on different markers-labeled IMC images. (e) SCC evaluation between markers with similar profiles, in which each dot corresponds to the SCC calculated from the single cell profiles in a single tissue slide (*n*=15). (f)–(i) Evaluations of denoising algorithms with manual gating strategies on single cell data. The numbers in (f), (h) and (i) are the cell counts of the corresponding ranges; and those in (g) are the SCC between CD4 and CD8a of the CD3+ CD14-gated cells. DIMR slightly enhances the single cell analysis over raw data, while DeepSNF further enhances the DIMR results and performs better than semi-automated DIMR_Ilastik-processing. Scale bar: (a) Top: 50 *µ*m, bottom: 35 *µ*m. (d) 107 *µ*m.

For downstream IMC analysis, we were curious to evaluate the impact of IMC-Denoise on single cell profiles. Using segmented cell masks, we extracted the cell intensities of CD38, MPO, CD14, CD71, CD11b, CD4, CD169, CD20, CD8a, CD15, CD3 and CD235a markers for 96232 cells in total (Methods). Please note that segmentation masks were identical for each comparison, using masks generated from DeepSNF, so that the impact of variability in segmentation algorithms can be neglected. In Supplementary Figs. 28 and 29a, the comparison of the single cell profiles of raw, DIMR and DeepSNF data show that DIMR corrects false positive data only, and DeepSNF corrects all cell profiles. Typically, larger mean positive marker expressions lead to lighter corrections by DeepSNF (Supplementary Fig. 29b). This follows from the logic that larger ion counts have lower shot noise levels.

Subsequently, we benchmarked the single cell data from the raw, DIMR, DIMR_Ilastik and DeepSNF-processed images (Fig. 2d and Supplementary Fig. 30a). The Spearman correlation coefficients (SCC) between the single cell profiles of myeloid (CD11b and CD15), erythroid (CD71 and CD235a) populations were first used to evaluate these methods (Fig. 2e and Supplementary Figs. 31, 32). The SCC between CD71 and CD235a increases after DIMR correction because of the high density of hot pixels in CD235a-labeled images, while the SCC between CD11b and CD15 remains almost unchanged after DIMR. By contrast, after DeepSNF operation the SCC for both pairs increase significantly, P<0.001. Interestingly, DIMR_Ilastik lowered the SCC for both the pairs. We deduce that the Ilastik-based approach removed unspecific staining signal which may lead to the SCC decrease. Next, manual gating approaches with prior knowledge of cell markers were applied to more deeply digest the impact of DIMR, DIMR_Ilastik and DeepSNF on IMC data (Fig. 2f–i and Supplementary Fig. 30b). For example, there should be no MPO, CD235a double positive cells on the CD14-cell subsets. In Fig. 2f, DIMR improved the raw data (from 1122 to 1075 counts) by correcting high hot pixel densities in both MPO and CD235a. Nevertheless, the double positive cells do not decrease further by DIMR_Ilastik (1044) or DeepSNF (1006), due to the low noise levels of these two markers (Supplementary Fig. 29b). The remaining double positive cells are likely related to segmentation artifacts between adjacent cells of different types.

As a second example, among T cells (CD3-positive, CD14-negative) CD4 and CD8a should be negatively correlated with each other, and myeloid and erythroid markers should be absent. In Fig. 2g, the SCC between CD4 and CD8a decrease slightly after DIMR correction (from -0.2936 to -0.2974), and further decrease after DIMR_Ilastik and DeepSNF operations. Notably, DeepSNF achieved lower SCC than DIMR_Ilastik (-0.3519 to -0.3319). Similarly, DIMR_Ilastik and DeepSNF further removed false positive myeloid (Fig. 2h) and erythroid markers (Fig. 2i) after the slight improvement of DIMR. However, DeepSNF decrease 41.6% false positive cells for myeloid and 31.6% for erythroid while DIMR_Ilastik decrease 31.3% and 28.9%, respectively. The manual gatings in Supplementary Fig. 30b indicate the same trends as well. Overall, as expected DIMR could enhance the single cell analysis to a limited extent. DeepSNF and DIMR_Ilastik enable further enhancement, and the former achieves better performance than the latter on this task.

### 2.4 DeepSNF in IMC-Denoise enhances automated cell phenotyping

Cell phenotype annotation plays a key role in tissue microenvironment analysis. Indeed, false annotation of cell phenotypes has the potential to lead to false biological or clinical conclusions. Hot pixel removal is normally conducted before automated cell phenotyping [14, 15, 17]. Therefore, we focused on whether DeepSNF in IMC-Denoise could impact phenotypic annotation of cell types. Here, a subset of markers from the human bone marrow dataset were used for phenotypic annotation, including CD38, MPO, CD14, CD71, CD11b, CD4, CD169, CD20, CD8a, CD15, CD3 and CD235a. We clustered the DIMR dataset by the K nearest neighbours(KNN)-based Jaccard graph construction [42] and the Leiden community detection algorithm [43] (Methods). The generated clusters were then annotated as immune cell subsets (B cell, CD4+ T cell, CD8+ T cell and plasma cell), monocyte/macrophages, erythroid, myeloid, and other CD4+ cells and others. To better demonstrate the modifications of DeepSNF denoising, we then utilized a weighted KNN approach to map the DeepSNF data into the DIMR-based clusters (Methods). The weights were acquired by calculating the Jaccard index between each DeepSNF-processed cell profile with all the DIMR cells (which is identical to the Jaccard graph construction of DIMR). For visualization, the cell markers of DIMR and DeepSNF were also compressed into two dimensions by the fast interpolation-based *t*-SNE algorithm [44] as Supplementary Fig. 33 (Methods). The assigned phenotypes of DIMR and DeepSNF datasets are demonstrated in Fig. 3a and the relative changes of each cell sub-population after DeepSNF processing is shown in Fig. 3b. After DeepSNF processing, B cells, CD8 T cells, plasma cells, CD4 T cells and other CD4+ cells decrease (8.78%, 1.23%, 10.11%, 15.52% and 11.51%, respectively), monocytes/macrophages and others increase (1.5% and 3.49%, respectively), while erythroid and myeloid cells remain almost not changed.

**Figure 3.**
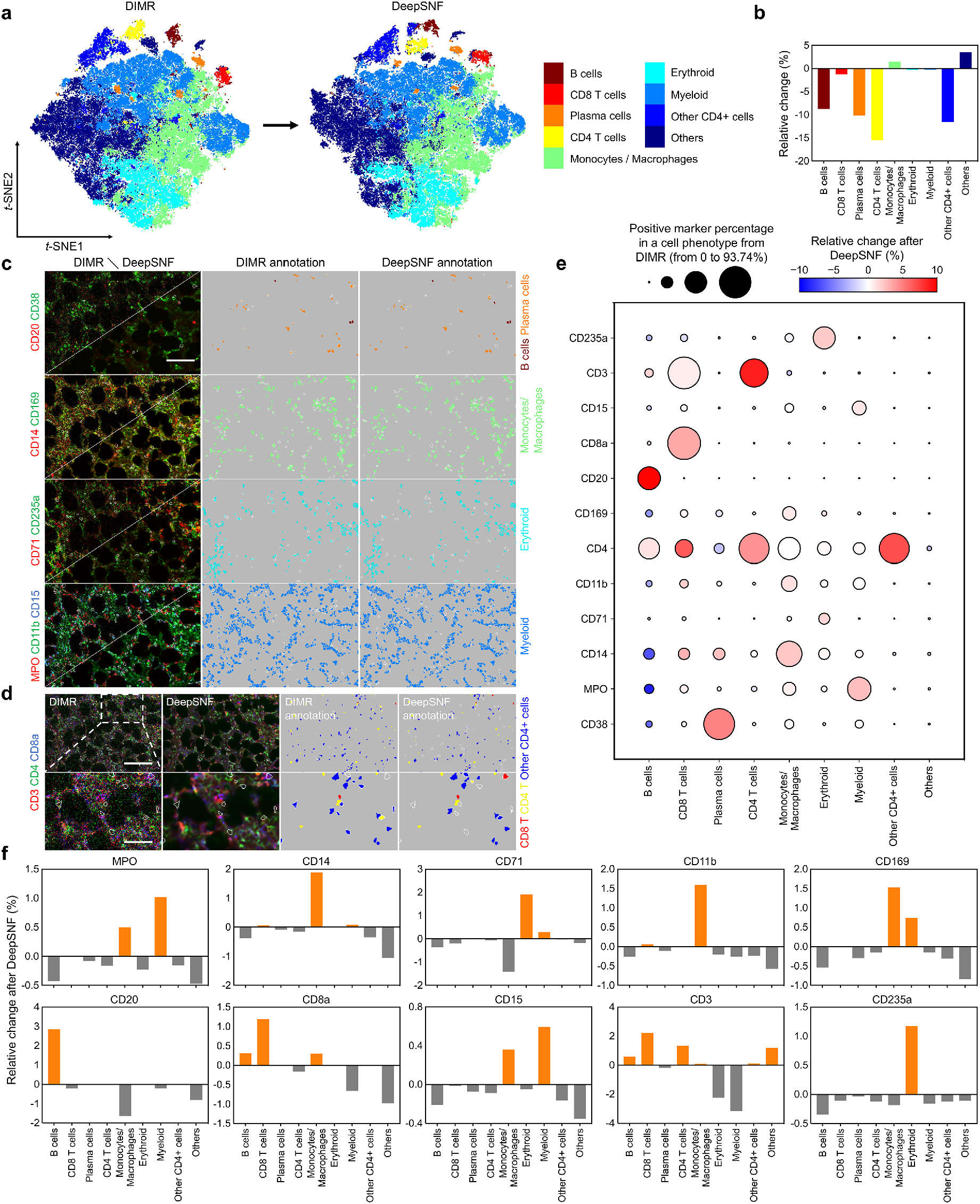
DeepSNF enhances automated cell phenotyping on human bone marrow IMC data. (a) *t*-SNE plots of DIMR and DeepSNF with cell phenotyping results. (b) The relative change in cell phenotypes before and after DeepSNF. (c), (d) Comparisons of DIMR and DeepSNF-processed IMC images labeled with different cell markers, and the corresponding cell annotation results. In (d), the bottom row corresponds to the white dashed box region selection in the first panel. In (c) and (d), the white contours represent the differential phenotyping results between DIMR and DeepSNF. (e) DeepSNF enhances the sensitivity of cell phenotyping. After DeepSNF processing, the non-specific marker signals reduce while the specific ones enrich in the cell types, respectively. The circle size indicates the positive marker percentage in a particular phenotype of DIMR, and the circle colour indicates the relative changes of the positive rate for the particular markers after DeepSNF enhancement. (f) DeepSNF enhances the specificity of cell phenotyping. With DeepSNF denoising, the ratios of specific phenotypes increase while those of non-specific phenotypes decrease in the positive markers. The relative change is the difference in percentage composition of each cell type before and after DeepSNF enhancement. Scale bar: (c) 110 *µ*m. (d) Top: 145 *µ*m, bottom: 50 *µ*m.

The phenotyping results of DIMR and DeepSNF were mapped back into their segmentation masks and images (Fig. 3c,d and Supplementary Fig. 34). To highlight cells where DeepSNF changes the cell phenotyping results, conflicting annotations between DIMR and DeepSNF were labeled with white contours, and the changes were quantified for cell phenotype and marker enrichments (Methods). After DeepSNF denoising, non-specific markers are reduced, while specific markers are enriched within the cell phenotypes (Fig. 3e). For example, we observed the positive rate increased for CD20 in B cells (7.91%), CD8a in CD8 T cells (2.71%), CD3 and CD4 in CD4 T cells (7% and 3.42%), MPO in myeloid cells (2.06%), CD38 in plasma cells (3.89%) and CD4 in other CD4+ cells (5.51%). Conversely, we observed a decrease of non-specific markers, such as CD14 and MPO in B cells (5.38% and 7.02%), CD3 in erythroid (1.56%) and myeloid (1.18%) cells, and all marker signals in “other” cells. Furthermore, the identified cell types were enriched in a marker-specific manner after DeepSNF (Fig. 3f). For instance, we observed a post-DeepSNF enrichment of monocytes/macrophages in CD14+ cells (1.89%) and CD11b+ cells (1.59%), and enrichment of B cells in CD20+ cells (2.86%). Similarly, myeloid cells were enriched in MPO+ (1.02%), and erythroid cells in CD71+, CD235a+ cells (1.91% and 1.18%). DeepSNF also yielded an enriched composition of CD8 and CD4 T cells (2.23% and 1.35%), and reduced composition of myeloid and erythroid cells (2.24% and 3.15%) in CD3+ cells. However, we noticed the enrichment of erythroid cells in CD169+ cells (0.74%), which may result from an artifact of the current segmentation approach due to the close relationship and irregular morphology at the boundaries between erythroids and macrophages within the bone marrow [45]. Cell phenotyping by immunostaining of FFPE tissues is also inherently limited by antibody specificity and antigen retrieval protocols. In this tissue, CD38+ and CD14+ antibody staining is not strictly restricted to single lineages, and these markers can be aberrantly expressed in myeloid neoplasms included in this data set (Supplementary Fig. 35). On manual inspection, DeepSNF improves the ability to identify co-localization of cell surface markers (Fig. 3c,d and Supplementary Fig. 34). Overall, DeepSNF enhances the sensitivity and specificity of cell phenotyping.

We observed that the enhancements in cell phenotyping and marker enrichments in Fig. 3e,f are related to the noise level of the IMC images. Specifically, DeepSNF has the highest impact on CD20 and CD3 related phenotypes, improvement for CD15, MPO and CD235a related phenotypes is limited, with moderate changes for other cell classes. These findings agree with Supplementary Fig. 29b, where we plot the STD of the normalized positive marker differences between DIRM and DeepSNF against intensity. To investigate the influence of DeepSNF on phenotyping results more deeply, we applied a leave-one-out DeepSNF strategy for CD20, CD3, CD71, CD235a and MPO (Methods, Supplementary Figs. 36–40). Briefly, by only processing one marker for hot pixel removal with DIMR, e.g. CD20, and all the other markers by both DIMR and DeepSNF. Then the same weighted KNN approach was applied on these leave-one-out DeepSNF datasets. Similar to the conclusions from Fig. 3e,f and Supplementary Fig. 29b, CD20 and CD3 denoised by DeepSNF could improve cell phenotyping remarkably because of the high noise level of the corresponding IMC images. DeepSNF has moderate impacts on CD71 due to better IMC image quality than those of CD20 and CD3, and has minor impact on MPO and CD235a because of their good IMC image qualities.

## 3 DISCUSSION

With the rise of novel multiplexed technologies for the characterization of cellular context in health and disease, IMC has surfaced as a valuable tool to investigate immunophenotypes while preserving spatial information. Differing from traditional multiplexed imaging approaches based upon fluorescence microscopy, IMC allows for simultaneous acquisition of more than 40 cell markers with greatly suppressed channel crosstalk, and avoids tissue and marker degradation in multi-round staining protocols. Furthermore, it eliminates autofluorescence and background signal issues that are inherent in fluorescent microscopy. The high-dimensional datasets then enable complex microenvironment analysis. However, IMC suffers from unique hot pixel and shot noise features. Analyzing raw IMC data without further restoration may lead to distortions, even errors, in downstream analysis. Contemporary denoising strategies are usually not adaptive or effective for these particular noise conditions. For example, the parameters of some methods need to be tuned manually, which are not suitable for large datasets and may cause subjective, batch, and channel-specific errors.

In this work, we propose IMC-Denoise to account for the specific technical noise present in IMC images. In this pipeline, the DIMR algorithm is first applied to adaptively remove hot pixels. It does not use a pre-set threshold or a particular range to define hot pixels, eliminating the impact of the density and intensity variations of hot pixels in different datasets or markers. Instead, it builds a histogram from the differential maps of raw images followed by an iterative outlier detection algorithm. In comparison with other methods, it achieves more robust hot pixel detection capability and normal pixel preserving performance. After hot pixel removal, the DeepSNF algorithm is proposed to restore image quality. I-divergence is derived as the optimal loss function for this denoising task. Due to the difficulty to acquire noise-free IMC images and repeated scanning for training labels, we applied the masking strategy with stratified sampling from Noise2Void to enable self-supervised training for this denoising task, in which multiple pixels are randomly masked and replaced by its adjacent pixels. With the continuity of biomarker signals in IMC, Hessian regularization is added in the loss function to boost the denoising performance. In DeepSNF, we train a single network for a single marker, which reduces the memory allocated for training. Nevertheless, we note that DeepSNF also works on multiple markers training (Supplementary Fig. 41). The trained network can be employed to other datasets which share the similar features as well (Supplementary Fig. 21). To determine the applicability of our approach, reference denoising algorithms were utilized to rigorously evaluate IMC-Denoise on multiple datasets and simulated data of a wide range of realizations. Compared to other methodologies, both DIMR and DeepSNF achieve the best denoising performance, qualitatively and quantitatively. Orthogonal approaches that have not been previously tested in evaluation of IMC restoration are also used to verify the image quality improvement by IMC-Denoise. Moreover, spillover can be corrected after hot pixel removal and shot noise filtering if needed, as indicated in Eq. (1). MIBI shares several features with IMC with respect to image formation and noise sources. Therefore, the denoising pipeline deployed here can also enhance MIBI datasets.

IMC-Denoise is effective at removing background noise and enhancing downstream analysis of IMC data with limited, subjective, user-input. Multiple datasets processed by DIMR and DeepSNF were tested with the existing IMC background removal methods, including single threshold binarization, semi-automated Ilastik-based, and MAUI, using the F1 score as the accuracy metric to evaluate the results. The qualitative and quantitative results indicate DeepSNF can affect significant background noise removal, and is superior than tedious semi-automated approaches. In particular, DeepSNF could unmix the IMC signal from background noise. Therefore, even the thesholding approach for background removal is not essential after DeepSNF denoising. Conventional workflows use manual gating strategies combined with prior cell marker knowledge to identify and compare cell types in pathological samples. We used real world data and these methods to evaluate the IMC denoising algorithm, and in comparison to DIMR, DIMR_Ilastik and DeepSNF, for single cell analyses. Automated IMC-Denoise performs equally or superior to the semi-manual Ilastik-based method on downstream single cell analysis. DeepSNF in particular enhances the cell clustering and annotation. Quantitative evaluations of cell phenotyping results indicate the improvement of sensitivity and specificity by DeepSNF denoising.

As noted, DeepSNF enhances all the markers and their downstream analysis. However, the marker channels with high noise levels benefit to a larger degree. Because of the signal-dependent characteristics of shot noise, the noise components of high SNR channels contribute less to overall image quality, and thus have lower impact on downstream analysis. While we find that further denoising by DeepSNF can be omitted when the mean expressions of positive markers are larger than 7 (MPO, CD15 and CD235a), predicated on our empirical findings, denoising all marker channels improves performance and is not computationally intensive. Limitations of IMC-Denoise include the inability to remove large hot pixel clusters, as DIMR cannot discriminate these larger areas of outliers from signal (Supplementary Fig. 42); and the self-supervised DeepSNF algorithm cannot reach the accuracy of supervised denoising methods due to unavailability of ground truths (Supplementary Figs. 7 and 8). Nevertheless, DIMR can remove single hot pixels and small hot clusters with several consecutive pixels, and DeepSNF performs better than other unsupervised and self-supervised denoising methods on IMC datasets. To conclude, we have developed the content aware IMC-Denoise for improved IMC image quality. Predicated on novel combination of differential map-based and self-supervised CNN-based algorithms, this pipeline removed hot pixels and effectively suppressed shot noise in IMC data. Multiple image and cell-based analyses on different IMC datasets verified the enhancements brought by this approach. We expect IMC-Denoise to become a widely used pipeline in IMC analysis due to its adaptability, effectiveness and flexibility.

## 4 METHODS

### 4.1 Human bone marrow dataset

Bone marrow sections were obtained from pathology archived formalin-fixed paraffin-embedded (FFPE) blocks from patients with normal morphology, myelodysplastic syndromes, or acute leukemia. Use of specimens for secondary analysis in this study was approved by the Washington University in St. Louis Institutional Review Board (#201912110).

### 4.2 Tissue staining and IMC data acquisition

Descriptions of cell markers and isotope tags are provided in Supplementary Tables 2–5. Staining was performed according to Fluidigm IMC recommendations for FFPE as follows. Briefly, tissue sections were dewaxed in xylene and rehydrated in a graded series of alcohol. Epitope retrieval was conducted in a water bath at 96 °C in Tris-EDTA buffer at pH 9 for 30 minutes, then cooled and washed in metal-free PBS. Blocking with Superblock (ThermoFisher) plus 5% FcX TruBlock (Biolegend) was followed by staining with antibody cocktail prepared in 0.5% BSA and metal-free PBS overnight at 4 °C. Sections were washed in 0.02% TritonX100 followed by metal-free PBS, then nuclear staining was performed using 1:200 or 1:300 dilution of Intercalator-Ir (125 *µ*M, Fluidigm) solution for 30 minutes, followed by ddH_2_O for 5 minutes. Slides were air-dried before IMC measurement.

The abundance of bound antibody was quantified using the Hyperion imaging system (Fluidigm) controlled by CyTOF Software (version 7.0.8493), with UV-laser set at 200 Hz. Count data were then converted to tiff image stacks for further analysis using MCD Viewer (version 1.0.560.6, Fluidigm) or imctools (Bodenmiller lab, https://github.com/BodenmillerGroup/imctools).

### 4.3 Tissue staining and IF data acquisition

For IF staining, tissue was prepared as described above, then stained overnight at 4 °C with a single metal-conjugated primary antibody (CD3, CD4, CD169 or CD61 in Supplemental Table 5). The single-stained tissue was washed, then stained with secondary antibody (donkey anti-rabbit AF647 or goat anti-mouse AF750, Invitrogen, 2mg/mL diluted 1:400 in 0.5% BSA in PBS) at room temperature 1 hour, followed by additional washing in PBS and DAPI (1ug/mL) staining. Slides were mounted with SlowFade Glass antifade reagent (ThermoFisher) and # 1 1/2 coverslips. Images were acquired using Leica DMi8 inverted widefield microscope with Lumencor SOLA SE U-nIR light engine, DAPI/FITC/TRITC/Cy5/Cy7 filters, DFC9000 GT sCMOS camera, PL APO 20x/0.80 objective and LAS X software (version 3.7.3.23245). After image acquisition, coverslips were removed with gentle agitation in PBS, then Ir-intercalator staining, washing and drying performed as above for subsequent Hyperion data acquisition.

### 4.4 Human pancreatic, breast cancer IMC datasets and MIBI dataset

We applied the human pancreatic [10], breast cancer [12] IMC datasets and a MIBI dataset [19] to verify the flexibility of IMC-Denoise. All of these datasets are publicly available. The links are provided in the corresponding papers. In breast cancer dataset, CD3, CD20, CD45, CD68, c-Myc, EGFR, EpCAM, Ki-67, Rabbit IgG H L, Slug, Twist and vWF were selected; in pancreatic cancer dataset, CD3, CD4, CD8, CD11b, CD14, CD31, CD44, CD45, CD45RO, CD56, Foxp3 and pS6 were selected; and in the MIBI dataset, CD3, CD4, CD8, CD11b, CD11c, CD14, CD20, CD31, CD45, CD68, CD206 and HLA-DR were selected. The two IMC datasets were processed by both DIMR and DeepSNF. The MIBI dataset is only processed by DeepSNF because no hot pixels are observed, and the hot clusters observed in MIBI images can be removed by the MAUI software package [19]. Details on software implementation can be found in the relevant sections below.

### 4.5 Neural network implementation

The neural network follows the U-Net architecture [35] with Res-block modules [36], in which the input and output images share the same size (Supplementary Fig. 11). Starting with the input, the encoder path gradually condenses the spatial information into high-level feature maps with growing depths; the decoder path reverses this process by recombining the information into feature maps with gradually increased lateral details. The information in adjacent feature maps transfers by convolving with 3 × 3 convolutional filters. The down-sampling and up-sampling are done by 2 × 2 max-pooling and 2 × 2 up-sampling operations, respectively. Res-blocks are applied to facilitate efficient training. Each res-block contains convolution layer, batch normalization and the rectified linear unit nonlinear activation. Drop out layers are also added with 0.5 dropout rate after the central two res-blocks to mitigate overfitting. Skip connections tunnel the high-frequency information from shallower layers to deeper layers with the same spatial scales. We use the softplus function as the activation function of the final layer and Eq. (2) as the loss function.

The hot pixel-removed images are split into multiple 64 × 64 patches. Then, the patches are rotated by 90°, 180° and 270°, and flipped as a data augmentation approach. In IMC images, foreground objects of interest might be distributed sparsely. In this case, the model might overfit the background areas and fail to learn the structure of foreground objects if the entire image is used indiscriminately for training. Therefore, patches from the background regions are excluded from training. In IMC images, pixels with intensity value 0 are considered as background. Afterwards, we define the background pixel ratio *r* as ratio of the number of background pixels and that of total pixels in the patch. Patches are considered as the background regions if *r* ≤ *ρ*, where *ρ* is the threshold and set from 0.2 to 0.99 for different channels and datasets. We applied a smaller *ρ* for the datasets less sparse images and vice versa. For good generalization ability of the network, we recommend at least 5000 patches for training. Before training, all the generated patches were percentile normalized (99.9 to 99.999, Supplementary Note 1.3.3). The percentile of 99.9 was applied for those training sets with extremely bright markers and larger percentile with relatively homogeneous intensity distributions. To balance the training efficiency and accuracy, 0.2% pixels of each patch are masked and replaced by their neighbours using a stratified sampling strategy [24]. Finally, 85% of the patches are set as training set and the rest as validation set.

All models were trained using Keras [46] (version 2.3.1) on a single NVIDIA Quadro RTX 6000 GPU with 24 GB of VRAM. Adam optimizer [47] was applied as the optimization algorithm with a initial learning rate of 0.001 for 200 epochs and batch size of 128. Learning rate is multiplied by 0.6 if validation loss does not improve for 20 epoches. The training details for all the datasets are summarized as Supplementary Tables 6–11. Note that the training datasets for N2V, MN2V and DeepSNF-NR are the same as those for DeepSNF, and the training time of N2V, MN2V and DeepSNF-NR is approximately equal to that of DeepSNF.

### 4.6 Neural network inference details

Given a trained denoising model, we denoise full-size IMC images to avoid edge stitching effects. In order to achieve end-to-end prediction, we pad pixels around each image so their width and height are the multiples of 16 with reference to the network architecture (Supplementary Fig. 11). The padding pixels are the replications of the border pixels. Before prediction, the IMC images are normalized by the pre-calculated maximum of the corresponding channels in the training set. The outputs of the network are re-scaled and set as the denoised images. Given the trained denoising model, inference is fast. We are able to denoise IMC images with pixels of 1000 by 1000 less than 1 second per image on a single NVIDIA Quadro RTX 6000 GPU.

### 4.7 Semi-automated Ilastik-based background noise removal

The semi-automated strategy in [21] utilizes Ilastik segmentation [48] to remove background noise in IMC images. An expert annotates signal or background regions of IMC images, and then Ilastik trains a random forest classifier for background noise removal. To achieve good denoising quality, large areas of background require manual labeling, which is labor-intensive. Furthermore, low image quality may affect the accuracy of this method as well. After background removal, the images are binarized to solve batch effect issues. Then the single cell information is calculated by counting the positive signal frequency rather than the mean intensity of every single cell. Here, we only utilized Ilastik (version 1.3.2post1) for background noise removal of IMC images, and still applied the mean intensities as the single cell profiles. To better reveal the enhancement by DeepSNF, we applied the same labels for the trainings of DIMR and DeepSNF-processed images.

### 4.8 MAUI

MAUI software package [7, 19] includes spillover correction, noise removal and aggregate removal. All three steps require expert observation, which is also labor-intensive. Here, we only benchmarked the noise removal method in MAUI with our DeepSNF algorithm. Briefly, it calculates the distances between a non-zero pixel and its K nearest non-zero neighbours, then builds a histogram based on the summations of the distances for all the non-zero pixels. Next, a threshold is manually selected to remove the pixels with larger summations by observing the distribution of the histogram. This method is based on the assumption that noisy regions look more sparse than normal regions. MAUI was implemented by the software package from https://github.com/angelolab/MAUI. The parameter K and the threshold were manually tuned to guarantee the best performance of MAUI (Supplementary Fig. 27).

### 4.9 Pixel classification and cell segmentation

In single cell segmentation, the pixels in each image were defined as belonging to the nucleus, cytoplasm, or background compartment using the pixel classification module of Ilastik [48] (version 1.3.2post1) as described in https://github.com/BodenmillerGroup/ImcSegmentationPipeline. The Random Forest classifier was trained on the channels including CD38, MPO, CD14, CD71, CD11b, CD4, CD20, CD8a, CD15, Ki-67, CD3, CD45RO CD235a, Histone-H3 and Iridium. This allowed for the generation of maps that integrate for each pixel the probability of belonging to each of three compartments. All the images were denoised by IMC-Denoise before segmentation. Based on the trained classifier, probability maps were generated for the whole dataset and exported as tiff files in batch mode.

Subsequently, CellProfiler [49] (version 3.1.8) was used to define cell masks for marker expression quantification. To define cell borders, nuclei were first identified as primary objects based on ilastik probability maps and expanded through the cytoplasm compartment until either a neighboring cell or the background compartment was reached. Cell masks were generated for identification of single cells and used to extract single-cell information from IMC images.

### 4.10 Single-cell marker profile extraction

We used HistoCAT [50] (version 1.7.6) to extract single-cell marker profiles based on the IMC images and their segmentation masks. All the data were not transformed and used directly.

### 4.11 Positive cell identification

To obtain an easy way to identify the positive cells for each marker, we modified the method in [14]. Briefly, univariate Gaussian mixture models with scikit-learn [51] (version 1.0.2) were used to estimate the positive thresholds of each marker. Before threshold estimation, both DIMR and DeepSNF data was 99th-percentile normalized so that the impact of extremely bright cells can be eliminated. Then Z-score normalization was applied because we observed the data scales might shift slightly after DeepSNF (Supplementary Fig. 29a).

For each channel, we performed model selection with models with 6 to 15 mixtures for DIMR data, in order to estimate the positive threshold accurately. We selected the model on the basis of the Davies-Bouldin index [52] and identified a positive threshold for a given channel by considering both the distributions of cell profiles and the overall IMC image intensities. After choosing the positive thresholds for DIMR markers, we applied the same trained models on DeepSNF data and identified the thresholds with the same standard. The estimated thresholds for positive markers of DIMR and DeepSNF are summarized in Supplementary Table 12.

### 4.12 Cell-type annotation

A subset of markers from the human bone marrow dataset was utilized for cell phenotypic annotation, including CD38, MPO, CD14, CD71, CD11b, CD4, CD169, CD20, CD8a, CD15, CD3 and CD235a. Before analysis, data were 99th-percentile normalized followed by Z-score normalization. Then the DIMR data was clustered by searching 20 nearest neighbours of each cell and then constructing a Jaccard graph [42], followed by the Leiden community detection algorithm [43] with resolution of 6.0, which resulted in over clustering with 117 clusters. The generated clusters were manually labelled with a broad ontogeny and the channels that were most abundant in each cluster (Supplementary Fig. 33), resulting in 9 cell types, including immune cell subsets (B cell, CD4+ T cell, CD8+ T cell and plasma cell), monocyte/macrophages, erythroid, myeloid, and other CD4+ cells and others.

The DeepSNF data clustering and annotation utilized a weighted K-nearest neighbour (KNN) approach (K=20) to map the DeepSNF data into the DIMR clusters. It first constructs a Jaccard graph between each cell from DeepSNF and all the DIMR cells, and then maps the DeepSNF data into the DIMR clusters with the shortest weighted distance. The leave-one-out DeepSNF data was also annotated with the same approach. The Jaccard graph construction and Leiden algorithm were implemented by the software packages from https://github.com/jacoblevine/PhenoGraph and https://github.com/vtraag/leidenalg, respectively. DeepSNF data annotation was implemented with customized python scripts.

Notably, multiple strategies were applied to reduce the noise impact during DIMR clustering: (1) Z-score normalization is consistent for handling different sources of noise in multiplexed cell data, including low intensity signal, high background signal, segmentation noise, and imaging artifacts, as verified by [53]; (2) the Jaccard graph was constructed because of its robustness to noise, which is verified in [42]; and (3) over-clustering could improve the clustering accuracy [53]. Besides, we didn’t annotate the DeepSNF and leave-one-out DeepSNF data with the same approach of DIMR because (1) The community detection results by Leiden algorithm is random so that it is very difficult to compare the annotations from different data; and (2) the weighted KNN method for DeepSNF and leave-one-out DeepSNF clustering could clearly reveal the differences before and after the processing.

### 4.13 Fast interpolation-based *t*-SNE algorithm

For visualization, high-dimensional single-cell data of DIMR and DeepSNF were reduced to two dimensions using the nonlinear dimensionality reduction algorithm fast interpolation-based *t*-SNE [44]. This algorithm was implemented by the software package in https://github.com/KlugerLab/FIt-SNE. Before the analysis, data were 99th-percentile normalized followed by Z-score normalization. The *t*-SNE parameters with perplexity of 50 and theta of 0.5 were used. The random seeds for the individual runs were recorded.

### 4.14 Enrichment calculation of positive cell markers after DeepSNF and leave-one-out DeepSNF

To evaluate the effect of positive cell marker enrichment after DeepSNF and leave-one-out DeepSNF, the cell-type annotations before and after the processing were selected, and the percentage of positive markers on each cell types was calculated. The relative change was then defined as the difference between the percentage of positive markers after and before the processing.

### 4.15 Enrichment calculation of cell types after DeepSNF and leave-one-out DeepSNF

To evaluate the effect of cell type enrichment after DeepSNF and leave-one-out DeepSNF, the positive cells for a given marker before and after the processing were selected, and the percentage of each cell type based on cell-type annotation was calculated. The relative change was then defined as the difference between the percentage of cell-type composition after and before the processing.

### 4.16 Accuracy metrics

In simulation, the accuracy metrics including root mean squared error, peak SNR and structural similarity [41] are used to access the image qualities because of the availability of ground truths. They are defined in Supplementary Note 3.2 in detail.

For the real experimental data, five types of metrics were used for quantitative evaluations. The standard deviation of background (STDB) and contrast-to-noise ratio (CNR) were used to evaluate the noise level and contrast of IMC images. CNR is defined as Eq. (3),

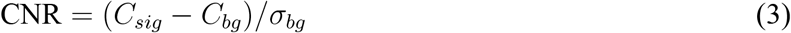

where *C*_*sig*_ and *C*_*bg*_ are the mean of the signal and background and *σ*_*bg*_ is the STDB. In this metric, the signal and background regions of IMC images are manually annotated.

Pearson’s correlation coefficient (PCC) was used as the spatial metric to reflect the similarity between images. Alternatively, Spearman’s correlation coefficient (SCC) was used as the agreement between the marker expressions in single cell scale. The PCC between image *Y* and the reference *Y*_*ref*_ is defined as Eq. (4),

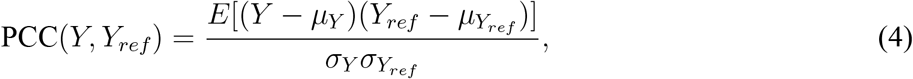

where *µ*_*Y*_ and 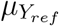 are the mean values of images *Y* and *Y*_*ref*_, respectively; *σ*_*Y*_ and 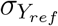 are the standard deviations of *Y* and *Y*_*ref*_, respectively; and *E* represents arithmetic mean.

Furthermore, F1 score was used to evaluate the accuracy of background noise removal, which can be formulated as Eq. (5).

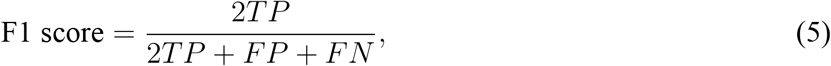

where *TP, FP* and *FN* are the pixel number of true positives, false positives and false negatives, respectively. All of the evaluation process was implemented with customized MATLAB (R2021a, MathWork) scripts. RMSE, PSNR, SSIM, PCC, SCC and F1 score were computed using MATLAB built-in functions.

### 4.17 Statistical analysis

Quantitative data are presented as box-and-whisker plots (center line, median; limits, 75% and 25%; whiskers, maximum and minimum). The paired two-side Student’s t-test was used for the statistical significance determination. All the statistical tests are implemented with Prism 9 (GraphPad Software Inc.). Statistical significance at P<0.05, 0.01, 0.001 and 0.0001 are denoted by *, **, *** and ****, respectively. “ns” means “no significance”.

## Data availability

The human bone marrow IMC data and simulated data are available from Zenodo https://doi.org/10.5281/zenodo.6533905. The human pancreatic cancer IMC dataset [10], the human breast cancer IMC dataset [12], and the MIBI data set [19] can be downloaded through the links provided by the authors in their papers, respectively. All the other data supporting the results in this paper can be accessed from https://doi.org/10.5281/zenodo.6516116.

## Code availability

The code used in this study and the corresponding tutorial are publicly available at https://github.com/PENGLU-WashU/IMC_Denoise.

## Acknowledgement

Research reported in this publication was supported in part by NIH NCI R01CA240711, R01CA229893, K12 CA167540 and 1P50CA171063; NHLBI R21HL150636; American Society of Hematology Scholar Award; and Evans Foundation Edward P. Evans Center for MDS. We thank the Alvin J. Siteman Cancer Center at Washington University School of Medicine and Barnes-Jewish Hospital in St. Louis, MO., for the use of the Bursky Center for Human Immunology and Immunotherapy Programs Immunomonitoring Laboratory, which provided IMC service. The Siteman Cancer Center is supported in part by NCI Cancer Center Support Grant #P30 CA091842. The content is solely the responsibility of the authors and does not necessarily represent the official views of the NIH.

## Author contributions

P.L., K.A.O. and D.L.J.T. conceived and designed the project. P.L. developed and implemented the software. K.A.O., D.E.B., S.T.O., M.B.R., D.A.C.F. and D.C.L. obtained the human bone marrow tissues and performed the IMC staining and imaging. P.L., K.A.O. and D.L.J.T. conducted image and downstream analysis. R.K.P., W.N.B., K.G.S., S.T.O. and D.C.L. helped with analysis. D.L.J.T. supervised the project. P.L., K.A.O. and D.L.J.T. wrote the manuscript with input from all authors.

## Competing interests

The authors declares no competing interests.

## Notes

### Competing Interest Statement

The authors have declared no competing interest.

## References

[1] Y. Saeys, S. Van Gassen, B. N. Lambrecht, Computational flow cytometry: helping to make sense of high-dimensional immunology data, Nature Reviews Immunology 16 (7) (2016) 449–462.

[2] A. Adan, G. Alizada, Y. Kiraz, Y. Baran, A. Nalbant, Flow cytometry: basic principles and applications, Critical reviews in biotechnology 37 (2) (2017) 163–176.

[3] J. W. Hickey, E. K. Neumann, A. J. Radtke, J. M. Camarillo, R. T. Beuschel, A. Albanese, E. McDonough, J. Hatler, A. E. Wiblin, J. Fisher, et al., Spatial mapping of protein composition and tissue organization: a primer for multiplexed antibody-based imaging, Nature methods 19 (3) (2022) 284–295.

[4] J.-R. Lin, B. Izar, S. Wang, C. Yapp, S. Mei, P. M. Shah, S. Santagata, P. K. Sorger, Highly multiplexed immunofluorescence imaging of human tissues and tumors using t-cycif and conventional optical microscopes, Elife 7 (2018).

[5] Y. Goltsev, N. Samusik, J. Kennedy-Darling, S. Bhate, M. Hale, G. Vazquez, S. Black, G. P. Nolan, Deep profiling of mouse splenic architecture with codex multiplexed imaging, Cell 174 (4) (2018) 968–981.

[6] M. Angelo, S. C. Bendall, R. Finck, M. B. Hale, C. Hitzman, A. D. Borowsky, R. M. Levenson, J. B. Lowe, S. D. Liu, S. Zhao, et al., Multiplexed ion beam imaging of human breast tumors, Nature medicine 20 (4) (2014) 436–442.

[7] L. Keren, M. Bosse, D. Marquez, R. Angoshtari, S. Jain, S. Varma, S.-R. Yang, A. Kurian, D. Van Valen, R. West, et al., A structured tumor-immune microenvironment in triple negative breast cancer revealed by multiplexed ion beam imaging, Cell 174 (6) (2018) 1373–1387.

[8] C. Giesen, H. A. Wang, D. Schapiro, N. Zivanovic, A. Jacobs, B. Hattendorf, P. J. Schüffler, D. Grolimund, J. M. Buhmann, S. Brandt, et al., Highly multiplexed imaging of tumor tissues with subcellular resolution by mass cytometry, Nature methods 11 (4) (2014) 417–422.

[9] H. Baharlou, N. P. Canete, A. L. Cunningham, A. N. Harman, E. Patrick, Mass cytometry imaging for the study of human diseases—applications and data analysis strategies, Frontiers in immunology 10 (2019) 2657.

[10] Y. J. Wang, D. Traum, J. Schug, L. Gao, C. Liu, M. A. Atkinson, A. C. Powers, M. D. Feldman, A. Naji, K.-M. Chang, et al., Multiplexed in situ imaging mass cytometry analysis of the human endocrine pancreas and immune system in type 1 diabetes, Cell metabolism 29 (3) (2019) 769–783.

[11] N. Damond, S. Engler, V. R. Zanotelli, D. Schapiro, C. H. Wasserfall, I. Kusmartseva, H. S. Nick, F. Thorel, P. L. Herrera, M. A. Atkinson, et al., A map of human type 1 diabetes progression by imaging mass cytometry, Cell metabolism 29 (3) (2019) 755–768.

[12] H. W. Jackson, J. R. Fischer, V. R. Zanotelli, H. R. Ali, R. Mechera, S. D. Soysal, H. Moch, S. Muenst, Z. Varga, W. P. Weber, et al., The single-cell pathology landscape of breast cancer, Nature 578 (7796) (2020) 615–620.

[13] H. R. Ali, H. W. Jackson, V. R. Zanotelli, E. Danenberg, J. R. Fischer, H. Bardwell, E. Provenzano, O. M. Rueda, S.-F. Chin, S. Aparicio, et al., Imaging mass cytometry and multiplatform genomics define the phenogenomic landscape of breast cancer, Nature Cancer 1 (2) (2020) 163–175.

[14] A. F. Rendeiro, H. Ravichandran, Y. Bram, V. Chandar, J. Kim, C. Meydan, J. Park, J. Foox, T. Hether, S. Warren, et al., The spatial landscape of lung pathology during covid-19 progression, Nature (2021) 1–6.

[15] M. Wu, M. Y. Lee, V. Bahl, D. Traum, J. Schug, I. Kusmartseva, M. A. Atkinson, G. Fan, K. H. Kaestner, H. Consortium, et al., Single-cell analysis of the human pancreas in type 2 diabetes using multi-spectral imaging mass cytometry, Cell reports 37 (5) (2021) 109919.

[16] D. Moldoveanu, L. Ramsay, M. Lajoie, L. Anderson-Trocme, M. Lingrand, D. Berry, L. J. Perus, Y. Wei, C. Moraes, R. Alkallas, et al., Spatially mapping the immune landscape of melanoma using imaging mass cytometry, Science Immunology 7 (70) (2022) eabi5072.

[17] L. Kuett, R. Catena, A. Özcan, A. Plüss, P. Schraml, H. Moch, N. de Souza, B. Bodenmiller, Threedimensional imaging mass cytometry for highly multiplexed molecular and cellular mapping of tissues and the tumor microenvironment, Nature Cancer 3 (2022) 122–133.

[18] S. Chevrier, H. L. Crowell, V. R. Zanotelli, S. Engler, M. D. Robinson, B. Bodenmiller, Compensation of signal spillover in suspension and imaging mass cytometry, Cell Systems 6 (5) (2018) 612–620.

[19] A. Baranski, I. Milo, S. Greenbaum, J.-P. Oliveria, D. Mrdjen, M. Angelo, L. Keren, Maui (mbi analysis user interface)—an image processing pipeline for multiplexed mass based imaging, PLOS Computational Biology 17 (4) (2021) e1008887.

[20] V. Zanotelli, B. Bodenmiller, Imc segmentation pipeline: a pixel classification based multiplexed image segmentation pipeline, Zenodo https://doi.org/10.5281/zenodo3841960 (2017).

[21] M. E. Ijsselsteijn, A. Somarakis, B. P. Lelieveldt, T. Höllt, N. F. de Miranda, Semi-automated background removal limits data loss and normalizes imaging mass cytometry data, Cytometry Part A (2021).

[22] A. Buades, B. Coll, J.-M. Morel, A non-local algorithm for image denoising, in: 2005 IEEE Computer Society Conference on Computer Vision and Pattern Recognition (CVPR’05), Vol. 2, IEEE, 2005, pp. 60–65.

[23] K. Dabov, A. Foi, V. Katkovnik, K. Egiazarian, Image denoising by sparse 3-d transform-domain collaborative filtering, IEEE Transactions on image processing 16 (8) (2007) 2080–2095.

[24] A. Krull, T.-O. Buchholz, F. Jug, Noise2void-learning denoising from single noisy images, in: Proceedings of the IEEE/CVF Conference on Computer Vision and Pattern Recognition, 2019, pp. 2129–2137.

[25] F. J. Anscombe, The transformation of poisson, binomial and negative-binomial data, Biometrika 35 (3/4) (1948) 246–254.

[26] M. F. T. B. C. Russell, W. T. Freeman, Exploiting the sparse derivative prior for super-resolution and image demosaicing, in: Proceedings of the Third International Workshop Statistical and Computational Theories of Vision, 2003, pp. 1–28.

[27] B. W. Silverman, Density estimation for statistics and data analysis, Routledge, 2018.

[28] M. Makitalo, A. Foi, Optimal inversion of the anscombe transformation in low-count poisson image denoising, IEEE transactions on Image Processing 20 (1) (2010) 99–109.

[29] L. Finesso, P. Spreij, Nonnegative matrix factorization and i-divergence alternating minimization, Linear Algebra and its Applications 416 (2-3) (2006) 270–287.

[30] M. Weigert, U. Schmidt, T. Boothe, A. Müller, A. Dibrov, A. Jain, B. Wilhelm, D. Schmidt, C. Broaddus, S. Culley, et al., Content-aware image restoration: pushing the limits of fluorescence microscopy, Nature methods 15 (12) (2018) 1090–1097.

[31] Z. Wang, Y. Xie, S. Ji, Global voxel transformer networks for augmented microscopy, Nature Machine Intelligence 3 (2) (2021) 161–171.

[32] J. Chen, H. Sasaki, H. Lai, Y. Su, J. Liu, Y. Wu, A. Zhovmer, C. A. Combs, I. Rey-Suarez, H.-Y. Chang, et al., Three-dimensional residual channel attention networks denoise and sharpen fluorescence microscopy image volumes, Nature Methods (2021) 1–10.

[33] J. Lehtinen, J. Munkberg, J. Hasselgren, S. Laine, T. Karras, M. Aittala, T. Aila, Noise2noise: Learning image restoration without clean data, arXiv preprint arXiv:1803.04189 (2018).

[34] J. Batson, L. Royer, Noise2self: Blind denoising by self-supervision, in: International Conference on Machine Learning, PMLR, 2019, pp. 524–533.

[35] O. Ronneberger, P. Fischer, T. Brox, U-net: Convolutional networks for biomedical image segmentation, in: International Conference on Medical image computing and computer-assisted intervention, Springer, 2015, pp. 234–241.

[36] K. He, X. Zhang, S. Ren, J. Sun, Identity mappings in deep residual networks, in: European conference on computer vision, Springer, 2016, pp. 630–645.

[37] P. Lu, N. Benabdallah, W. Jiang, B. W. Simons, H. Zhang, R. F. Hobbs, D. Ulmert, B. Baumann, R. K. Pachynski, A. K. Jha, et al., Blind image restoration enhances digital autoradiographic imaging of radiopharmaceutical tissue distribution, Journal of Nuclear Medicine (2021).

[38] X. Huang, J. Fan, L. Li, H. Liu, R. Wu, Y. Wu, L. Wei, H. Mao, A. Lal, P. Xi, et al., Fast, longterm, super-resolution imaging with hessian structured illumination microscopy, Nature biotechnology 36 (5) (2018) 451–459.

[39] W. Zhao, S. Zhao, L. Li, X. Huang, S. Xing, Y. Zhang, G. Qiu, Z. Han, Y. Shang, D.-e. Sun, et al., Sparse deconvolution improves the resolution of live-cell super-resolution fluorescence microscopy, Nature biotechnology (2021) 1–12.

[40] R. Rashid, G. Gaglia, Y.-A. Chen, J.-R. Lin, Z. Du, Z. Maliga, D. Schapiro, C. Yapp, J. Muhlich, A. Sokolov, et al., Highly multiplexed immunofluorescence images and single-cell data of immune markers in tonsil and lung cancer, Scientific data 6 (1) (2019) 1–10.

[41] Z. Wang, A. C. Bovik, H. R. Sheikh, E. P. Simoncelli, Image quality assessment: from error visibility to structural similarity, IEEE transactions on image processing 13 (4) (2004) 600–612.

[42] J. H. Levine, E. F. Simonds, S. C. Bendall, K. L. Davis, D. A. El-ad, M. D. Tadmor, O. Litvin, H. G. Fienberg, A. Jager, E. R. Zunder, et al., Data-driven phenotypic dissection of aml reveals progenitor-like cells that correlate with prognosis, Cell 162 (1) (2015) 184–197.

[43] V. A. Traag, L. Waltman, N. J. Van Eck, From louvain to leiden: guaranteeing well-connected communities, Scientific reports 9 (1) (2019) 1–12.

[44] G. C. Linderman, M. Rachh, J. G. Hoskins, S. Steinerberger, Y. Kluger, Fast interpolation-based t-sne for improved visualization of single-cell rna-seq data, Nature methods 16 (3) (2019) 243–245.

[45] A. Chow, M. Huggins, J. Ahmed, D. Hashimoto, D. Lucas, Y. Kunisaki, S. Pinho, M. Leboeuf, C. Noizat, N. Van Rooijen, et al., Cd169+ macrophages provide a niche promoting erythropoiesis under homeostasis and stress, Nature medicine 19 (4) (2013) 429–436.

[46] F. Chollet, keras, https://github.com/fchollet/keras (2015).

[47] D. P. Kingma, J. Ba, Adam: A method for stochastic optimization, arXiv preprint arXiv:1412.6980 (2014).

[48] C. Sommer, C. Straehle, U. Koethe, F. A. Hamprecht, Ilastik: Interactive learning and segmentation toolkit, in: 2011 IEEE international symposium on biomedical imaging: From nano to macro, IEEE, 2011, pp. 230–233.

[49] L. Kamentsky, T. R. Jones, A. Fraser, M.-A. Bray, D. J. Logan, K. L. Madden, V. Ljosa, C. Rueden, K. W. Eliceiri, A. E. Carpenter, Improved structure, function and compatibility for cellprofiler: modular high-throughput image analysis software, Bioinformatics 27 (8) (2011) 1179–1180.

[50] D. Schapiro, H. W. Jackson, S. Raghuraman, J. R. Fischer, V. R. Zanotelli, D. Schulz, C. Giesen, R. Catena, Z. Varga, B. Bodenmiller, histocat: analysis of cell phenotypes and interactions in multiplex image cytometry data, Nature methods 14 (9) (2017) 873.

[51] F. Pedregosa, G. Varoquaux, A. Gramfort, V. Michel, B. Thirion, O. Grisel, M. Blondel, P. Prettenhofer, R. Weiss, V. Dubourg, et al., Scikit-learn: Machine learning in python, the Journal of machine Learning research 12 (2011) 2825–2830.

[52] D. L. Davies, D. W. Bouldin, A cluster separation measure, IEEE transactions on pattern analysis and machine intelligence (1979) 224–227.

[53] J. W. Hickey, Y. Tan, G. P. Nolan, Y. Goltsev, Strategies for accurate cell type identification in codex multiplexed imaging data, Frontiers in Immunology (2021) 3317.

