## Supplementary Materials for "IMC-Denoise: a content aware denoising pipeline to enhance Imaging Mass Cytometry"

**Supplementary Figures** **30**

**Supplementary Tables** **62**

**References** **70**

### Supplementary Note 1: IMC-Denoise framework

#### 1.1 Imaging mass cytometry imaging model

In the IMC imaging process, there are three major noise sources: hot pixels, ion shot noise, and spillover. Hot pixels are concentrated areas of high counts which are uncorrelated with any biological structures. In IMC images this artifact is most often observed as single hot pixels; however, small areas or clusters with several consecutive pixels may also be found. Ion shot noise exists because of the ion counting imaging process. The higher ion counts, the lower the shot noise level will be. Spillover, which is signal detected in one channel that is originating from an adjacent channel, can be neglected if the marker panel is well designed and properly titrated. Even with the existence of spillover, the contributions to total noise are very weak, which is approximately only one percent of the original intensity from the originating channel [1]. While applying low concentrations of staining antibodies minimizes spillover concerns, this will also result in even lower ion counts, and thus higher shot noise levels.

Here, we model the ion counting imaging as a Poisson process and hot pixels as outliers with much larger intensity than their adjacent pixels. As a result, the IMC imaging model is built as Eq. (1).

$$R = \mathcal{P}[X + X^{\text{spillover}}] + Q, \tag{1}$$

where $R = \{r_p\}$ is the raw image set, $X = \{x_p\}$ the true signals without noise, $X^{\text{spillover}} = \{x_p^{\text{spillover}}\}$ the spillover signals without noise, $Q = \{q_p\}$ the hot pixels, $p$ the pixel index and $\mathcal{P}[x]$ the Poisson noise with mean $x$.

In this paper, we only consider the hot pixels and ion shot noise in raw IMC images. If spillover is observed, it should be corrected after the restoration of these two noise sources, as its signal is contaminated by them as well. Therefore, the IMC imaging model is simplified as Eq. (2).

$$R = \mathcal{P}[X] + Q, \tag{2}$$

#### 1.2 Differential intensity map-based restoration algorithm for hot pixel removal (DIMR)

##### 1.2.1 Hot pixel unmixing

The Poisson model can be feasibly estimated as a Gaussian process [2] so that Eq. (2) can be converted as

$$R = \mathcal{N}(X, X) + Q. \tag{3}$$

From Eq. (3), larger true signal $X$ will result in larger variance. Thus, the contaminated signal $\mathcal{P}[X]$ with larger $X$ is more likely to be considered as hot pixels $Q$ and vice versa. To avoid such false detection, we stabilize the variance of the signal $X$ with the Anscombe transformation [3] as Eq. (4).

$$R' = X' + \mathcal{N}(0, 1) + Q', \tag{4}$$

where $\mathcal{N}(0, 1)$ is the additive noise with standard Gaussian distribution, and $R'$, $X'$ and $Q'$ are the transformed raw image, "clean" signal and hot pixels, respectively. As hot pixels are local maxima in IMC images, we detect them by comparing adjacent pixels in a 3×3 sliding window. Considering the nonlinearity of the Anscombe transformation [4], pixels with intensities lower than 4 in $R$ are omitted directly in order to exclude the impact of very low intensity regions, which cannot be outliers. Additionally, the difference between adjacent pixels can be fitted as a generalized Gaussian distribution [5]. Thus, in the 3×3 sliding window, we derive Eq. (5) by calculating the differences between the center pixel and its 8 neighbours. In Eq. (5), $i$ is the neighbour index in the sliding window ($i \in \{1, 2, ..., 8\}$), and $\mathcal{G}(\mu, \alpha, \beta)$ is a generalized Gaussian distribution with location $\mu$, scale $\alpha$ and shape $\beta$. Without the hot pixel component $Q' - Q''_i$, $D_i$ can also be approximated as a generalized Gaussian distribution.

$$\begin{aligned} D_i = R' - R''_i &= X' - X''_i + \mathcal{N}(0, 2) + Q' - Q''_i \\ &= \mathcal{G}(\mu_i, \alpha_i, \beta_i) + \mathcal{N}(0, 2) + Q' - Q''_i. \end{aligned} \tag{5}$$

Similar to fluorescence microscopy, in IMC images the tissue or background pixels should be continuous. Thus, for any normal pixel $p$ the distance between $d_i^p$ and the mean $\mu_i$ is always less than that of a single hot pixel, where $d_i^p$ is the pixel $p$'s value in the distribution $D_i$. Therefore, we can define $\triangle_i^p = |d_i^p - \mu_i|$ as the measure to determine whether a pixel $p$ is a hot pixel. However, this might not hold for consecutive hot pixels. For instance, if two consecutive hot pixels sharing similar intensities, their difference may be very close to $\mu_i$. To detect consecutive hot pixels, it is reasonable to assume there are at least $n$ pixels close to a normal pixel $p$. $n$ is normally set as 4, which corresponds to half neighbours. Consequently, we sort $\triangle_i^p$ for every pixel $p$ in an ascending direction as Eq. (6).

$$\triangle_{(i)}^p = \text{sort}(\triangle_i^p), \tag{6}$$

where $(i)$ is the sorted index and $i \in \{1, 2, ..., 8\}$. Then we define the sum of the first $n$ $\triangle_{(i)}^p$ as Eq. (7).

$$s_n^p = \sum_{i=1}^{n} \triangle_{(i)}^p, \tag{7}$$

For a normal pixel $p$, its $s_n^p$ should be less than that of a hot pixel. Because $s_n^p$ measures the relationship between the center pixel and its multiple neighbours, it is more robust than a single $\triangle_i^p$, especially for consecutive hot pixels.

Due to the spatial continuity and isotropic resolution of IMC images, $\mu_i$ from different directions can be regarded as equal to each other. For the sake of simplicity, we define $\mu_i = \mu$ for $i \in \{1, 2, ..., 8\}$. To further separate normal and hot pixels, Eq. (8) is consequently derived based on the triangle inequality [6].

$$s_n^p = \sum_{i=1}^{n} \left| d_{(i)}^p - \mu \right| \geq \left| t_n^p - n\mu \right|, \tag{8}$$

where $t_n^p = \sum_{i=1}^{n} d_{(i)}^p$. The equality holds only when $d_{(i)}^p \geq \mu$ or $d_{(i)}^p \leq \mu$ for all $i \in \{1, 2, ..., n\}$. The first case is always true for single hot pixels and pixels with the largest intensities in hot clusters. Otherwise, $t_n^p$ will shrink towards $n\mu$. Therefore, $t_n^p$ will further unmix hot pixels from normal ones. Combining all the $t_n^p$, a new distribution $T_n$ is generated, and outliers are located beyond its right tail. In fact, some consecutive hot pixels can shrink towards $n\mu$ as well. For example, in the case of a hot pixel that is larger than all of its normal neighbours but smaller than the largest hot pixel in the 3×3 sliding window, it is possible that $t_{n-1}^p - (n-1)\mu = \mu - d_{(n)}^p > 0$. To solve this issue, we implement multiple iterations of the sorting to adequately remove the hot pixel noise. Normally the iteration number $N_{iter}$ is set as 3 such that at least 3 consecutive hot pixels will be removed after 3 iterations.

##### 1.2.2 Hot pixel detection

In order to search outliers, the shape of $T_n$ should be investigated first. Therefore, let us also define $u_n^p = \sum_{i=1}^{n} d_i^p$, and combine $u_n^p$ from all pixels to form a distribution $U_n$. Without hot pixels, $U_n$ is a generalized Gaussian distribution with mean $n\mu$. Because

$$\sum_{p}\sum_{i=1}^{n}(\triangle_{(i)}^{p})^2 \leq \sum_{p}\sum_{i=1}^{n}(\triangle_{i}^{p})^2, \tag{9}$$

where the two items are proportional to the variances of $T_n$ and $U_n$, respectively, $T_n$ can be regarded as a generalized Gaussian distribution with mean $n\mu$ and smaller variance than $U_n$.

Because the histogram of $T_n$ is discretized, we apply the kernel density estimation algorithm [7], as Eq. (10) shows, to fit a continuous curve.

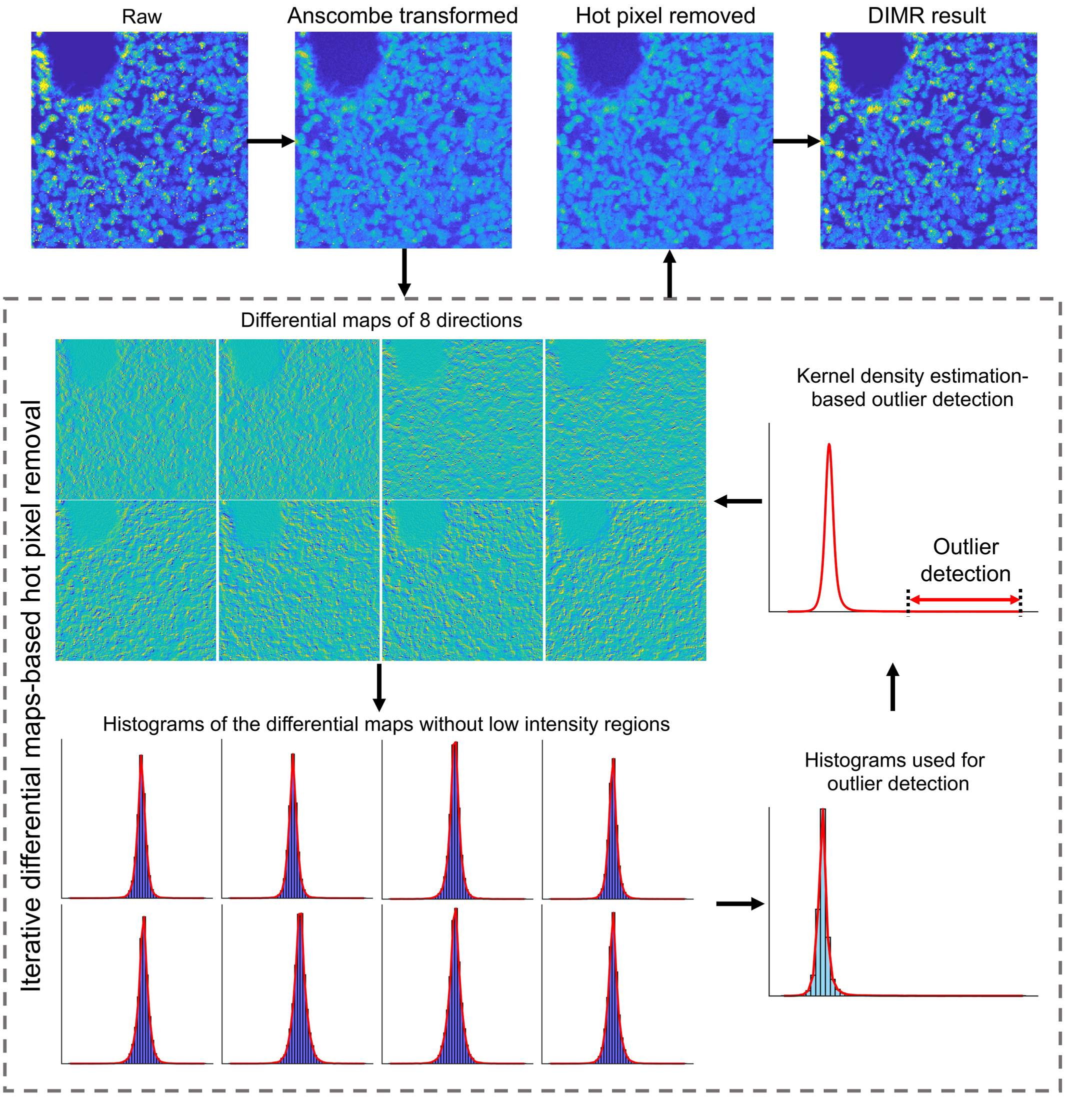

**Supplementary Figure 1.** The schematic of the DIMR algorithm.

$$\hat{g}_h(x) = \frac{1}{mh}\sum_{i=1}^{m} K(\frac{x - x_i}{h}), \tag{10}$$

where $K$ is a Gaussian kernel, $h$ the bandwidth, $m$ the point number and $x_i$ the sampled points with interval of 1 for adequate sampling. Subsequently, a moving mean filter with window size of 3 is utilized to further eliminate minor fluctuations of the fitted curve. The bandwidth $h$ is set as $1.06\hat{\sigma}n^{-\frac{1}{5}}$ [7], where $\hat{\sigma}$ is the standard deviation of the sampled points. With the fitted curve $(x, \hat{g}_h(x))$, a threshold point $x_T$ is defined and then any points $x > x_T$ are considered as outliers. Because outliers are located beyond the right tail of $T_n$, it is reasonable to set $x_T$ when $\frac{d\hat{g}_h(x)}{dx} \to 0$, which means the current distribution ends. Likewise, the shape of the curve will not change from convex to concave on its right tail. Thus, it is also reasonable to set $x_T$ when $\frac{d^2\hat{g}_h(x-dx)}{dx^2} \geq 0$ and $\frac{d^2\hat{g}_h(x)}{dx^2} \leq 0$. After the outliers are detected in each iteration, they are filtered by a 3×3 median filter. When the iterations terminate, the hot pixel removed image is transformed to its original scale by the direct algebraic inverse Anscombe transformation [8]. Additionally, we substitute the mean $\mu_i^p$ of each distribution $D_i$ with the median $\tilde{\mu}_i^p$ when implementing this algorithm, as median is more robust than mean when encountering outliers.

The DIMR algorithm is summarized as Supplementary Algorithm 1 and Supplementary Figure 1.

---

**Supplementary Algorithm 1** DIMR algorithm

---

**Input:** Raw image $R$;
    Hyper-parameters $n$ and $N_{iter}$;
**Output:** Hot pixel removed image;
1: Apply the Anscombe transformation to $R$;
2: **for** each $k = 1, 2, ..., N_{iter}$ **do**
3:     Calculate the 8 differential maps of $R'$ in a sliding $3 \times 3$ window;
4:     Remove the pixels with value less than 4 for each differential map and form the distributions $D_i$ where $i \in \{1, 2, ..., 8\}$;
5:     For every remaining pixel $p$, calculate $\triangle_i^p = |d_i^p - \tilde{\mu}_i|$, where $\tilde{\mu}_i$ is the median of $D_i$;
6:     Sort all the $\triangle_i^p$ and get the corresponding sorted index $(i)$ for every pixel $p$. With the index $(i)$, calculate $t_n^p$ for each pixel $p$ to form a new distribution $T_n$;
7:     Apply the kernel density estimation algorithm (Eq. (10)) followed by a moving mean filter to generate a continuous curve $(x, \hat{g}_h(x))$ from the histogram of $T_n$.
8:     Starting from the right tail of the curve, when (1) $\frac{d\hat{g}_h(x)}{dx} \to 0$ or (2) $\frac{d^2\hat{g}_h(x-dx)}{dx^2} \geq 0$ and $\frac{d^2\hat{g}_h(x)}{dx^2} \leq 0$, set the point $x$ as $x_T$;
9:     **if** There is no $x > x_T$ **then**
10:         Break
11:     **else**
12:         Set any points $x > x_T$ as outliers and filter the corresponding pixels with a 3×3 median filter.
13:     **end if**
14: **end for**
15: Apply the direct algebraic inverse Anscombe transformation to the processed image.

---

#### 1.3 Self-supervised deep learning-based algorithm for shot noise filtering (DeepSNF)

##### 1.3.1 Optimal loss function derivation

After hot pixel removal, the IMC imaging model is simplified as Eq. (11).

$$R = \mathcal{P}[X]. \tag{11}$$

In a supervised learning framework, the noisy and clean images $R$ and $X$ are both known. Therefore, for all the pixels $p$ from 1 to $P$, pairs of $(r_p, x_p)$ are formed as a training set. A deep convolutional neural network with forward model $\mathcal{F}_\theta$ ($\mathbb{E}^P \mapsto \mathbb{E}^P$) is then built to filter the Poisson noise such that $X = \mathcal{F}_\theta[R]$, where $\theta$ reflects the parameters of the neural network. Ideally, $\mathcal{F}_\theta[r_p]$ should be identical to that of $x_p$ for all pixels. In this sense, the optimal estimator for denoising can be derived through a binary signal detection task. This task requires an observer to classify the denoised signals $\mathcal{F}_\theta[r_p]$ under a hypothesis $H_0$ from the true clean signals $x_p$ under another hypothesis $H_1$. Given noisy data $r_p$, these two hypotheses can be described as:

$$\begin{aligned} &H_0 : \text{denoised signals} : \mathcal{F}_\theta[r_p], 1 \leq p \leq P \\ &H_1 : \text{true signals} : x_p, 1 \leq p \leq P \end{aligned} \tag{12}$$

Because the IMC data follows a Poisson distribution, the corresponding likelihood functions for these two hypotheses are expressed as Supplementary Eqs. (13) and (14). From Eq. (14), $x_p$ can be regarded as the maximum likelihood estimation (MLE) solution of $r_p$.

$$Pr(R|\mathcal{F}_\theta[R]) = \prod_{p=1}^{P} \exp(-\mathcal{F}_\theta[r_p])\mathcal{F}_\theta[r_p]^{r_p}/r_p!. \tag{13}$$

$$Pr(R|X) = \prod_{p=1}^{P} \exp(-x_p)x_p^{r_p}/r_p!. \tag{14}$$

The log-likelihood functions of the two hypotheses are derived as Supplementary Eqs. (15) and (16).

$$\mathrm{log}Pr(R|\mathcal{F}_\theta[R]) = \sum_{p=1}^{P}(r_p\mathrm{log}(\mathcal{F}_\theta[r_p]) - \mathcal{F}_\theta[r_p] - \mathrm{log}r_p!). \tag{15}$$

$$\mathrm{log}Pr(R|X) = \sum_{p=1}^{P}(r_p\mathrm{log}(x_p) - x_p - \mathrm{log}r_p!). \tag{16}$$

Then, we define the log-likelihood ratio $\mathcal{L}(H_0, H_1)$ to measure the difference between the two hypotheses:

$$\begin{aligned}\mathcal{L}(H_0, H_1) &= \mathrm{log}Pr(R|X) - \mathrm{log}Pr(R|\mathcal{F}_\theta[R]) \\ &= \sum_{p=1}^{P}(r_p\mathrm{log}\frac{x_p}{\mathcal{F}_\theta[r_p]} - x_p + \mathcal{F}_\theta[r_p]).\end{aligned} \tag{17}$$

The learning process aims to train $\mathcal{F}_\theta$ such that $x_p = \mathcal{F}_\theta[r_p]$ for all the $p$ from 1 to $P$. Thus, the expectation of this loss function $\mathcal{L}$ under hypothesis $H_1$, $E[\mathcal{L}(H_0, H_1)|H_1]$, as Eq. (18) demonstrates, should be as small as possible. In that sense, the observer in the binary detection task will be more difficult to differ these two hypotheses as $E[\mathcal{L}(H_0, H_1)|H_1]$ decreases. When $E[\mathcal{L}(H_0, H_1)|H_1] = 0$, the two hypotheses are identical and the denoised signal $\mathcal{F}_\theta[r_p]$ will be identical to the true signal $x_p$ for all the pixels, which means $\mathcal{F}_\theta[r_p]$ will be the MLE solution of $r_p$ as well.

$$\begin{aligned}E[\mathcal{L}(H_0, H_1)|H_1] &= \sum_{p=1}^{P}(E[r_p|H_1]\mathrm{log}\frac{x_p}{\mathcal{F}_\theta[r_p]} - x_p + \mathcal{F}_\theta[r_p]) \\ &= \sum_{p=1}^{P}(x_p\mathrm{log}\frac{x_p}{\mathcal{F}_\theta[r_p]} - x_p + \mathcal{F}_\theta[r_p]).\end{aligned} \tag{18}$$

Eq. (18) is generally known as I-divergence [9]. Here, we set it as the loss function, and thus the optimal parameter of the network $\hat{\theta}^*$ is defined as Eq. (19).

$$\hat{\theta}^* = argmin_\theta\sum_{p=1}^{P}(x_p\mathrm{log}\frac{x_p}{\mathcal{F}_\theta[r_p]} - x_p + \mathcal{F}_\theta[r_p]). \tag{19}$$

Due to the difficulties associated with acquiring high SNR images as ground truths and the impossibility to repetitively scan the same tissue in IMC, a supervised learning approach is unavailable here. Fortunately, we find self-supervised approaches are also qualified for this denoising task. Let us define a function $f$ demonstrating a random pixel masking approach for images such as those in Noise2Void [10] and Noise2Self [11]. In such strategies, multiple pixels are randomly masked and replaced by their adjacent pixels or random values. The new pixel value $f(r_p)$ at pixel $p$ can be regarded as the true value $x_p$ contaminated by another noise process. In this case, the true value $x_p$ can be predicted by the adjacent pixels because of the spatial continuity of IMC images. Therefore, in a self-supervised learning problem, the loss function will be

$$\begin{aligned}\mathcal{L}(R, \mathcal{F}_\theta[f(R)]) &= \sum_{p=1}^{P}(r_p \log \frac{r_p}{\mathcal{F}_\theta[f(r_p)]} - r_p + \mathcal{F}_\theta[f(r_p)]) \\ &= \sum_{p=1}^{P}(r_p \log r_p - (x_p + r_p - x_p)\log \mathcal{F}_\theta[f(r_p)] - (x_p + r_p - x_p) + \mathcal{F}_\theta[f(r_p)]) \\ &= \sum_{p=1}^{P}(x_p \log \frac{x_p}{\mathcal{F}_\theta[f(r_p)]} - x_p + \mathcal{F}_\theta[f(r_p)]) + \sum_{p=1}^{P}(r_p \log r_p - x_p \log x_p) \\ &\quad - \sum_{p=1}^{P}(r_p - x_p) - \sum_{p=1}^{P}(r_p - x_p)\log \mathcal{F}_\theta[f(r_p)]. \end{aligned} \tag{20}$$

In order to ensure that the training process with the self-supervised learning loss is identical to that with known true signals, the last three terms in Eq. (20) should be constants. Obviously, the second term $\sum_{p=1}^{P}(r_p \log r_p - x_p \log x_p)$ fulfills this requirement. On the other hand, in the last two terms, $r_p - x_p$ and $\mathcal{F}_\theta[f(r_p)]$ are the noise component and predicted value of pixel $p$, respectively. Because the Poisson noise is a pixel-independent stochastic process and $\mathcal{F}_\theta[f(r_p)]$ is determined by its neighbours in the proposed self-supervised learning framework, they are uncorrelated with each other, except the masked pixel $p$ is replaced by itself. As a result, the last two term in Eq. (20) can be approximated as Eq. (21).

$$\begin{aligned}\sum_{p=1}^{P}(r_p - x_p) + \sum_{p=1}^{P}(r_p - x_p)\log \mathcal{F}_\theta[f(r_p)] &= \sum_{p=1}^{P}(r_p - x_p) + \sum_{p=1}^{P}(r_p - x_p)\sum_{p=1}^{P}\log \mathcal{F}_\theta[f(r_p)] \\ &= \sum_{p=1}^{P}(r_p - x_p)\left[1 + \sum_{p=1}^{P}\log \mathcal{F}_\theta[f(r_p)]\right]. \end{aligned} \tag{21}$$

Because $E[r_p] = E[x_p]$ for a Poisson noise model, Eq. (21) is approximated as 0 when $P$ is large enough, also suggested by [11]. From this viewpoint, training with noisy images and their masked pairs is identical to training with noisy and clean image pairs, as long as the masked pixels are not replaced by themselves. To avoid learning identity in the training, only the loss of the masked pixels are considered, as

Eq. (22) shows.

$$\mathcal{L}(R, \mathcal{F}_\theta[f(R)]) = \sum_p M_p \cdot \left[r_p \log\frac{r_p}{\mathcal{F}_\theta[f(r_p)]} - r_p + \mathcal{F}_\theta[f(r_p)]\right] \Big/ \sum_p M_p, \tag{22}$$

where $M$ is the pixel mask ($M \in \{0, 1\}$). If $M_p = 1$, then the pixel $p$ is masked; otherwise, it is not. To guarantee the output is non-negative, a softplus function $\log(1 + \exp(x))$ is set as the activation of the network's output layer.

##### 1.3.2 Hessian regularization as a booster for the denoising task

Even with the optimal estimator, the denoising performance is still sub-optimal: in the training process, the information of the masked pixels is always neglected; and only partial pixels are utilized. These limitations hinder the trained denoiser to further enhance its performance. To solve this issue, the characteristics of the IMC images should be utilized in the training process. As an imaging modality similar to traditional fluorescence microscopy, IMC is always used for detecting the phenotype of biological structures. Thus, the spatial continuity between biological structures can be used as a *priori*. In fact, the Hessian regularization is widely used to describe such spatial continuity in biological imaging data [2, 12, 13]. However, to the best of our knowledge, this advanced statistical prior has never been used in deep learning-based denoising task. To implement this prior in IMC images, let us first define the Hessian operator $\mathcal{R}_{Hessian}$ as Eq.(23).

$$\mathcal{R}_{Hessian} = \begin{bmatrix} \partial_{xx} & \partial_{xy} \\ \partial_{yx} & \partial_{yy} \end{bmatrix}, \tag{23}$$

where $\partial_{xx} = \partial^2/\partial x^2$, $\partial_{xy} = \partial_{yx} = \partial^2/\partial x \partial y$, $\partial_{yy} = \partial^2/\partial y^2$, and $x$ and $y$ are the vertical and horizontal directions of an image, respectively. Then, for any estimated image $\mathcal{F}_\theta[f(R)]$, the corresponding Hessian regularization term is defined as Eq. (24).

$$|\mathcal{R}_{Hessian}(\mathcal{F}_\theta[f(R)])| = |\partial_{xx}\mathcal{F}_\theta[f(R)]| + |\partial_{yy}\mathcal{F}_\theta[f(R)]| + \sqrt{2}|\partial_{xy}\mathcal{F}_\theta[f(R)]| \tag{24}$$

When involving the Hessian regularization as a prior, the optimization problem will convert from Eq.

(18) to Eq. (25).

$$\hat{\theta}^* = argmin_\theta \sum_p \big[E[\mathcal{L}(H_0, H_1)|H_1] - \lambda_{Hessian} \log Pr_{Hessian}(r_p;\theta)\big], \tag{25}$$

where $Pr_{Hessian}(r_p;\theta) = \exp(-|\mathcal{R}_{Hessian}(\mathcal{F}_\theta[f(R)])|_p)$ and $\lambda_{Hessian}$ is the Hessian regularization parameter. Therefore, the loss function Eq. (22) will be

$$\begin{aligned}\mathcal{L}(R, \mathcal{F}_\theta[f(R)]) = &\sum_p M_p \cdot \Big[r_p \log \frac{r_p}{\mathcal{F}_\theta[f(r_p)]} - r_p + \mathcal{F}_\theta[f(r_p)]\Big] \Big/ \sum_p M_p \\ &+ \lambda_{Hessian} \sum_p |\mathcal{R}_{Hessian}(\mathcal{F}_\theta[f(R)])|_p \Big/ \sum_p .\end{aligned} \tag{26}$$

Note that the first term works only for the selected masked pixels, while the second regularization term utilizes all the information of images. Thus, Eq. (26) overcomes the limitations of Eq. (22) and further enhances the performance of the denoiser.

##### 1.3.3 Image normalization

For both training and prediction, it is important to normalize the input images to a common range. However, some cells or structures may still exhibit extremely bright signals even after hot pixel removal. Consequently, all input and output data are percentile-normalized between 0 and 1. IMC images are always non-negative and consist of pixels with 0 value, this normalization is defined as Eq. (27).

$$\mathrm{N}(R;q) = \frac{R}{1.1 \times perc(R,q)}, \tag{27}$$

where $perc(R;q)$ is the $q$-th percentile of all pixel values in the training set. We typically use values of $q \in (99.9, 99.999)$. Note that only in training phase any values which are larger than 1 are set as 1. After prediction, the denoised images are re-transformed to their original scales.

This normalization approach does not affect the selection of regularization parameter $\lambda_{Hessian}$. The Hessian operator (23) is a linear operator, so we have

$$|\mathcal{R}_{Hessian}(\alpha_{scale}\mathcal{F}_\theta[f(R)])| = \alpha_{scale}|\mathcal{R}_{Hessian}(\mathcal{F}_\theta[f(R)])|, \tag{28}$$

where $\alpha_{scale} = 1/(1.1 \times perc(R; q))$. From here, we are able to derive

$$
\begin{aligned}
&\mathcal{L}(\alpha_{scale}R, [\mathcal{F}_\theta[f(R)]]_{scaled}) \\
=&\sum_p M_p \cdot \left[\alpha_{scale} r_p \log \frac{\alpha_{scale} r_p}{[\mathcal{F}_\theta[f(r_p)]]_{scaled}} - \alpha_{scale} r_p + [\mathcal{F}_\theta[f(r_p)]]_{scaled}\right] \Big/ \sum_p M_p \\
&+ \lambda_{Hessian} \sum_p |\mathcal{R}_{Hessian}(\mathcal{F}_\theta[f(R)])|_p \Big/ \sum_p \\
=&\alpha_{scale} \left\{ \sum_p M_p \cdot \left[r_p \log \frac{r_p}{[\mathcal{F}_\theta[f(r_p)]]_{scaled}/\alpha_{scale}} - r_p + [\mathcal{F}_\theta[f(r_p)]]_{scaled}/\alpha_{scale}\right] \Big/ \sum_p M_p \right. \\
&\left. + \lambda_{Hessian}\alpha_{scale} \sum_p |\mathcal{R}_{Hessian}(\mathcal{F}_\theta[f(R)]/\alpha_{scale})|_p \Big/ \sum_p \right\}. \qquad (29)
\end{aligned}
$$

Therefore, $[\mathcal{F}_\theta[f(r_p)]]_{scaled} = \alpha_{scale}\mathcal{F}_\theta[f(r_p)]$ and the normalization does not affect the strength of regularization.

The DeepSNF algorithm is summarized as Supplementary Algorithm 2.

**Supplementary Algorithm 2** DeepSNF algorithm

**Input:** Hot pixel removed images $R$;
**Output:** Noise filtered images;

1: Generate a training set for a specific hot pixel removed marker channel, in which all the images are percentile normalized between 0 and 1 with Eq. (27);
2: Train a denoising network for the marker channel with Eq. (26) as the loss function;
3: Normalize the hot pixel removed images with the pre-calculated maximum of the training set;
4: Filter shot noise for the normalized images with the trained network;
5: De-normalize the predicted images to their original scales.

### Supplementary Note 2: Reference methods

#### 2.1 Hot pixel removal methods

Currently, two thresholding methods are mostly applied to remove hot pixels, which are neighbour-based threshold hot pixel removal method [14–16] (NTHM) and median-based threshold hot pixel removal method [17] (MTHM). NTHM is simple and straightforward. It pre-sets a threshold $\sigma_{thresh}$. Then single pixels with intensity greater than this threshold of the maximum value in its $3 \times 3$ neighbourhood are detected as outliers. This method is very similar to our DIMR algorithm with hyper-parameter $n = 1$, which will always overlook consecutive hot pixels. In addition, $\sigma_{thresh}$ needs to be manually set. However, the hot pixels in different tissues and channels have different scales. That is, one fixed threshold does not work for different images. These limitations downgrades its performance in real applications. The NTHM algorithm is summarized as Supplementary Algorithm 3, and the default value of $\sigma_{thresh}$ is set 50.

**Supplementary Algorithm 3** NTHM for hot pixel removal

**Input:** Raw image $R$;
Hyper-parameter $\sigma_{thresh}$;
**Output:** Hot pixel removed image;
1: In a sliding $3 \times 3$ window, select the maximum value after excluding the center pixel;
2: **if** The difference between the center pixel and the maximum value is larger than $\sigma_{thresh}$ **then**
3: Substitute the center pixel's value with the maximum value.
4: **end if**

MTHM is an automated approach and can remove consecutive hot pixels, of which two thresholds $\sigma_{thresh1}$ and $\sigma_{thresh2}$ are needed to be manually set. This method first searches the top $\sigma_{thresh1}$ pixels and regards any pixels as outliers if they are $\sigma_{thresh2}$ times higher than the median in a sliding $5 \times 5$ window. However, this method may falsely remove normal pixels located at the border between tissues and background. Besides, with a larger $\sigma_{thresh1}$ or smaller $\sigma_{thresh2}$, false negatives may also be generated. Normally, $\sigma_{thresh1}$ and $\sigma_{thresh2}$ are set as 2% and 4 separately [17]. This algorithm is summarized as Supplementary Algorithm 4. The default values of $\sigma_{thresh1}$ and $\sigma_{thresh2}$ are set as 2% and 4, respectively.

**Supplementary Algorithm 4** MTHM for hot pixel removal

**Input:** Raw image $R$;
Hyper-parameters $\sigma_{thresh1}$ and $\sigma_{thresh2}$;
**Output:** Hot pixel removed image;
1: Search all the pixels with top $\sigma_{thresh1}$ values in an image;
2: **if** The pixels are larger than $\sigma_{thresh2}$ times of the medians in their $5 \times 5$ window **then**
3: Substitute the pixels' values with the medians.
4: **end if**

#### 2.2 Deep learning-based shot noise filtering methods

##### 2.2.1 Noise2Void

Noise2Void uses noisy images as the input and output to train a denoising network. It randomly masks several pixels and replaces them with their neighbours. As Eq. (30) shows, minimizing the mean squared error (MSE) between the noisy image pairs is equal to minimizing that of the noisy and clean image pairs plus the variance of the noise. This equality holds as long as the function $\mathcal{F}$ is "$\mathcal{J}$-invariant" [11], which means $\mathcal{F}[f(r_p)] - x_p$ and $r_p - x_p$ are independent for any pixel $p$. Notably, the current masking strategies always fulfill this requirement if the noise is pixel-independent and the pixels are not replaced by themselves.

$$\mathcal{L}_{MSE}(R, \mathcal{F}_\theta[R_{masked}]) = E(\mathcal{F}_\theta[f(r_p)] - r_p)^2 \tag{30}$$

$$= E(\mathcal{F}_\theta[f(r_p)] - x_p)^2 + E(r_p - x_p)^2 \tag{31}$$

In Eq. (31), the second term corresponds to the noise variance at pixel $p$, which is a constant for a Gaussian noise model. In this condition, it can be neglected in the training process, and hence the self-supervised learning loss is equal to a supervised one.

However, this statement is not true for a Poisson noise model. The variance of Poisson noise is equal to its true signal intensity, i.e., $E(r_p - x_p)^2 = E(x_p)$. Therefore, the pixels with higher signal will gain more attention in Eq. (31), resulting in sub-optimal denoising for signal-limited area. In fact, even in a supervised learning task, MSE is still not an unbiased estimator for Poisson noise. When true signals are available, the optimizer can be derived by the MLE framework:

$$\hat{\theta}_{MLE} = argmin_\theta E\big[-\log\big(Pr(r_p|x_p;\theta)\big)\big]. \tag{32}$$

Approximating a Poisson process as Gaussian distribution, Eq. (32) is converted as

$$\hat{\theta}_{MLE} = argmin_{\theta} E\big[\frac{1}{2}(\mathcal{F}_{\theta}(r_p) - x_p)^2/x_p + \frac{1}{2}\log(2\pi x_p)\big]. \tag{33}$$

Obviously, minimizing this loss in Eq. (33) is not equivalent to minimizing the MSE between $\mathcal{F}_{\theta}(r_p)$ and $x_p$, because $x_p$ varies with different pixels $p$. As a result, MSE loss will generate bias in Poisson denoising. Beside this issue, in the original version of Noise2Void, a linear activation is used as the output layer's activation function of the network, which violates the non-negativity of IMC images.

##### 2.2.2 Modified Noise2Void with the Anscombe transformation and rectified linear unit (ReLU) activation

We propose a modified Noise2Void algorithm to correct the bias of Noise2Void. The Anscombe transformation is first applied to the IMC data, so that the Poisson noise is approximated as a Gaussian-distributed noise model. In this sense, the noise variance term in Eq. (30) can be regarded as a constant and the bias from MSE loss will be mitigated. By substituting linear activation with ReLU [18], the non-negativity of IMC images is satistied. Note that the denoised images should be re-transformed to their original scale by the exact unbiased inverse Anscombe transformation [8]. The exact unbiased inverse Anscombe transformation software package was downloaded from https://webpages.tuni.fi/foi/invansc/.

However, the Anscombe transformation is not accurate for very low values [4]. Unfortunately, there are usually a portion of pixels in IMC images suffering from extremely low counts. The variances of these pixels are still positively correlated with the counts. Thus, even this approach reduces the bias of Noise2Void, some errors are still inevitable.

Compared to the MSE loss used by Noise2Void and Noise2Self, our derived loss function is the optimal estimator for Poisson denoising and the corresponding outputs are inherently non-negative with a softplus function. Consequently, it does not generate any biases for IMC denoising.

##### 2.2.3 Noise2True

In Noise2True [19], clean images are available as ground truths, so that a supervised learning is possible. To achieve a learning with the optimal estimator, we use our derived I-divergence Eq. (18) as the loss function in Noise2True. In simulation, we compare all the above denoising methods with Noise2True. We are especially curious how our Hessian regularization could help boost self-supervised learning to approach the performance of Noise2True. The ground truths of IMC images are only easy to acquire in simulation. Thus, Noise2True is in fact not possible in real IMC denoising.

#### 2.3 Traditional statistics-based shot noise filtering methods

##### 2.3.1 Gaussian filter

Gaussian filter might be the most widely used noise filter. For all of our IMC images, we apply a Gaussian filter with kernel size of $5 \times 5$ and standard deviation of 0.8. Gaussian filter can only remove the high-frequency noise with the risk of filtering lots of true signal. As a result, its performance is sub-optimal than other smarter denoising algorithms.

##### 2.3.2 Non-local means (NLM) algorithm

NLM algorithm [20] takes a mean of all pixels in an image, weighted by how similar these pixels are to the target pixel. In our paper, this algorithm is implemented using the Matlab built-in function “imnlmfilt”. All the parameters are set as default.

##### 2.3.3 Batch-matching and 3D filtering (BM3D) algorithm with Anscombe transformation

BM3D [21] is modified from NLM algorithm, which is usually regarded as a state-of-the-art denoising algorithm due to its superior performance in multiple applications. Instead of simply average the pixel values, it collects similar patches of an image, and then apply hard thresholding and Wiener filtering in two stages. BM3D is built on a white Gaussian noise model. Therefore, the Anscombe transformation is applied to approximate the Poisson noise in IMC images to Gaussian noise. The same as the modified Noise2Void algorithm, the exact unbiased inverse Anscombe transformation is also applied to rescale the denoised images.

The BM3D algorithm software package was downloaded from https://webpages.tuni.fi/foi/GCF-BM3D/index.html#ref_software. When implementing BM3D, we set $\sigma_{noise} = 1$, $N_2 = 8$, $N_s = 17$, $\tau_{match} = 2500$, $\lambda_{thr3D} = 1$, $Ns_{wiener} = 25$ and $\tau_{match-weiner} = 600$. All the other parameters were set as default.

### Supplementary Note 3: Simulation

Due to the difficulty to acquire ground truths, it is infeasible to quantitatively evaluate the accuracy of hot pixel removal methods and the shot noise filtering algorithms with real IMC images. As a consequence, we propose to conduct a comprehensive and accurate quantitative evaluation with simulated data.

#### 3.1 Simulated data generation

We modify Eq. (2) as Eq. (34) to generate simulated data.

$$R = \mathcal{P}[X_{origin}/\gamma] + Q, \tag{34}$$

where $X_{origin}$ represent the original clean images used for simulation and $\gamma$ is the scale factor to control the overall ion counts level. A larger $\alpha$ indicates the overall ion counts of the ground truth image $X_{GT} = X_{origin}/\gamma$ is lower, and hence the shot noise level is higher. On the other hand, we assume hot pixels $Q$ obey a negative binomial distribution $NB(\tau, \eta)$. Additionally, another parameter $\omega$ is used to indicate the density of hot pixels.

Here we select the original clean images $X_{origin}$ from the $t$-CYCIF dataset [22] because of their very high SNR and similar resolution (1.06 $\mu$m) with IMC images. In particular, we choose a cell marker CD14 and structural marker Keratin from a lung tissue to evaluate the shot noise filtering algorithms. These two channels along with a DNA channel are used to evaluate the hot pixel removal methods.

We have four parameters $\gamma$, $\tau$, $\eta$ and $\omega$ to determine the noise conditions of the simulated images. For each channel, we set 4 different $\gamma$ which correspond to 4 different SNR levels including high, medium, low and very low. With different noise levels we set different $\tau$, $\eta$ and $\omega$ to represent different conditions of hot pixels. All the parameters are listed in Supplementary Table 1.

**Supplementary Table 1.** Simulation parameters

| Noise settings | | 1 | | | 2 | | | 3 | | | 4 | | |
|---|---|---|---|---|---|---|---|---|---|---|---|---|---|
| SNR levels | | high | | | medium | | | low | | | very low | | |
| Parameters | $\tau$ | $\gamma$ | $\eta$ | $\omega$ | $\gamma$ | $\eta$ | $\omega$ | $\gamma$ | $\eta$ | $\omega$ | $\gamma$ | $\eta$ | $\omega$ |
| CD14 & Keratin | 3 | 2000 | 0.05 | 1% | 5000 | 0.1 | 0.1% | 10000 | 0.2 | 0.01% | 16000 | 0.3 | 0.01% |
| DNA | 5 | 500 | 0.05 | 3% | 1000 | 0.1 | 1% | 2000 | 0.2 | 0.1% | 5000 | 0.25 | 0.01% |

#### 3.2 Accuracy metrics and statistical analysis in simulation

In simulation, the root mean squared error (RMSE) is used to evaluate the performance of the hot pixel removal methods, as Eq. (35) shows, in which $p$ is the pixel index, $Y^{clean}$ and $Y^{HM}$ are the simulated images without hot pixels and hot pixel removed images, separately.

$$\mathrm{RMSE}(Y^{HM}, Y^{clean}) = \sqrt{\frac{1}{P}\sum_{p=1}^{P}(Y_p^{HM} - Y_p^{clean})^2} \tag{35}$$

The peak signal-to-noise ratio (PSNR) and structural similarity (SSIM) [23] are used to evaluate the performance of the shot noise filtering algorithms. PSNR indicates the ratio between the maximum possible power of a signal and the power of corrupting noise that affects the fidelity of its representation. It is defined as Eq. (36), where $Y^{est}$ is the estimated image from denoising algorithms and $Y^{true}$ is the ground truth.

$$\mathrm{PSNR}(Y^{est}, Y^{true}) = 20\mathrm{log}\frac{\max(Y^{true})}{\sqrt{\frac{1}{P}\sum_{p=1}^{P}(Y_p^{est} - Y_p^{true})^2}} \tag{36}$$

The structure similarity [23] (SSIM) is a perception-based model that considers image degradation as perceived change in structural information, while also incorporating important perceptual phenomena, including both luminance masking and contrast masking terms. Compared to PSNR, it is supposed to give more information about image distortion by the computation of local image structure, luminance and contrast into a single local quality score. In this paper, the luminance and contrast are normalized and SSIM is defined as Eq. (37),

$$\mathrm{SSIM}(Y^{est}, Y^{true}) = \frac{2\mu_{Y^{est}}\mu_{Y^{true}} + C_1}{\mu^2_{Y^{est}} + \mu^2_{Y^{true}} + C_1} \cdot \frac{2\sigma_{Y^{est}Y^{true}} + C_2}{\sigma^2_{Y^{est}} + \sigma^2_{Y^{true}} + C_2} \tag{37}$$

where $Y^{est}$ is the estimated image from denoising algorithms, $Y^{true}$ is the ground truth, $\mu_{Y^{est}}$, $\mu_{Y^{true}}$, $\sigma_{Y^{est}}$, $\sigma_{Y^{true}}$ and $\sigma_{Y^{est}Y^{true}}$ are the local means, standard deviations and cross-covariance for images $Y^{est}$ and $Y^{true}$, $C_1$ and $C_2$ are the regularization constants to avoid instability for image regions where the local mean or standard deviation is close to zero.

In simulation, all the RMSE, PSNR and SSIM data are presented as box-and-whisker plots (center line, median; limits, 75% and 25% whiskers, maximum and minimum) along with all the data points. We use the paired one-way analysis of variation to do the multiple comparisons of these accuracy metrics. All the statistical tests are implemented with Prism 9 (GraphPad Software Inc.). Statistical significance at P<0.05,

0.01, 0.001 and 0.0001 are denoted by *, **, *** and ****, respectively. “ns” means “no significance”.

##### 3.3 Hot pixel removal methods evaluation

We benchmarked our DIMR algorithm with NTHM and MTHM on the simulated CD14, Kerain and DNA datasets. Each marker contains 4 hot pixel conditions and each condition contains 50 images. We set the hyperparameters of DIMR, $n$ and $N_{iter}$, as 4 and 3 for all the datasets. The hyperparameters of NTHM and MTHM were optimized for each noise setting to guarantee their best performance.

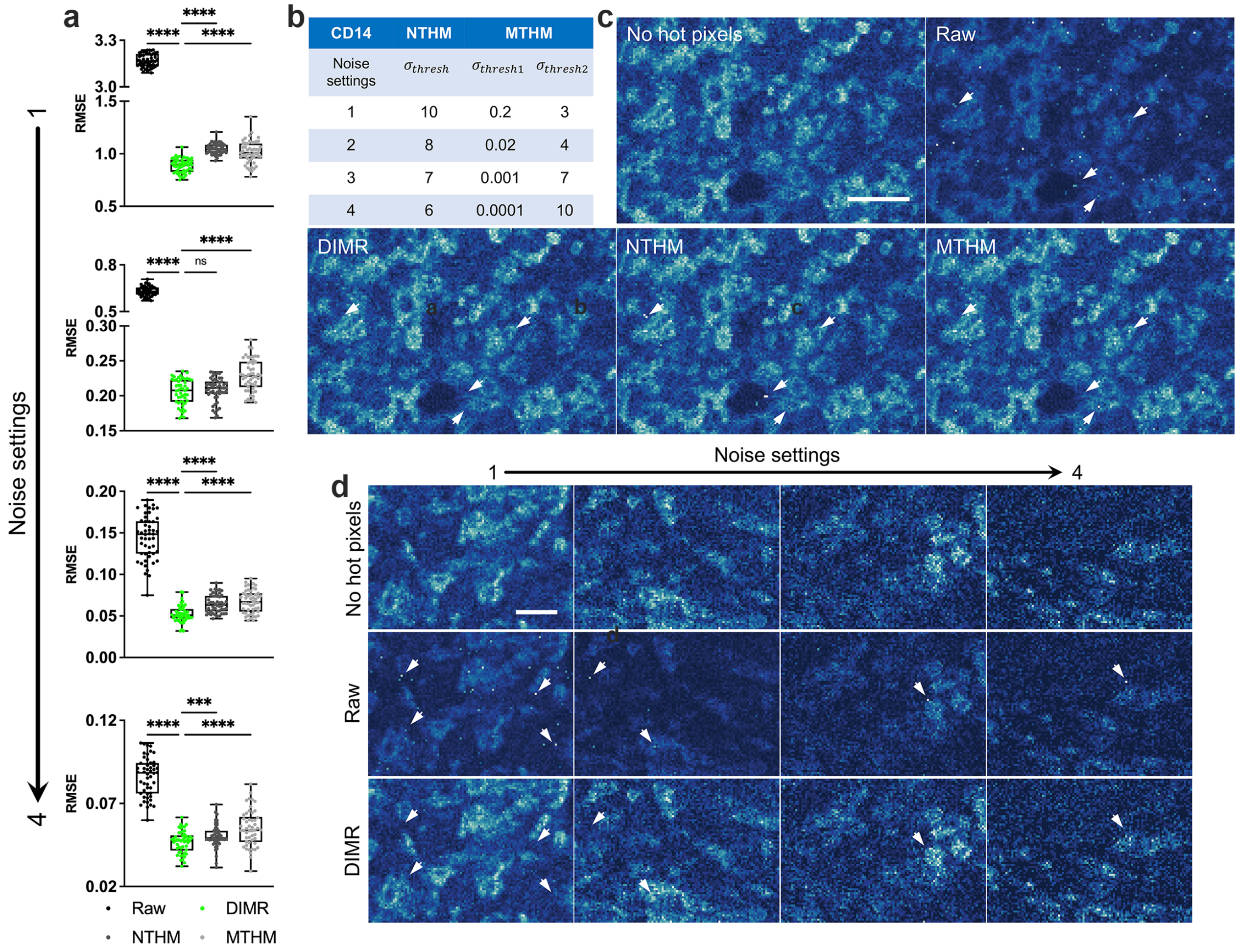

**Supplementary Figure 2.** Evaluation of the hot pixel removal methods on the simulated CD14 dataset ($N = 50$ per noise setting). (a) RMSE comparison under different noise settings. (b) The optimal hyper-parameters of NTHM and MTHM under different noise settings. (c) Visual inspection of the three hot pixel removal methods on a CD14 image with the highest hot pixel density under noise setting 1. (d) Visual inspection of the DIMR algorithm under different noise settings. Scale bar: (c) 20 $\mu$m, (d) 20 $\mu$m.

The evaluation results for these three markers are listed in Supplementary Figs. 2, 3 and 4, respectively. As these figures suggest, DIMR always removes hot pixels effectively and outperforms NTHM and MTHM on RMSE under different hot pixel conditions. Besides, the optimal hyper-parameters of NTHM and MTHM vary in a wide range with different settings of hot pixels (Supplementary Figs. 2b, 3b and 4b), while those of DIMR remain the same ($n = 4$ and $N = 3$). In fact, the hot pixel conditions vary under different images and markers. Therefore, it is labor-intensive to tune the hyper-parameters for every image. In comparison, the outlier detection of DIMR is based on the overall statistical features of the images, so that it is not essential to tune its hyper-parameters. To summarize, the DIMR algorithm is more accurate and flexible for removing hot pixels in IMC images.

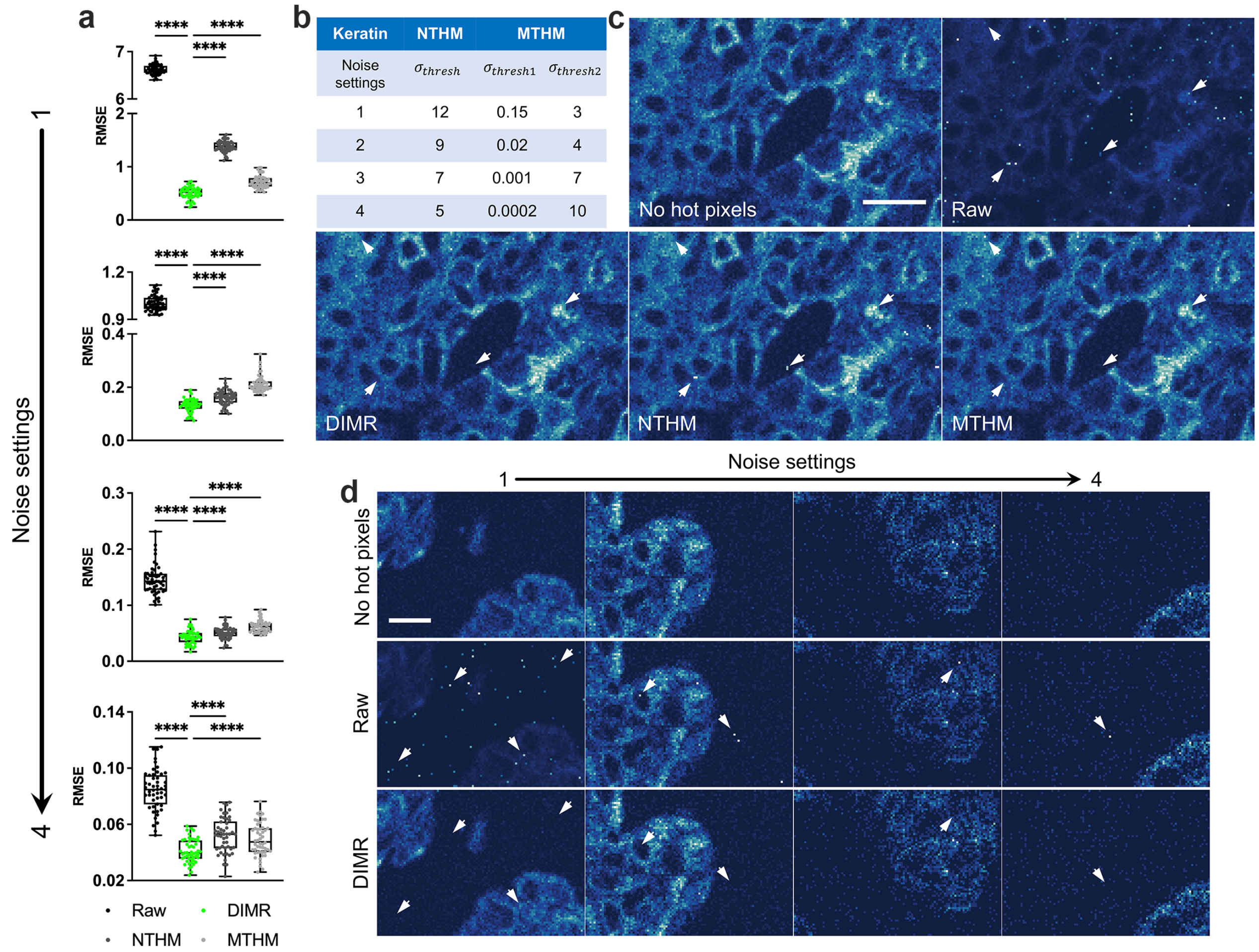

**Supplementary Figure 3.** Evaluation of the hot pixel removal methods on the simulated Keratin dataset ($N = 50$ per noise setting). (a) RMSE comparison under different noise settings. (b) The optimal hyper-parameters of NTHM and MTHM under different noise settings. (c) Visual inspection of the three hot pixel removal methods on a Keratin image with the highest hot pixel density under noise setting 1. (d) Visual inspection of the DIMR algorithm under different noise settings. Scale bar: (c) 20 $\mu$m, (d) 20 $\mu$m.

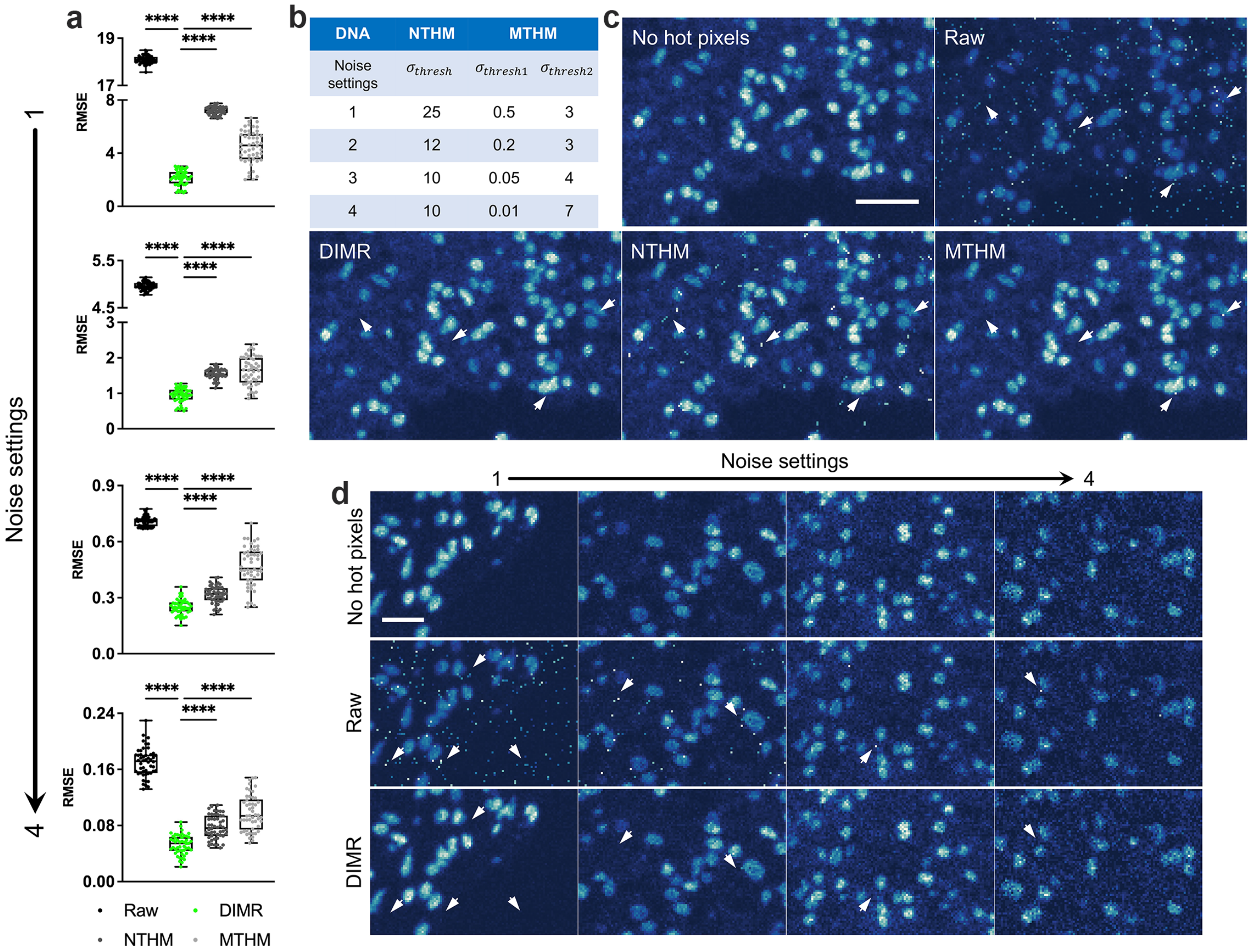

**Supplementary Figure 4.** Evaluation of the hot pixel removal methods on the simulated DNA dataset ($N = 50$ per noise setting). (a) RMSE comparison under different noise settings. (b) The optimal hyper-parameters of NTHM and MTHM under different noise settings. (c) Visual inspection of the three hot pixel removal methods on a DNA image with the highest hot pixel density under noise setting 1. (d) Visual inspection of the DIMR algorithm under different noise settings. Scale bar: (c) 20 $\mu$m, (d) 20 $\mu$m.

#### 3.4 Shot noise filtering methods evaluation

##### 3.4.1 Compare DeepSNF with Noise2Void and modified Noise2Void

In this part, we compared our DeepSNF algorithm to Noise2Void (N2V), modified Noise2Void (MN2V) and Noise2True (N2T) on the simulated CD14 and Keratin images under 4 different noise levels. For each condition, we generated training sets and trained separate networks with 25 images for DeepSNF, N2V, MN2V and N2T (Supplementary Table 6), and restored the other 25 images with the trained denoisers. Note that here DeepSNF with no regularization (DeepSNF-NR) was applied because we aimed to compare the performances of MSE and Eq. (18) as loss functions in the IMC denoising task. The denoised images were evaluated both visually and quantitatively, shown in Supplementary Figs. 5 and 6.

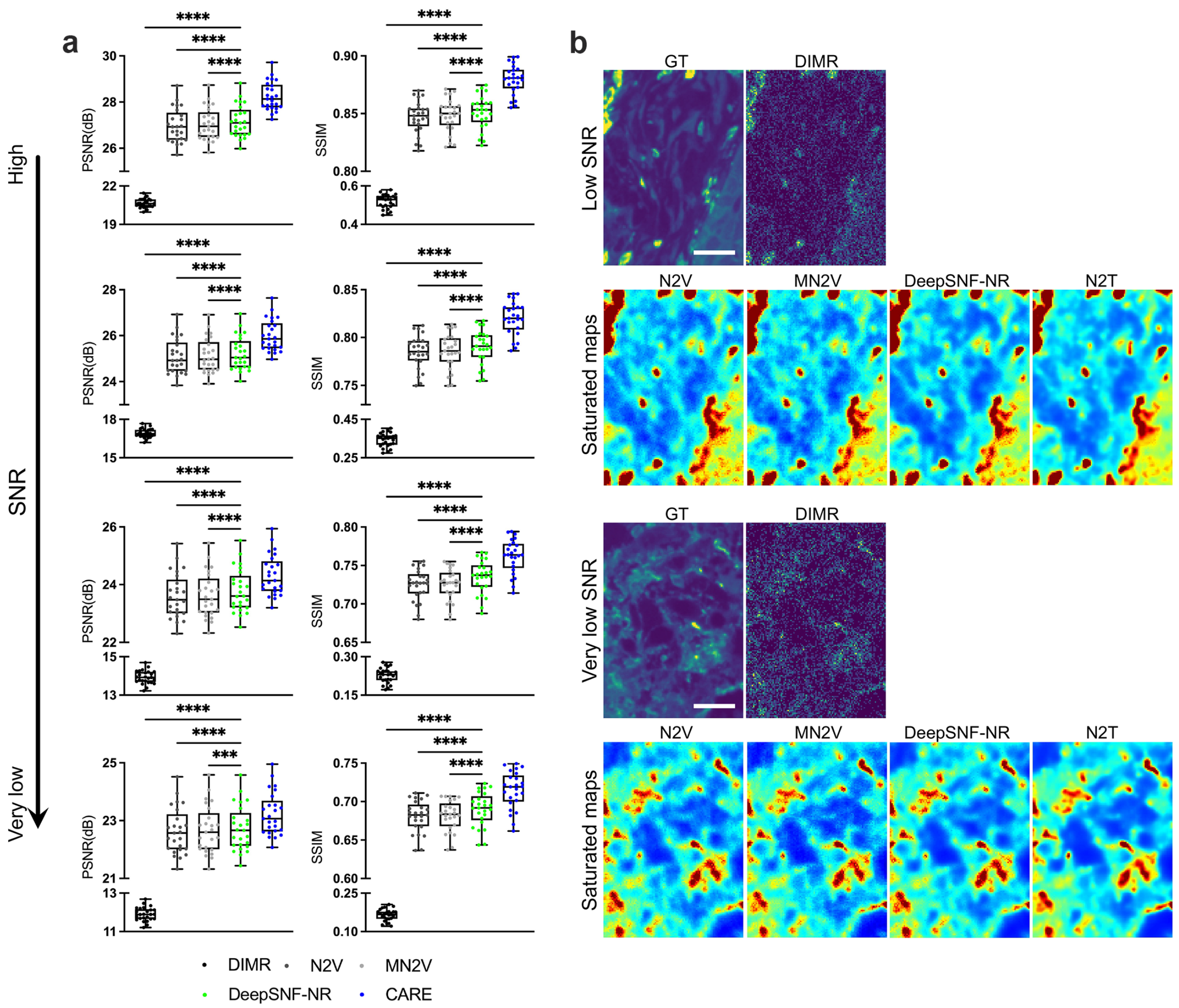

**Supplementary Figure 5.** Comparison of DeepSNF-NR with Noise2Void and modified Noise2Void on the simulated CD14 dataset with 4 noise levels ($N = 25$ per level). (a) PSNR and SSIM evaluation on the three algorithms. (b) Visual inspection of the three algorithms and N2T on denoising a low SNR image. (c) Visual inspection of the three algorithms and N2T on denoising a very low SNR image. Scale bar: (b) 20 $\mu$m, (c) 20 $\mu$m.

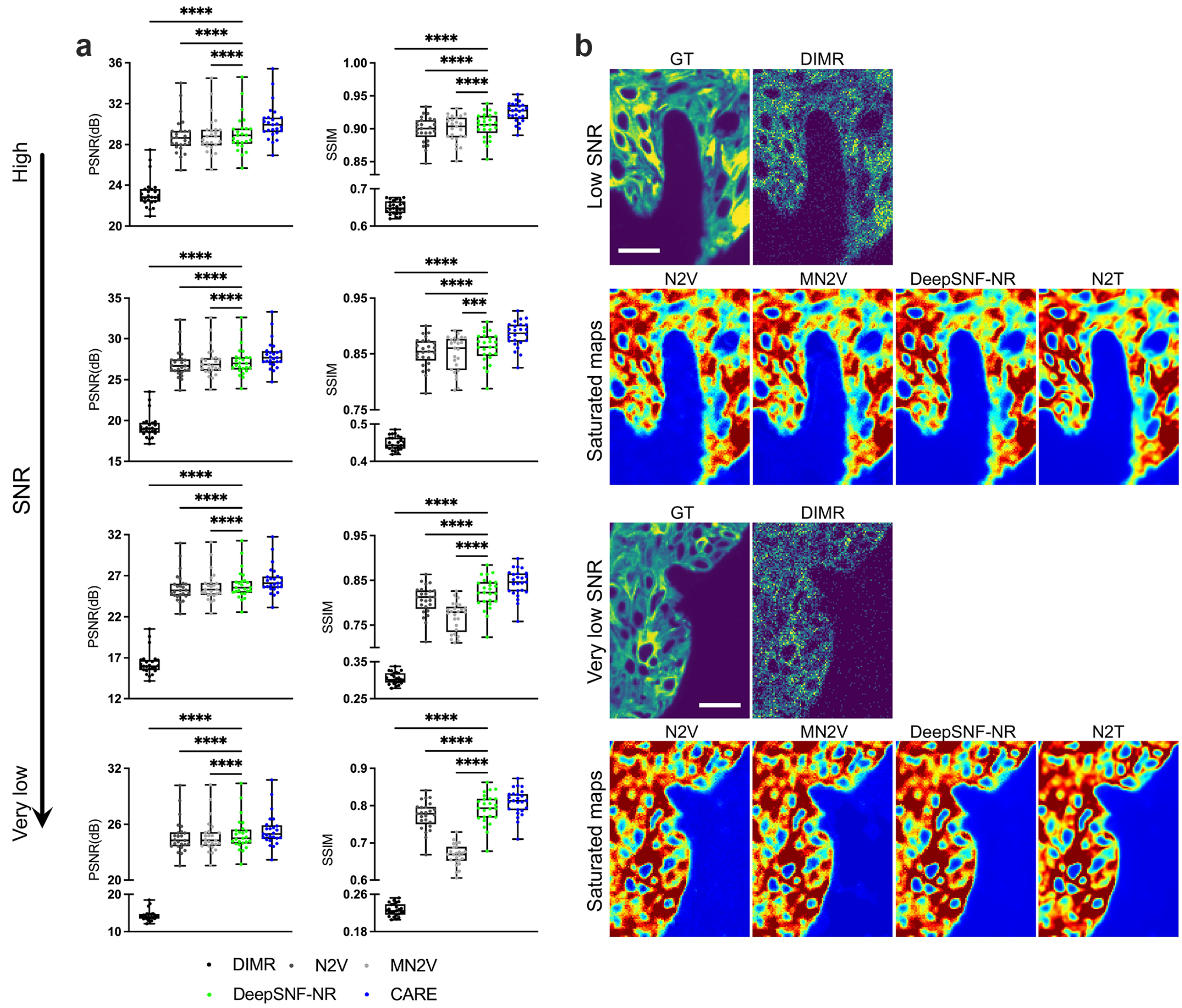

**Supplementary Figure 6.** Comparison of DeepSNF-NR with Noise2Void and modified Noise2Void on the simulated Keratin dataset with 4 noise levels ($N = 25$ per level). (a) PSNR and SSIM evaluation on the three algorithms. (b) Visual inspection of the three algorithms and N2T on denoising a low SNR image. (c) Visual inspection of the three algorithms and N2T on denoising a very low SNR image. Scale bar: (b) 20 $\mu$m, (c) 20 $\mu$m.

In both Supplementary Figs. 5a and 6a, N2T is always the best performer because of the availability of ground truths in training. Nevertheless, in self-supervised learning algorithms, DeepSNF-NR wins over N2V and MN2V on both PSNR and SSIM. The restored image qualities are also reflected in Figs. 5b, c and 6b, c, in which saturated maps are applied to better visualize the differences between the algorithms. The restored images of N2V and MN2V are noiser than the DeepSNF ones, which is more noticeable on low intensity regions . As we analyzed, this is because MSE is a biased estimator for Poisson noise, even with the Anscombe transformation. Interestingly, we observe the SSIM of MN2V is lower than N2V in several Keratin datasets (medium, low and very low SNRs). We infer this results from the ReLU activation of the output layer, which may suffer from the dead neuron problem. In this case, some pixels with very low intensities are always 0 and do not response the back propagation. The 0 value pixels destroy the overall structure and hence lower the SSIM. To conclude, we have verified that our derived I-divergence is a better estimator than MSE on Poisson denoising task, and our derived DeepSNF framework is more capable to restore IMC images than N2V and MN2V.

##### 3.4.2 The effect of Hessian regularization on DeepSNF

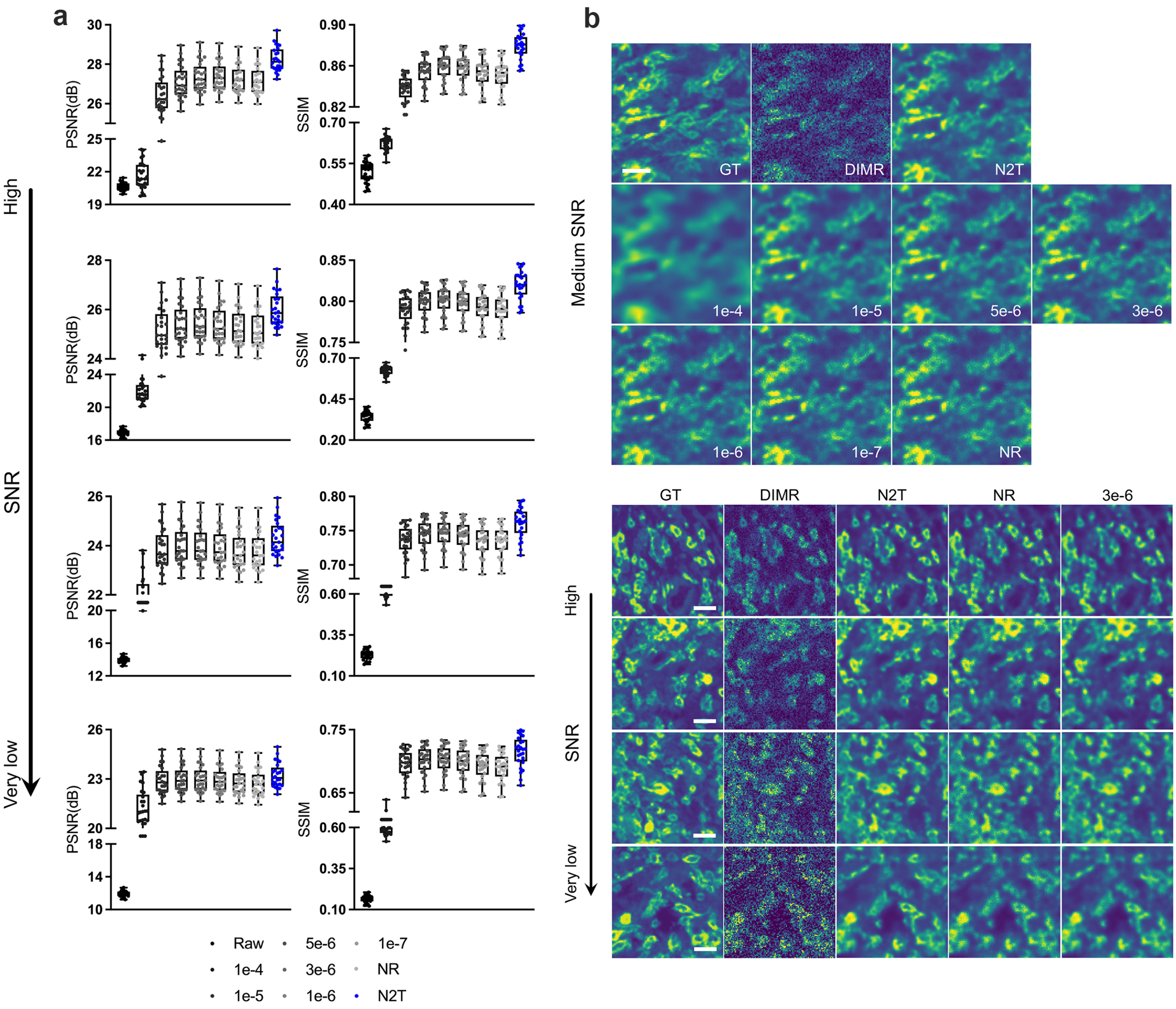

**Supplementary Figure 7.** The effect of Hessian regularization on the simulated CD14 image denoising ($N = 25$ per level). (a) PSNR and SSIM evaluation for DeepSNF with different Hessian regularization parameters $\lambda_{Hessian}$. (b) Visual inspection of DeepSNF with different Hessian regularization parameters $\lambda_{Hessian}$ on a simulated CD14 image with medium SNR. (c) Visual inspection of DeepSNF when $\lambda_{Hessian} = 0$ (NR) and $\lambda_{Hessian}$=3e-6 on simulated CD14 images with different noise levels. Scale bar: (b) 20 $\mu$m, (c) 20 $\mu$m.

Even though DeepSNF-NR is superior than N2V and MN2V on IMC denoising, it still suffers from the discontinuous artifacts and there remains a big gap between the performance of DeepSNF and N2T.

To further enhance DeepSNF, the Hessian regularization is applied in the loss function (26). In particular, $\lambda_{Hessian}$ determines the strength of regularization and thus further affects the denoising performance. To choose a proper $\lambda_{Hessian}$, we trained a series of networks with a range of $\lambda_{Hessian}$ from 1e-4 to 1e-7 and no regularization (NR) on the simulated CD14 and Keratin images. Here, 25 images were used for training and the other 25 for test in each dataset as well.

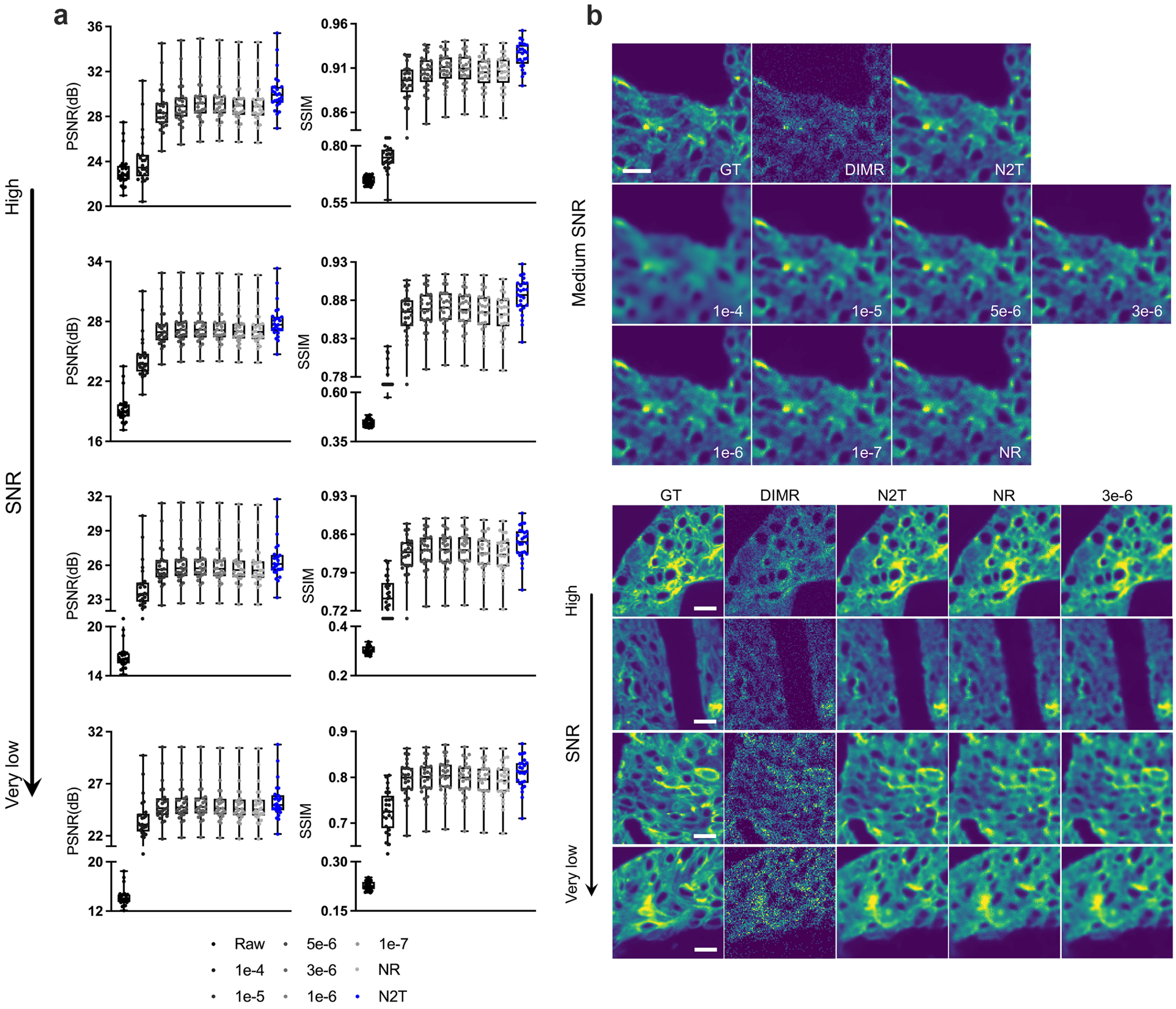

**Supplementary Figure 8.** The effect of Hessian regularization on the simulated Keratin image denoising ($N = 25$ per level). (a) PSNR and SSIM evaluation for DeepSNF with different Hessian regularization parameters $\lambda_{Hessian}$. (b) Visual inspection of DeepSNF with different Hessian regularization parameters $\lambda_{Hessian}$ on a simulated Keratin image with medium SNR. (c) Visual inspection of DeepSNF when $\lambda_{Hessian} = 0$ (NR) and $\lambda_{Hessian}$=3e-6 on simulated Keratin images with different noise levels. Scale bar: (b) 20 $\mu$m, (c) 20 $\mu$m.

The quantitative and qualitative results are presented in Supplementary Figs. 7 and 8. The PSNR and SSIM from both groups (Supplementary Figs. 7a and 8a) indicate Hessian regularization could enhance the performance of DeepSNF with a proper $\lambda_{Hessian}$. Specifically, we found DeepSNF achieves its optimal performance when $\lambda_{Hessian}$ is approximately 3e-6. We then investigated the qualities of the restored images from the medium SNR datasets (Supplementary Figs. 7b and 8b). The larger $\lambda_{Hessian}$, the stronger regularization will be. Therefore, the images look more blurry when $\lambda_{Hessian}$ = 1e-4 and noisier with no regularization. With $\lambda_{Hessian}$ = 3e-6, the restored images achieve their best qualities. Subsequently, by observing the denoised images from different noise levels (Supplementary Figs. 7c and 8c), the Hessian regularization effectively modifies the discontinuous artifacts in the images without regularization. We conclude that the Hessian regularization effectively improves the performance of DeepSNF, both quantitatively and qualitatively, even boost the performance of DeepSNF close to that of N2T. As a consequence, we set DeepSNF with $\lambda_{Hessian}$ = 3e-6 for the subsequent experiments due to its optimal accuracy.

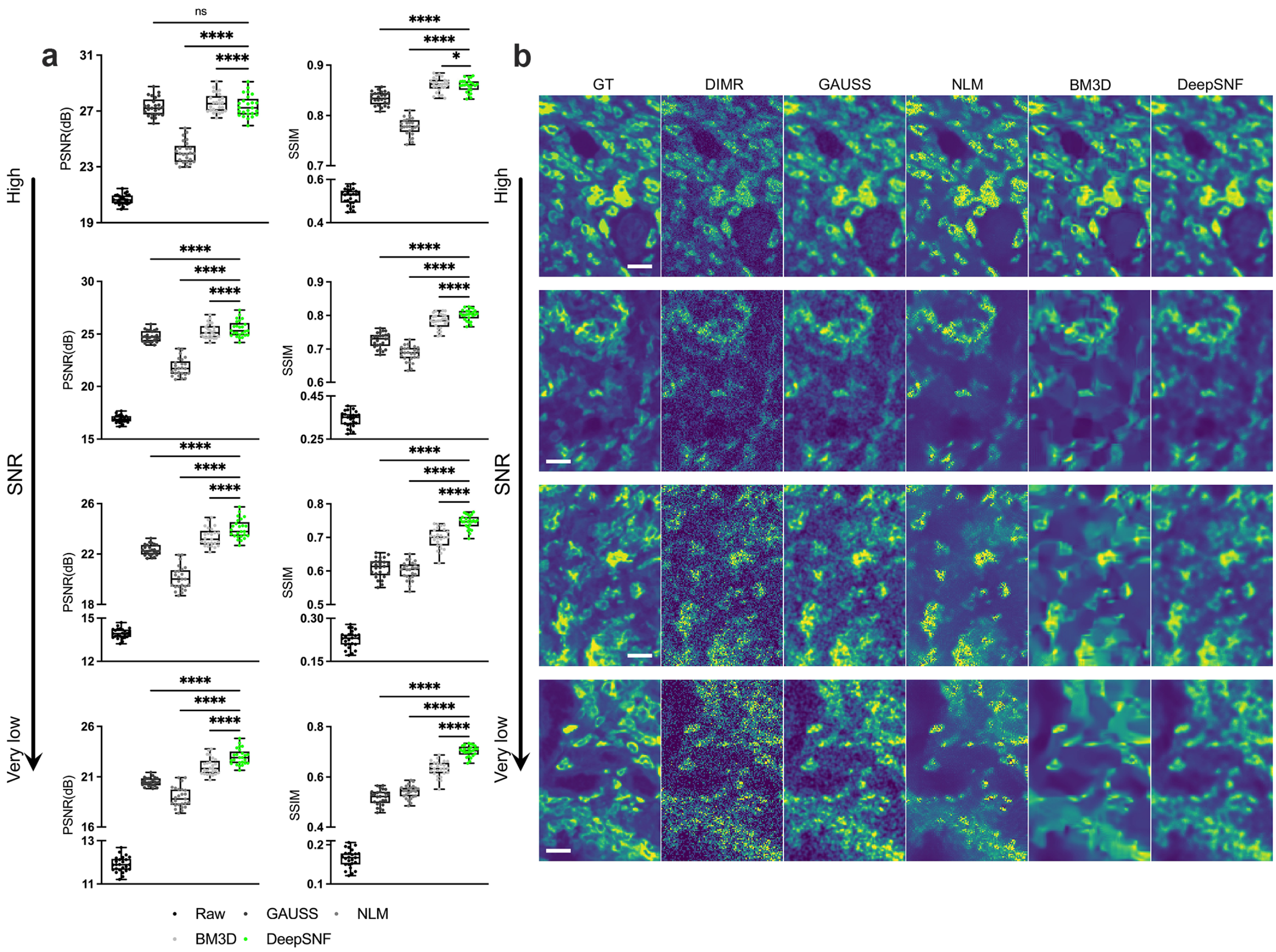

**Supplementary Figure 9.** Comparison of DeepSNF with GAUSS, NLM and BM3D algorithms on the simulated CD14 dataset with 4 noise levels ($N = 25$ per level). (a) PSNR and SSIM evaluation on the algorithms. (b) Visual inspection of the algorithms on denoising the images with different noise levels. Scale bar: 20 $\mu$m.

##### 3.4.3 Compare DeepSNF with traditonal denoising methods

We are also interested in comparing DeepSNF with traditional statistical denoising methods, including Gaussian filter (GAUSS), NLM and BM3D algorithms. These three algorithms along with DeepSNF were benchmarked on the simulated CD14 and Keratin datasets. Notably, they were only applied on the 25 test images of DeepSNF. The results are presented in Supplementary Figs. 9 and 10.

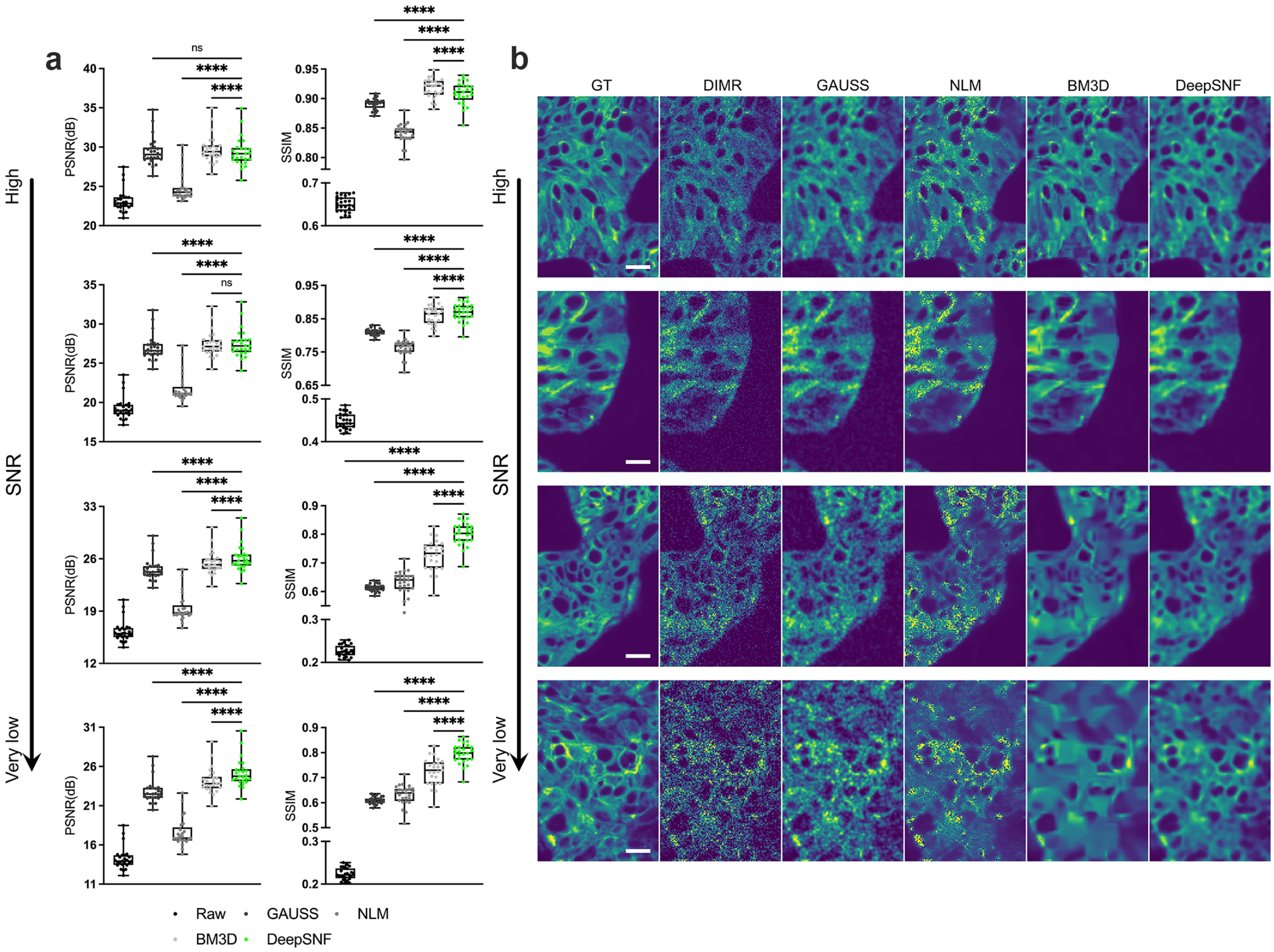

**Supplementary Figure 10.** Comparison of DeepSNF with GAUSS, NLM and BM3D algorithms on the simulated Keratin dataset with 4 noise levels ($N = 25$ per level). (a) PSNR and SSIM evaluation on the algorithms. (b) Visual inspection of the algorithms on denoising the images with different noise levels. Scale bar: 20 $\mu$m.

From the quantitative evaluation results (Supplementary Figs. 9a and 10a), DeepSNF is almost always better than the other algorithms on PSNR and SSIM, except on high SNR images. This is also in accordance with the visual inspection (Supplementary Figs. 9b and 10b). For high SNR images, BM3D performs slightly better than DeepSNF. Nevertheless, as SNR becomes lower, BM3D tends to over-smooth the restored images and lots of details are lost. In fact, the SNR of IMC images are always not high because of the limited ion counts. As a result, BM3D is not fit for IMC denoising, even though its PSNR and SSIM are the closest to those of DeepSNF. GAUSS and NLM are even not as good as BM3D. Particularly, GAUSS algorithm can be regarded as a low-pass filter and only the high frequency noises are filtered. Similar to BM3D, NLM algorithm also blurs the restored images. However, it still leaves lots of noisy regions and its denoising ability is even weaker than GAUSS. To conclude, traditional statistical denoising algorithms are not good at IMC denoising, and deep learning-based algorithms such as DeepSNF should be preferred.

### Supplementary Figures

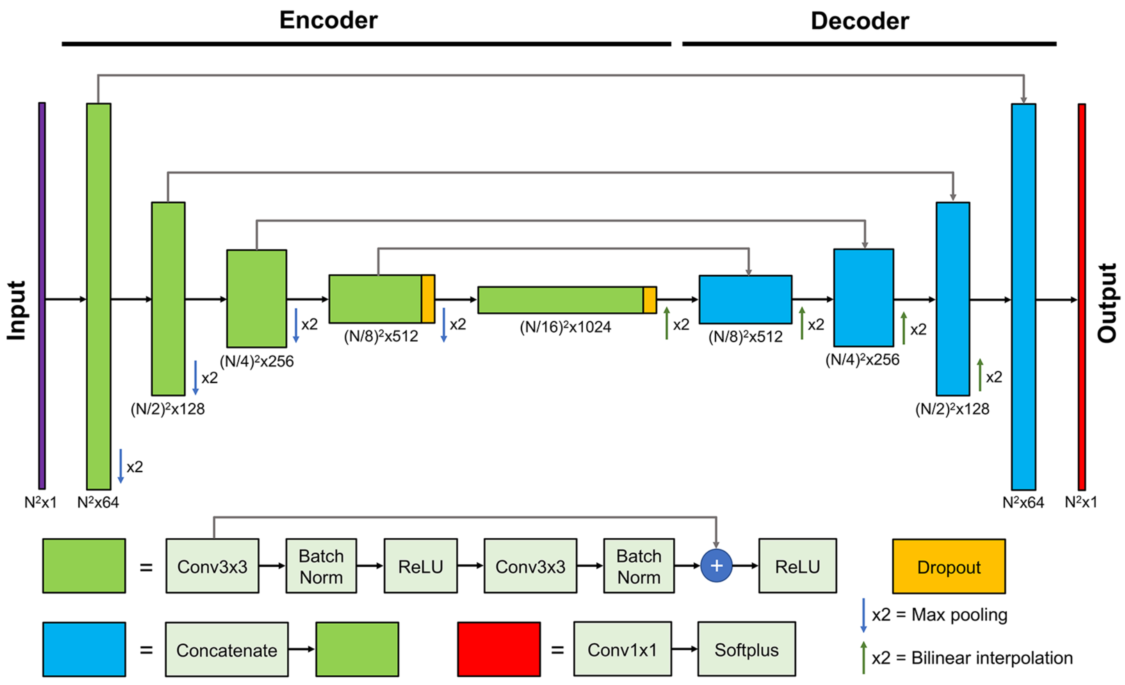

**Supplementary Figure 11.** The proposed convolutional neural network structure. The overall structure of the network follows the U-Net architecture modified with Res-block modules. The input and the output images share the same size. Starting with the noisy image, the encoder path gradually condenses the spatial information into high-level feature maps with growing depths (size marked by the number in the bottom of each block); the decoder path reverses the process by recombining the information into feature maps with gradually increased lateral details. The information in adjacent feature maps transfers by convolving with $3 \times 3$ convolutional filters. The down-sampling is done by $2 \times 2$ max-pooling operation and the up-sampling is done by $2 \times 2$ up-sampling operation. Res-blocks are applied to facilitate efficient training. Each Res-block contains convolution layer (Conv), batch normalization (Batch Norm) and the rectified linear unit (ReLU) nonlinear activation. We also added dropout layers with 0.5 dropout rate after the central two res-blocks to mitigate overfitting. Skip connections are added to tunnel the high-frequency information from shallower layers to deeper layers with the same spatial scales. We use the softplus activation function $\log(1 + \exp(x))$ in the final layer to restrict the final prediction non-negative.

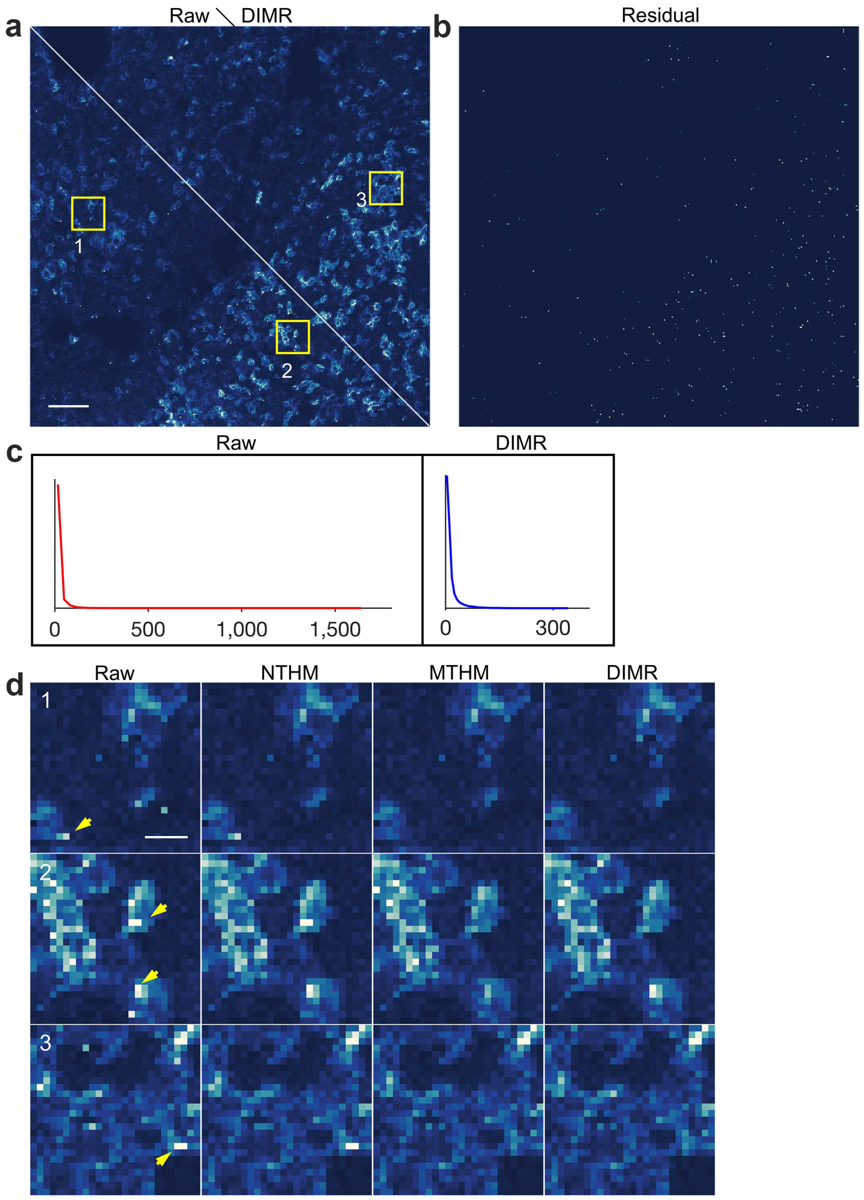

**Supplementary Figure 12.** DIMR algorithm enables adaptive hot pixel removal of CD235a in human bone marrow IMC dataset. (a) Comparison of the raw and DIMR-processed images, where the lower left and upper rights parts correspond to the raw and DIMR-processed images, respectively. (b) Difference between the raw and DIMR-processed images, in which the residual pixels correspond to the detected hot pixels. (c) The corresponding histograms of raw and DIMR-processed images in (a). (d) Comparisons between the raw, NTHM, MTHM and DIMR-processed images. (d1)–(d3) correspond to the sub-regions labeled from 1 to 3 in (a). Scale bar: (a) 40 $\mu$m, (d) 6 $\mu$m.

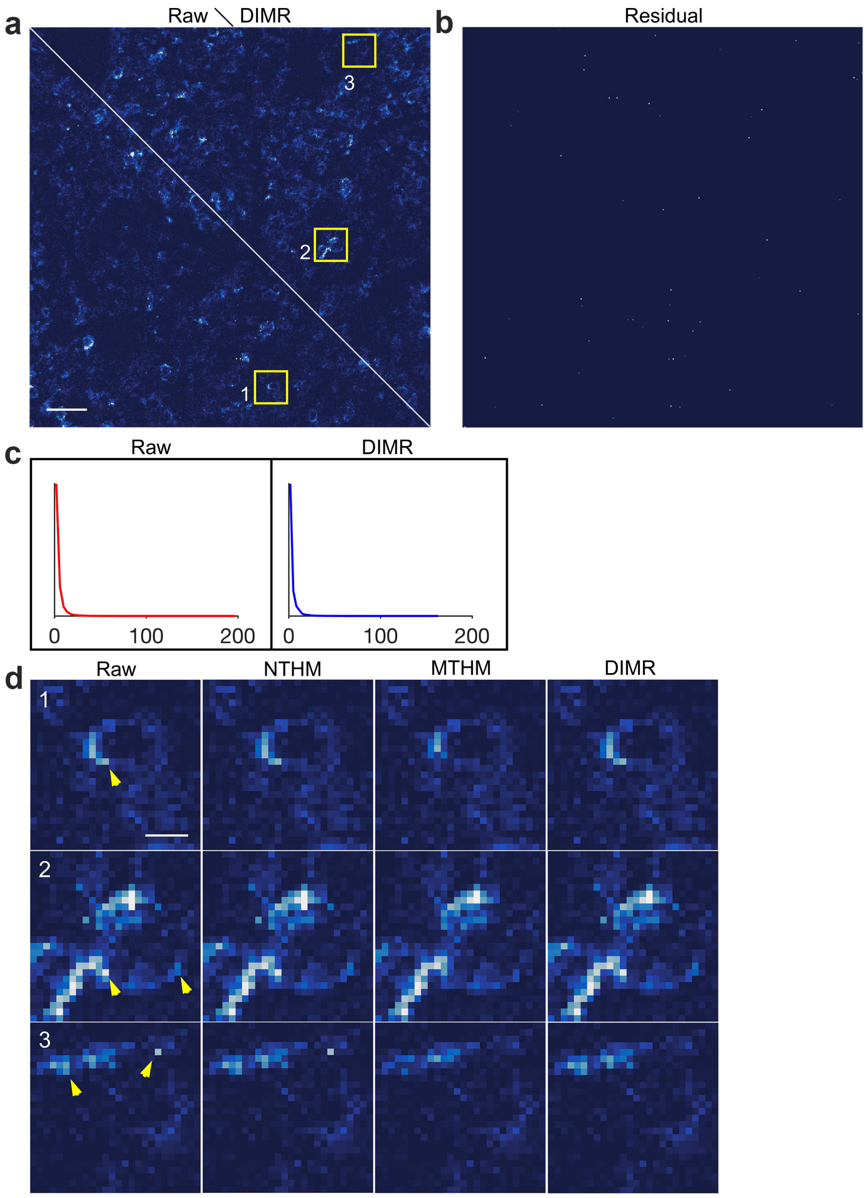

**Supplementary Figure 13.** DIMR algorithm enables adaptive hot pixel removal of MPO in human bone marrow IMC dataset. (a) Comparison of the raw and DIMR-processed images, where the lower left and upper rights parts correspond to the raw and DIMR-processed images, respectively. (b) Difference between the raw and DIMR-processed images, in which the residual pixels correspond to the detected hot pixels. (c) The corresponding histograms of raw and DIMR-processed images in (a). (d) Comparisons between the raw, NTHM, MTHM and DIMR-processed images. (d1)–(d3) correspond to the sub-regions labeled from 1 to 3 in (a). Scale bar: (a) 40 $\mu$m, (d) 6 $\mu$m.

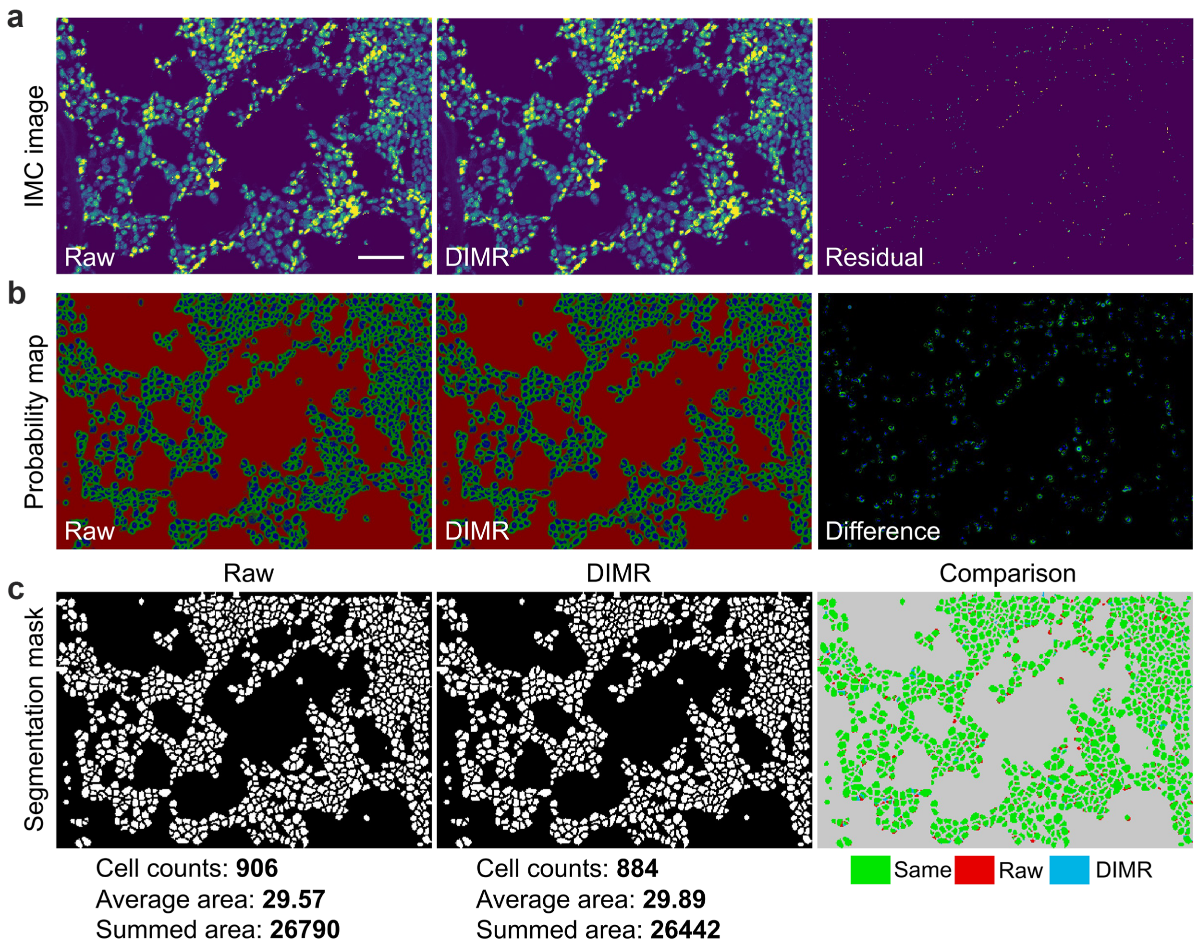

**Supplementary Figure 14.** DIMR slightly enhances single cell segmentation. (a) Comparison between the raw and DIMR-processed DNA images. (b) The probability maps of the raw and DIMR-processed images generated by Ilastik software, and their difference map (blue: nuclei, green: cytoplasm, and red:background). (c) The segmentation masks of the raw and DIMR-processed images generated by Cell Profiler software, and their overlaid comparison map. To account for the impact of hot pixels in raw image, the segmented masks with area smaller than 5 have been removed in both segmentation masks. By comparing the figures in the third column, the different segmented masks between raw and DIMR images are mainly caused by the hot pixels. Even with a cell size threshold, the hot pixels can still split cells, and falsely expand normal cell borders. Correspondingly, the raw image segmented a little more cells than that of DIMR (906 to 994). At the same time, the average area of cells from the raw image is a little smaller (29.57 to 29.89), but the summed area is slightly larger (26790 to 26442). Scale bar: 47 $\mu$m.

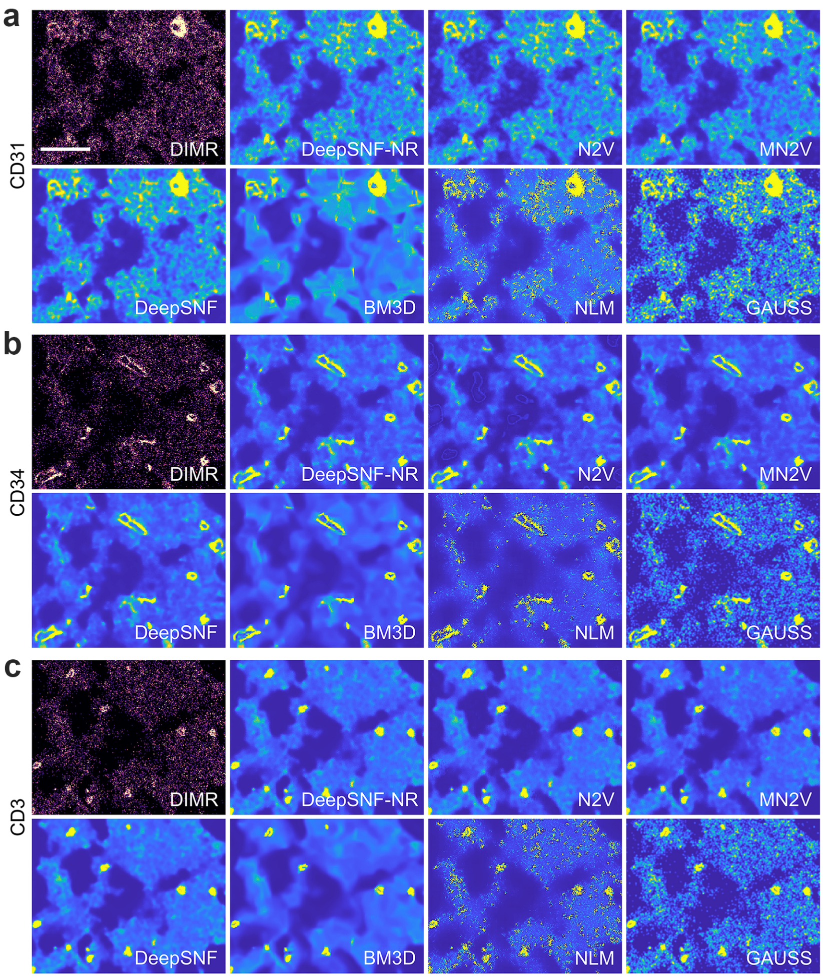

**Supplementary Figure 15.** Visual inspection of DeepSNF and other statistics-based denoising algorithms on denoising (a) CD31, (b) CD34, (c) CD3-labeled IMC images. Scale bar: 60 $\mu$m.

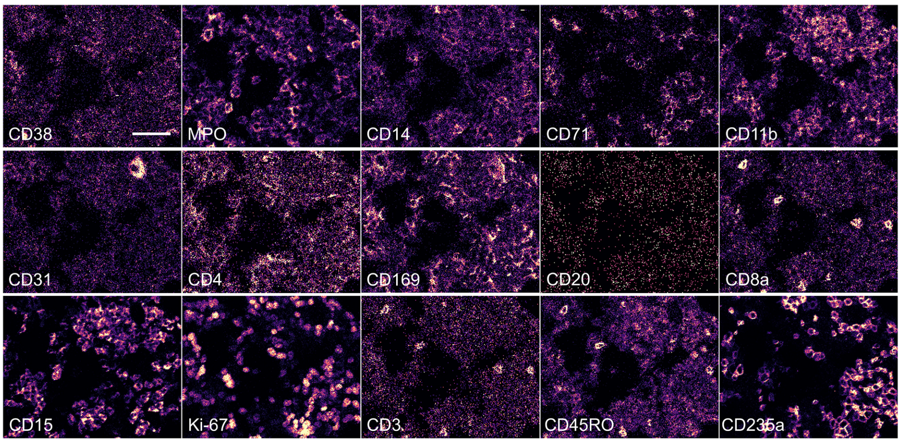

**Supplementary Figure 16.** The raw IMC images corresponding to Fig. 1f. Scale bar: 48 $\mu$m.

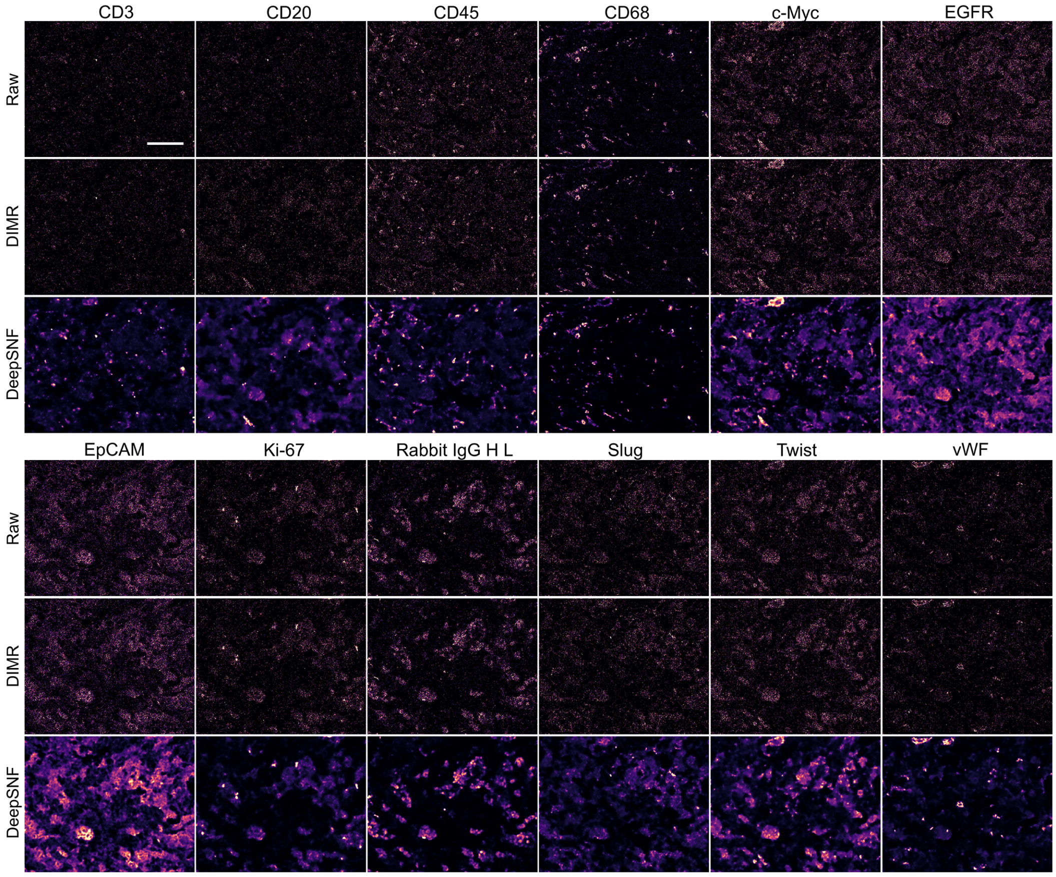

**Supplementary Figure 17.** IMC-Denoise enhances the human breast cancer IMC dataset. Scale bar: 100 $\mu$m.

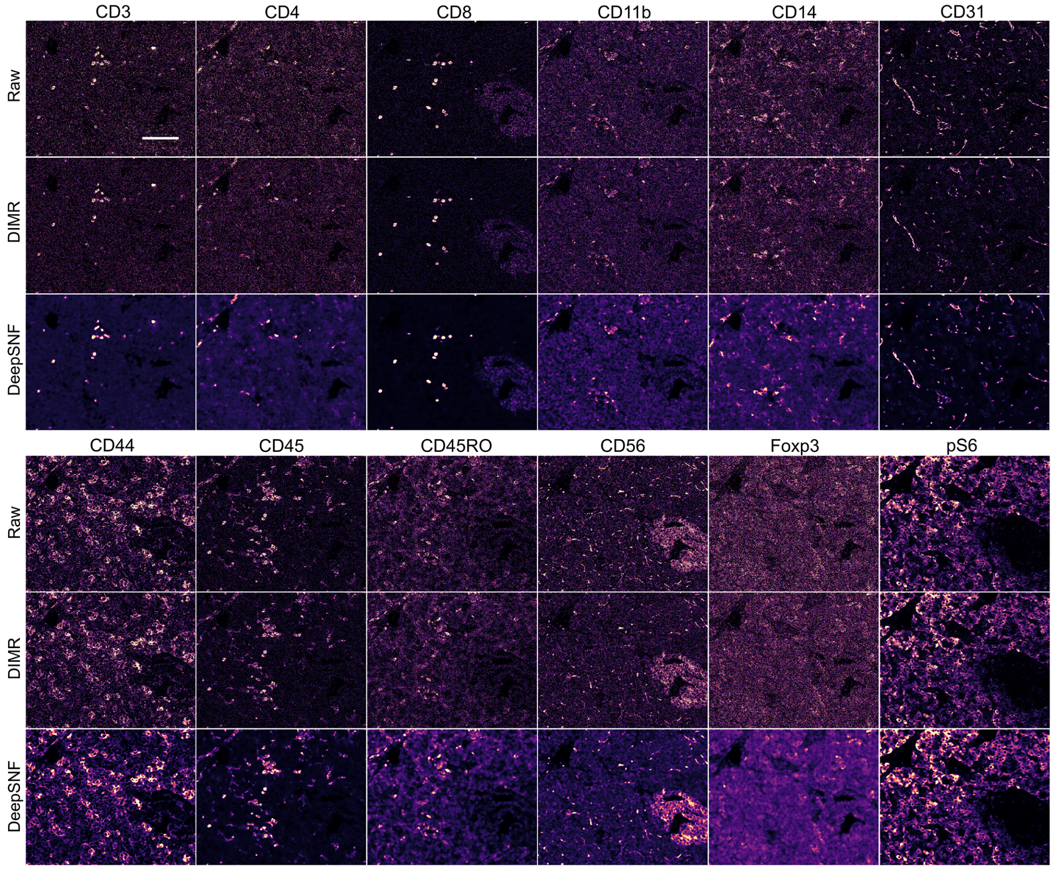

**Supplementary Figure 18.** IMC-Denoise enhances the human pancreatic cancer IMC dataset. Scale bar: 100 $\mu$m.

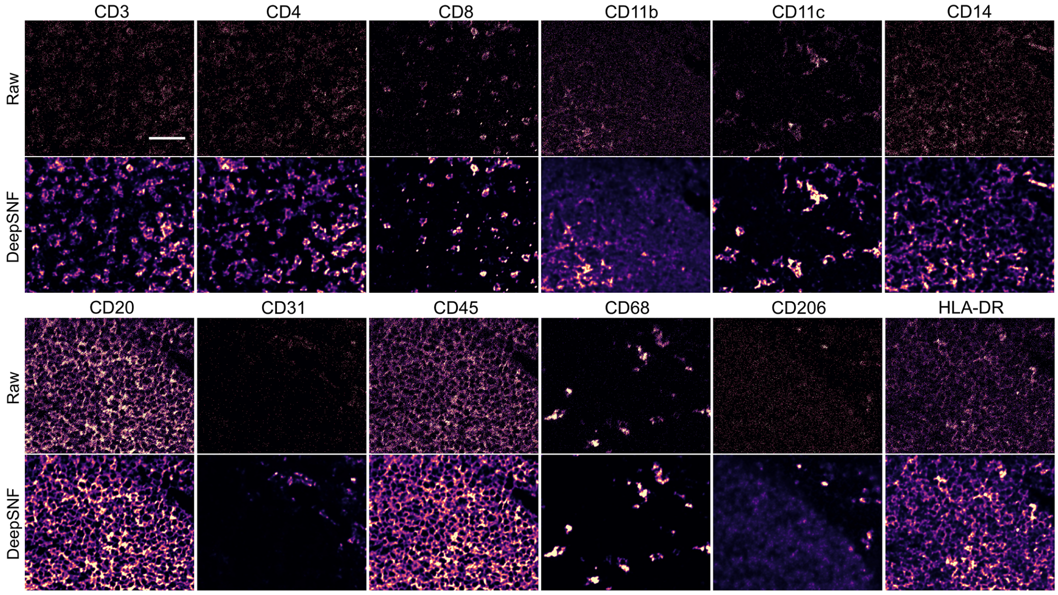

**Supplementary Figure 19.** IMC-Denoise enhances the MIBI dataset. Here, only DeepSNF was applied to the MIBI images because no hot pixels were observed. The channel crosstalk and hot clusters observed in MIBI images could be removed by the MAUI software package afterwards. Scale bar: 25 $\mu$m.

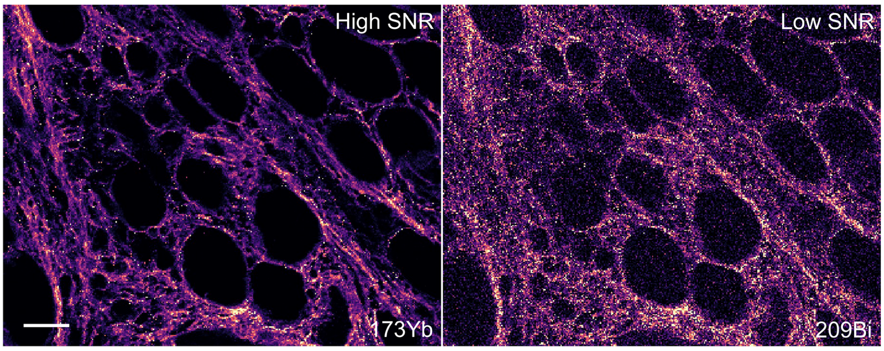

**Supplementary Figure 20.** The raw IMC images corresponding to Fig. 1g. Scale bar: 37 $\mu$m.

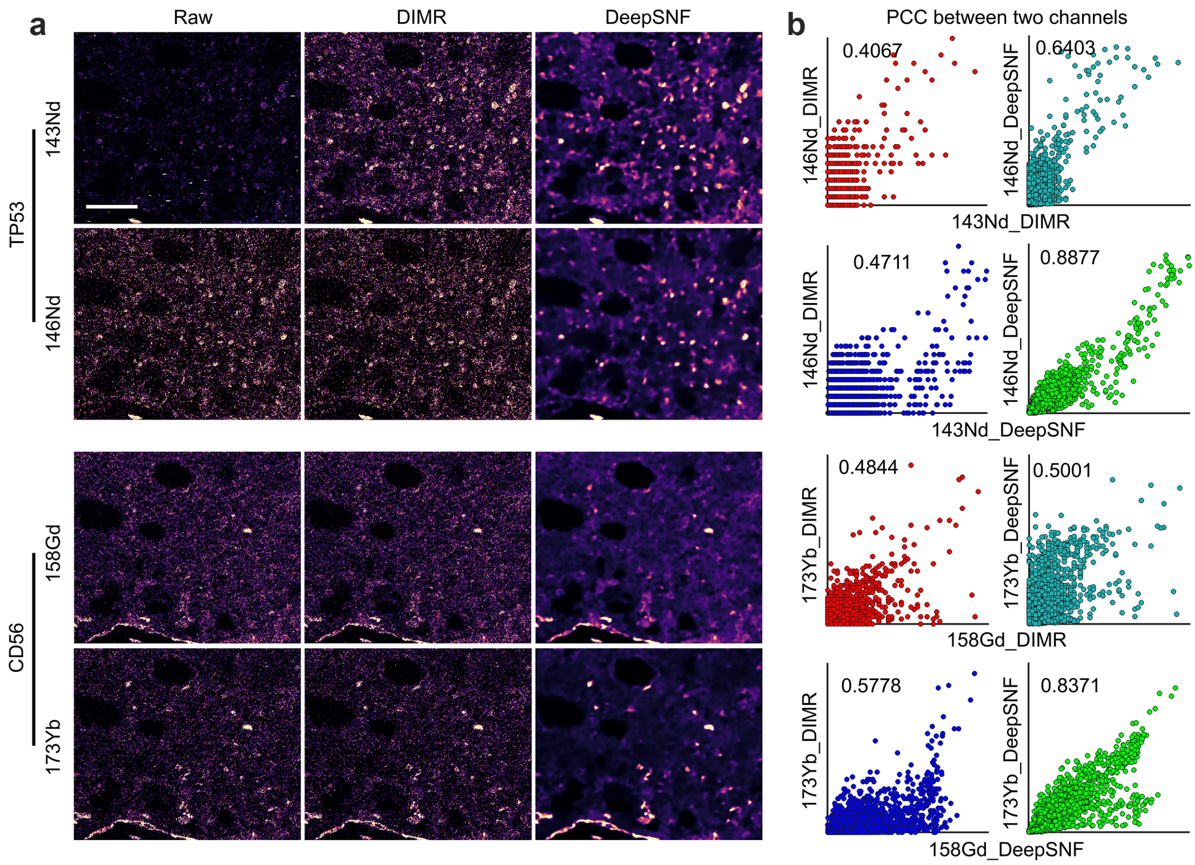

**Supplementary Figure 21.** IMC-Denoise enhances both IMC image quality and the Pearson correlations of IMC images labeled with the same markers. (a) A tissue was stained by 143Nd, 146Nd conjugated TP53, and 158Gd, 173Yb conjugated CD56, respectively, with different SNRs. The IMC images were firstly processed by DIMR to remove hot pixels. Then DeepSNF was employed to improve the image quality of all the images, because the qualities of the higher SNR images are still sub-optimal. (b) After DeepSNF processing, the Pearson correlation coefficients (PCC) improved, in which those of the double DeepSNF-processed images are the highest. Notably, the DeepSNF trained by the CD3 images from the human bone marrow dataset (Supplementary Tables 5 and 8) was used to denoise the DIMR-processed TP53 and CD56 images, due to their highly similar features and the lacking of sufficient TP53 and CD56 training sets. Scale bar: 75 $\mu$m.

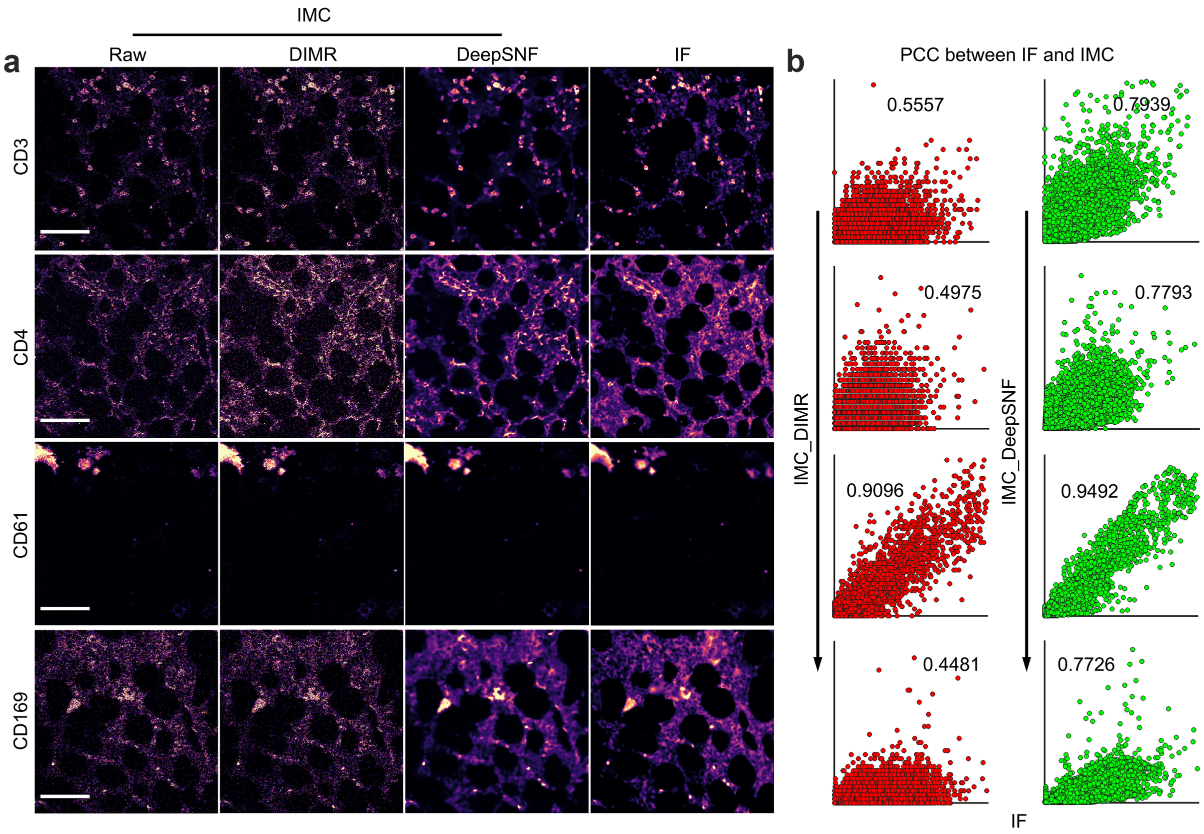

**Supplementary Figure 22.** IMC-Denoise enhances both IMC image quality and the Pearson correlations between IMC and IF images. (a) The same tissues were stained with CD3, CD4, CD61 and CD169 by IMC and IF, respectively. The low SNR IMC images were processed by DIMR to remove hot pixels and then by DeepSNF to improve image quality. (b) After DeepSNF processing, the PCC between IMC and IF improved, indicating DeepSNF is able to improve the IMC image quality. Scale bar: CD3: 98 $\mu$m. CD4: 110 $\mu$m. CD61: 69 $\mu$m. CD169: 87 $\mu$m.

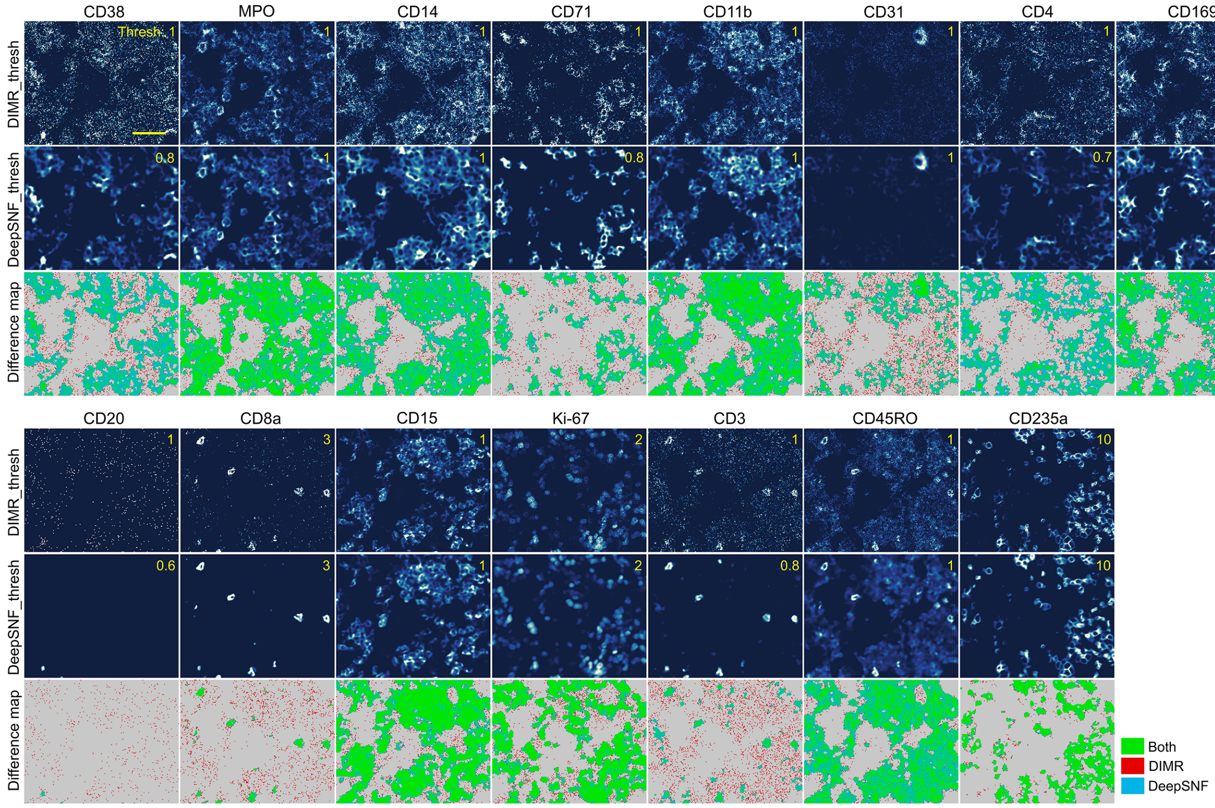

**Supplementary Figure 23.** DeepSNF eliminates background noise of the IMC images from human bone marrow dataset. The raw IMC images were processed by DIMR to remove hot pixels and then by DeepSNF to account for shot noise (Fig. 1f). Thresholds (upper right corner in every image) were selected to remove the background noise of the DIMR and DeepSNF-processed images. The signal masks of DIMR and DeepSNF-processed images were overlaid to compare the difference of background removal. Scale bar: 48 $\mu$m.

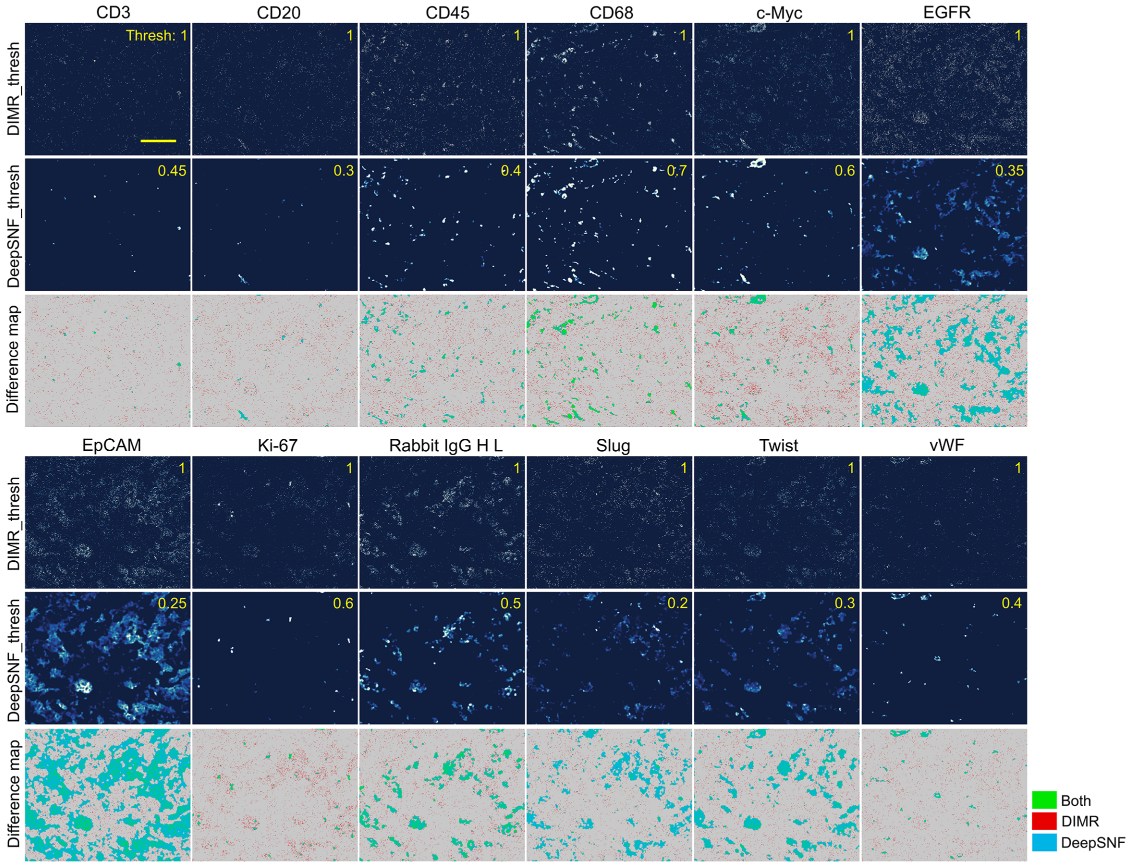

**Supplementary Figure 24.** DeepSNF eliminates background noise of the IMC images from human breast cancer dataset (Supplementary Fig. 17). Thresholds (upper right corner in every image) were selected to remove the background noise of the DIMR and DeepSNF-processed images. The signal masks of DIMR and DeepSNF-processed images were overlaid to compare the difference of background removal. Scale bar: 100 $\mu$m.

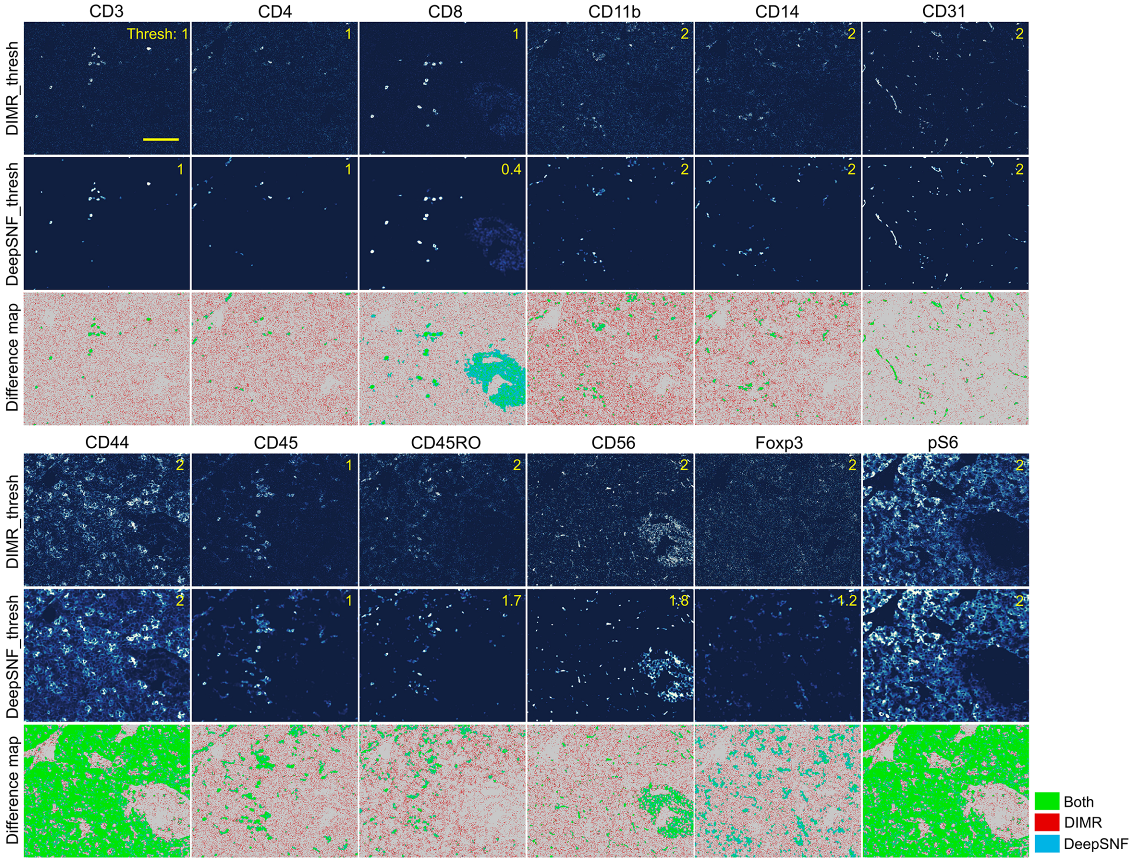

**Supplementary Figure 25.** DeepSNF eliminates background noise of the IMC images from human pancreas cancer dataset (Supplementary Fig. 18). Thresholds (upper right corner in every image) were selected to remove the background noise of the DIMR and DeepSNF-processed images. The signal masks of DIMR and DeepSNF-processed images were overlaid to compare the difference of background removal. Scale bar: 100 $\mu$m.

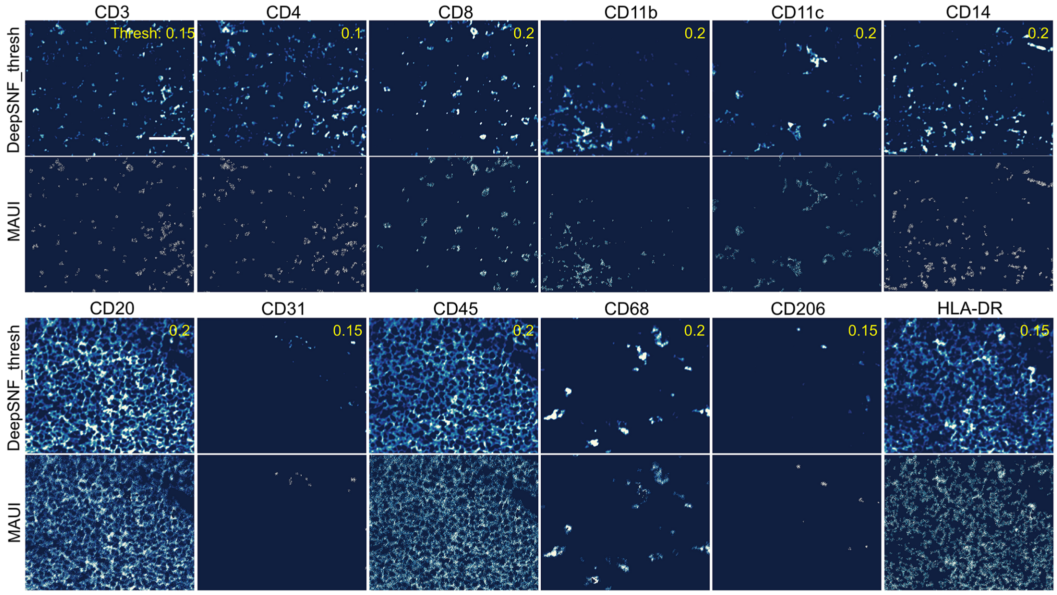

**Supplementary Figure 26.** DeepSNF eliminates background noise of the MIBI images (Supplementary Fig. 19). Thresholds (upper right corner in every image) were selected to remove the background noise of the DeepSNF-processed images. The MAUI software package was also used to remove the background noise of the MIBI images. Visual inspections indicate the performance of DeepSNF is comparable to that of the MAUI software package. Scale bar: 25 $\mu$m.

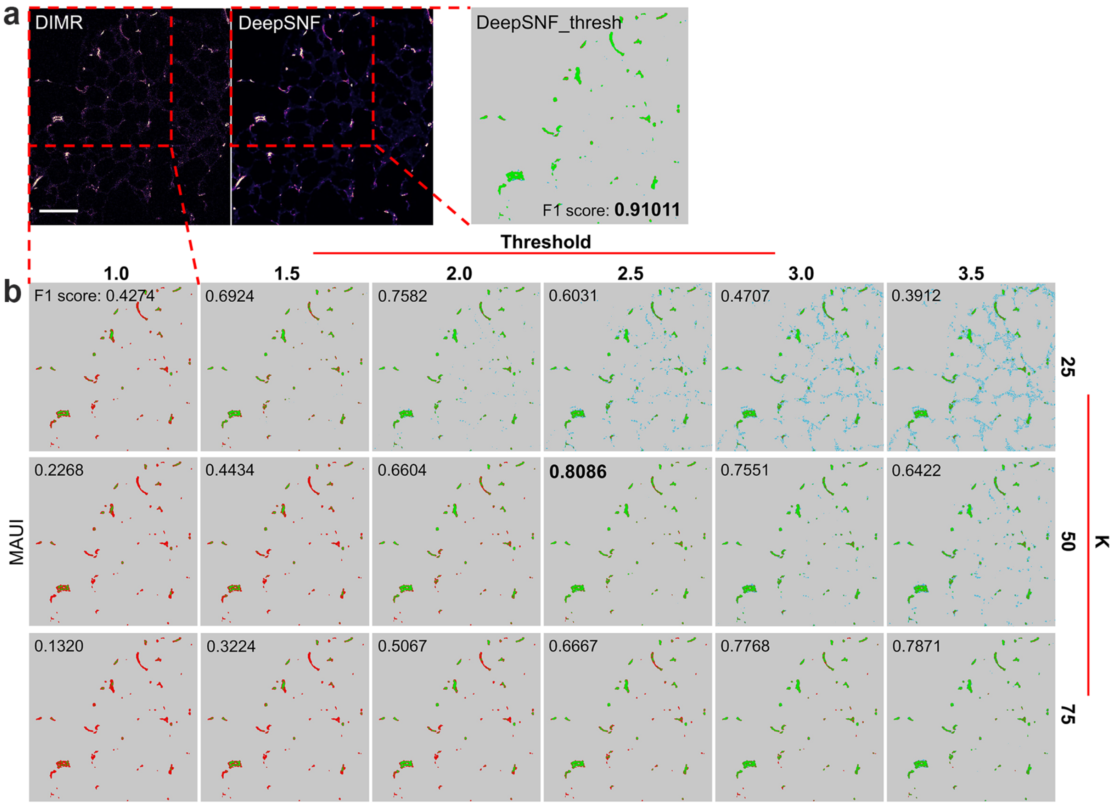

**Supplementary Figure 27.** DeepSNF performs better than MAUI on DIMR data to filter background noise. (a) A DIMR and DeepSNF-processed IMC image labeled with CD34. The DeepSNF-processed image was binarized by the threshold value 1 and then overlaid with manual annotated ground truth. (b) The DIMR-processed image was processed by the MAUI software package with a wide range of parameters to select the best background noise removal result and also overlaid with the manual annotated ground truth. The F1 score of the DeepSNF_thresh (lower right corner) result is always higher than that of the MAUI results (upper left corner in every image), indicating DeepSNF is better than MAUI on DIMR data in terms of background noise removal. Scale bar: 96 $\mu$m.

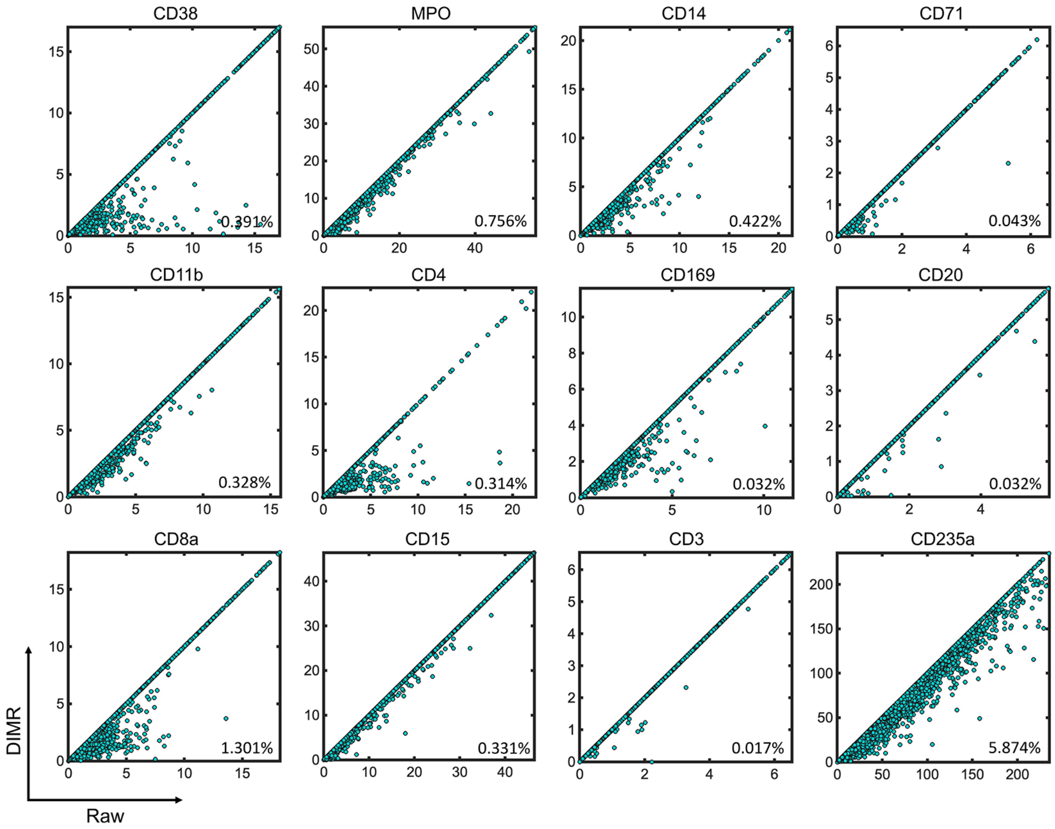

**Supplementary Figure 28.** The impact of DIMR on single cell data. Each sub-figure represents the one-on-one relationship between the raw and DIMR data of a particular marker in single cell scale. The bottom right value in each sub-figure represents the percentage of the difference between the raw and DIMR data.

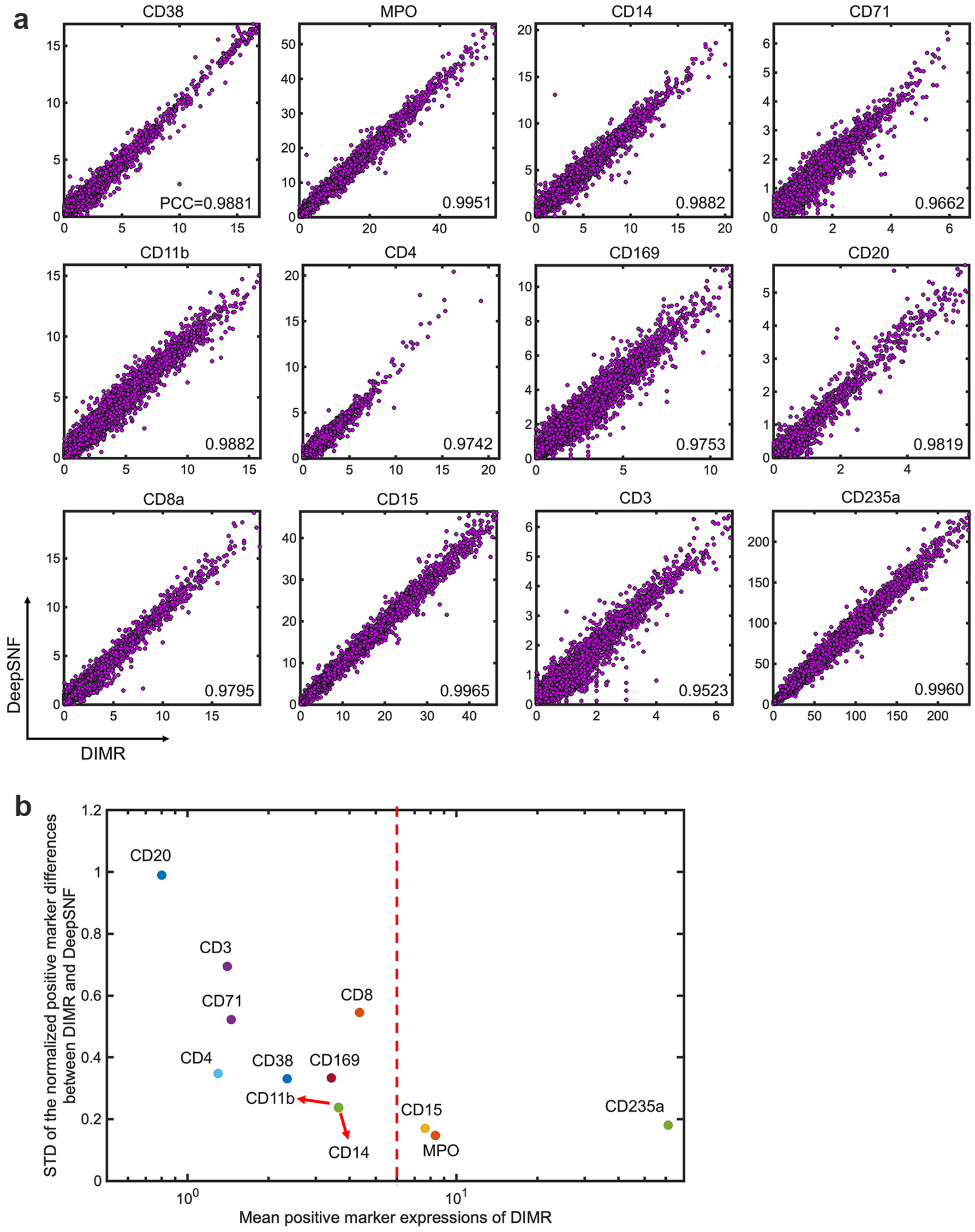

**Supplementary Figure 29.** The impact of DeepSNF on single cell data. (a) Each sub-figure represents the one-on-one relationship between the DIMR and DeepSNF data of a particular marker in single cell scale. The bottom right value in each sub-figure represents the PCC between the DIMR and DeepSNF data. (b) Since the DIMR and DeepSNF data are highly correlated, the standard deviation (STD) of the normalized positive marker differences between DIMR and DeepSNF are utilized to evaluate the impact of DeepSNF on single cell data. For almost all the markers, the larger the mean positive marker expressions, the smaller the STD will be, and then the lighter the impact of DeepSNF will be. This agrees with the fact that the larger the ion count is, the lower the shot noise level will be.

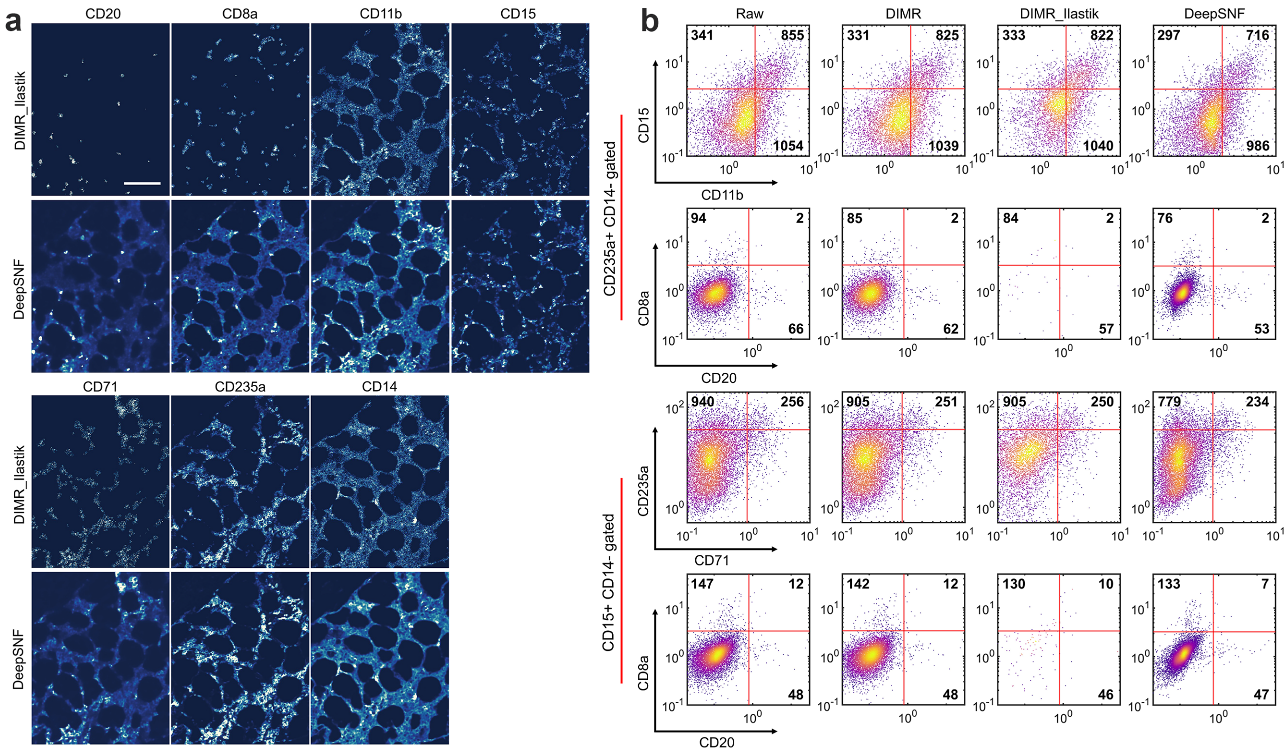

**Supplementary Figure 30.** DeepSNF performs better than Ilastik-based semi-automated denoising approach. (a) Visual inspection of DeepSNF and DIMR_Ilastik denoising results on the IMC images labeled with different markers. (b) Evaluations of denoising algorithms with manual gating strategies on single cell data. We evaluated the results based on the fact that: (1) Myeloid (CD11b and CD15) and erythroid (CD71 and CD235a) markers should be exclusive with each other; (2) CD8a and CD20 should be negative on both myeloid and erythroid cells. Basd on these prior knowledge, DIMR slightly enhances the single cell analysis than the raw data, while DeepSNF further enhances the DIMR results, and overall performs better than DIMR_Ilastik-processed data. Scale bar: 120 $\mu$m.

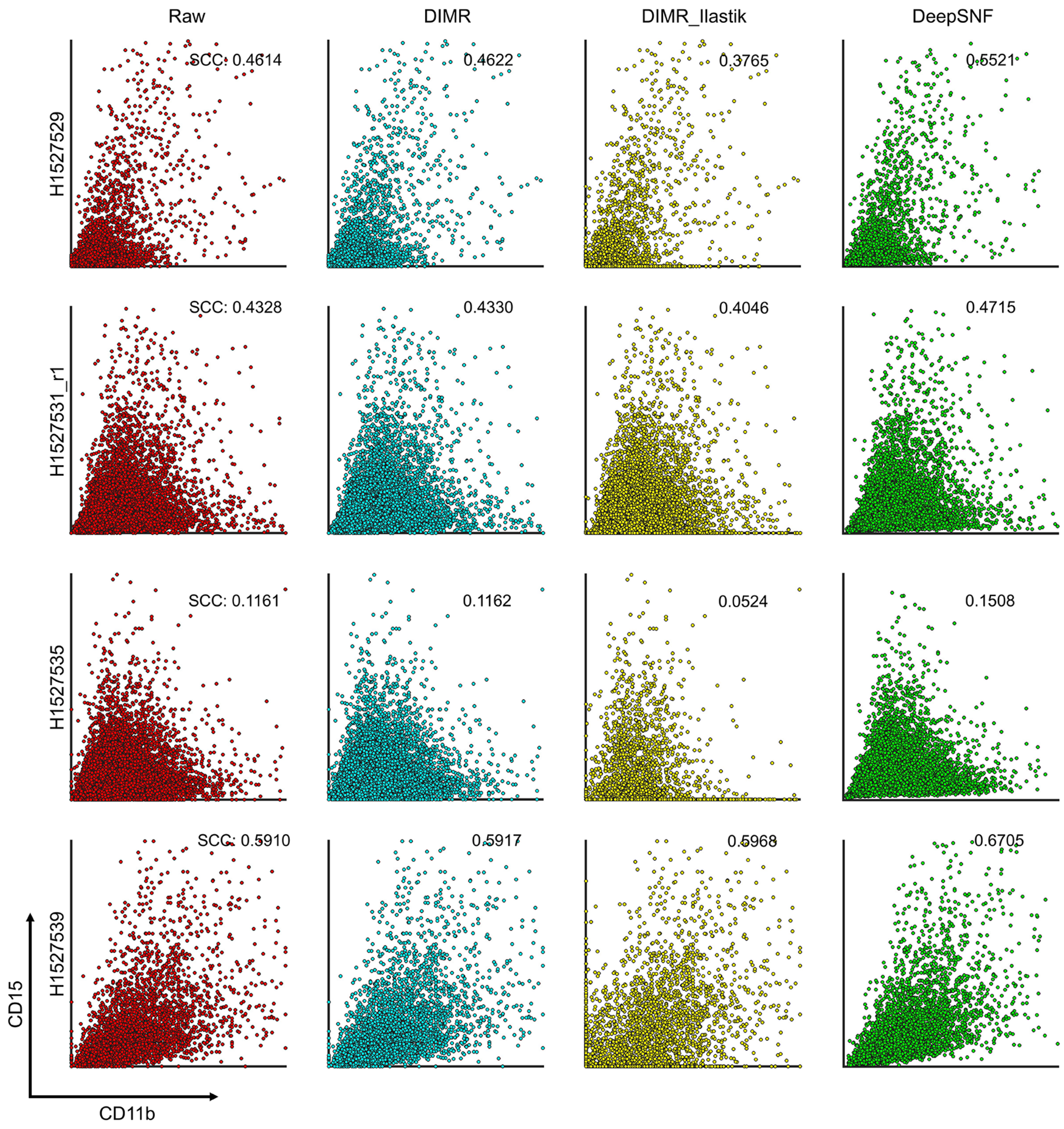

**Supplementary Figure 31.** Cell-to-cell scatter plots between CD11b and CD15 per slide for SCC comparisons of the raw, DIMR, DIMR_Ilastik and DeepSNF-processed IMC images. Particularly, Slides H1527539 and H1527535 correspond to high and low correlation examples, respectively. Slides H1527529 and H1527531_r1 correspond to moderate correlation examples.

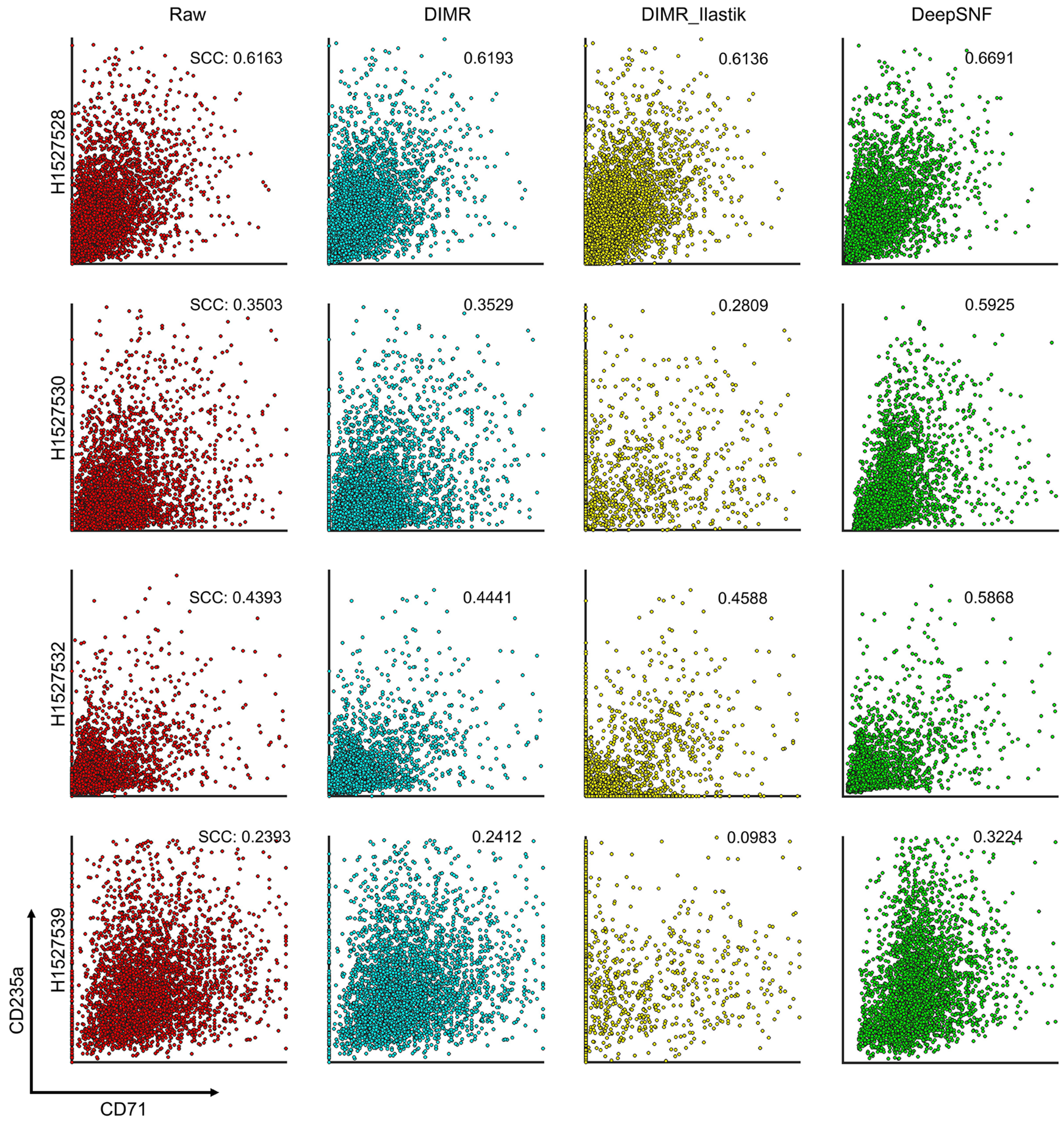

**Supplementary Figure 32.** Cell-to-cell scatter plots between CD71 and CD235a per slide for SCC comparisons of the raw, DIMR, DIMR_Ilastik and DeepSNF-processed IMC images. Particularly, Slides H1527528 and H1527539 correspond to high and low correlation examples, respectively. Slides H1527530 and H1527532 correspond to moderate correlation examples.

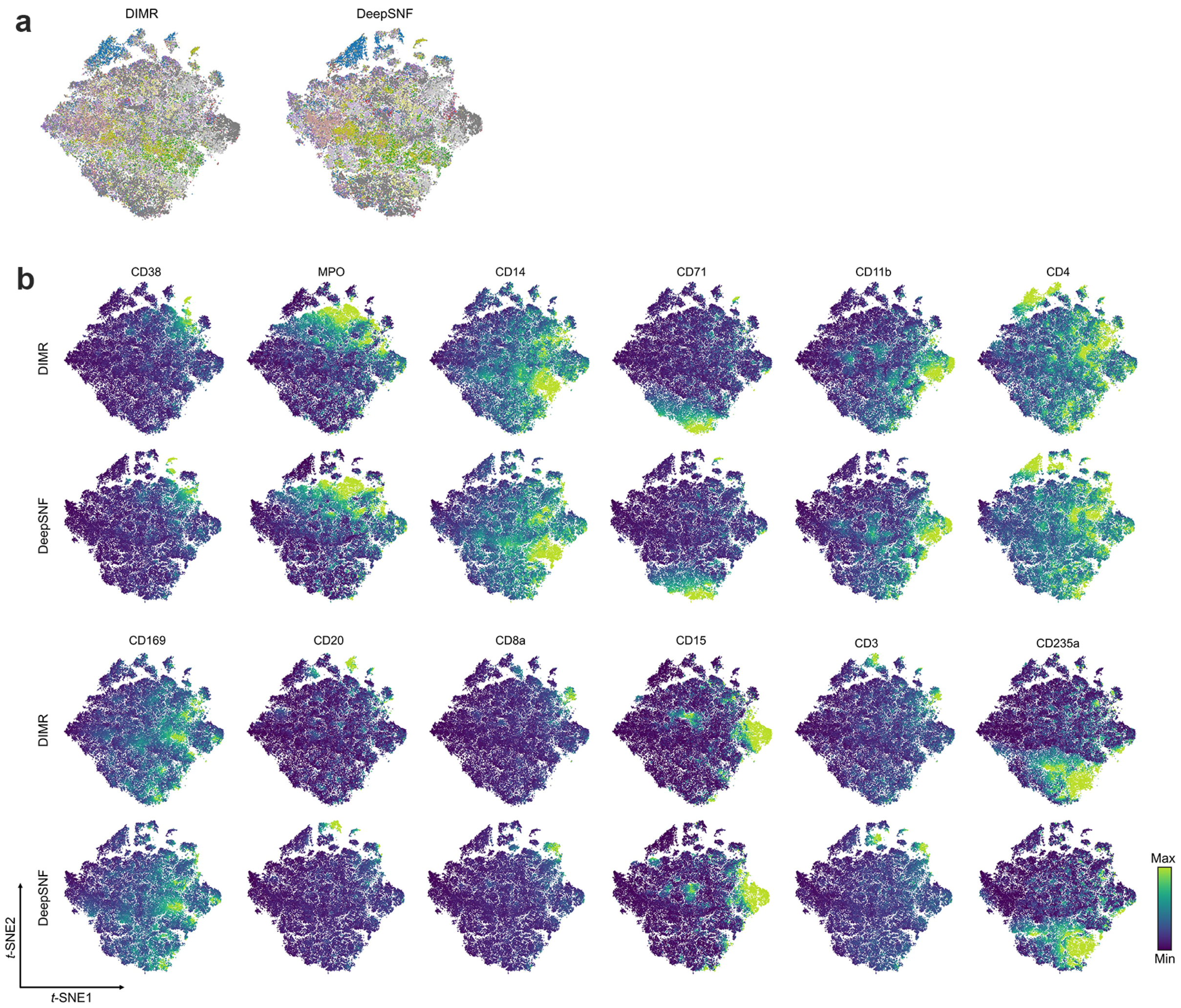

**Supplementary Figure 33.** *t*-SNE plots of the single cell data from the human bone marrow IMC dataset. (a) *t*-SNE plots colored by the cells from different tissues. (b) *t*-SNE plots colored by the single cell marker expressions of DIMR and DeepSNF, respectively.

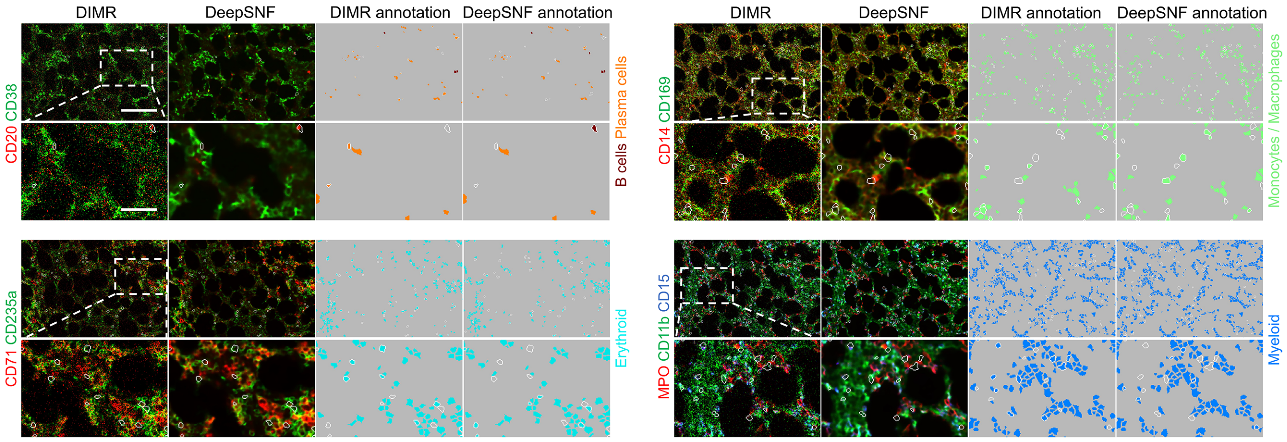

**Supplementary Figure 34.** Comparisons of DIMR and DeepSNF-processed IMC images labeled with different cell markers, and the corresponding cell annotation results (Fig. 3(c)). The bottom row corresponds to the white dashed box regions in the top row images. The white contours represent the different phenotyping results between DIMR and DeepSNF. Scale bar: Top: 145 $\mu$m, bottom: 50 $\mu$m.

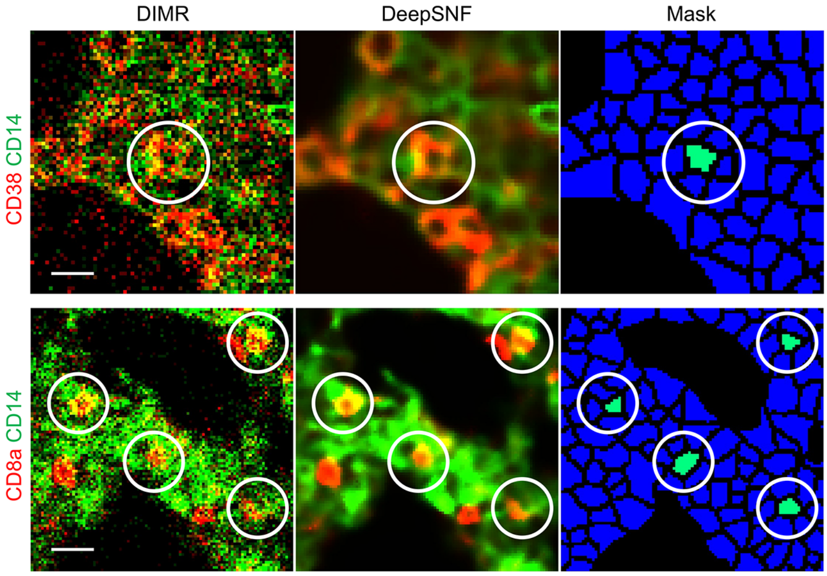

**Supplementary Figure 35.** Examples of double CD38+ CD14+ (top) and double CD8a+ CD14+ (bottom) cells. Scale bar: top: 11 $\mu$m, bottom: 16 $\mu$m.

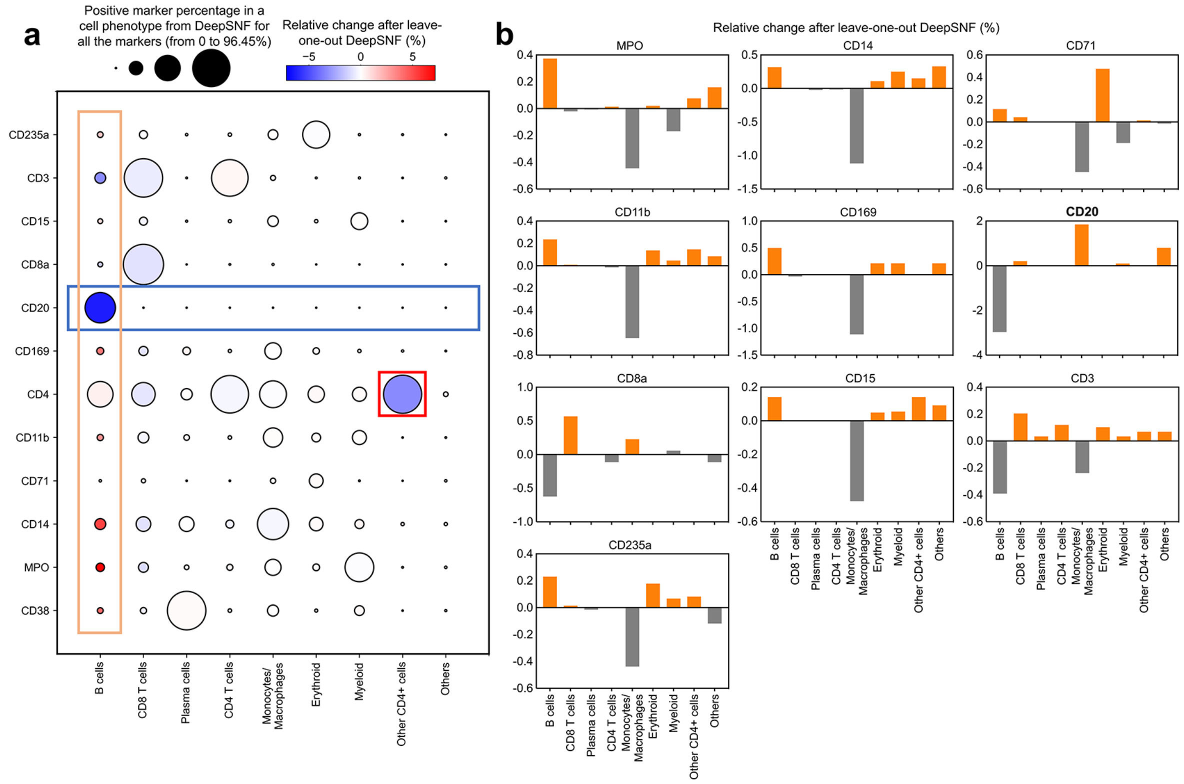

**Supplementary Figure 36.** The impact of CD20 denoising on cell phenotyping. Only CD20 was processed by DIMR while all the other markers were denoised by both DIMR and DeepSNF. Then the leave-one-out DeepSNF phenotyping result was compared to the DeepSNF for all the markers. (a) The sensitivity of cell phenotyping reduces for B cells. The specific marker signals reduce while the non-specific ones enrich in the B cells, respectively. The impacts on other cell types are limited compared to B cells. The circle size indicates the positive marker percentage in a particular phenotype from DeepSNF for all the markers, and the circle color indicates the relative changes of the positive rate for the particular markers after processed by the leave-one-out DeepSNF. (b) The specificity of cell phenotyping on CD20+ cells reduces. The ratio of B cells decreases while those of non-specific phenotypes increase in the CD20+ cells. The specificity of cell phenotyping on other positive markers, such as CD14 and CD169, also reduce, but the extent is limited.

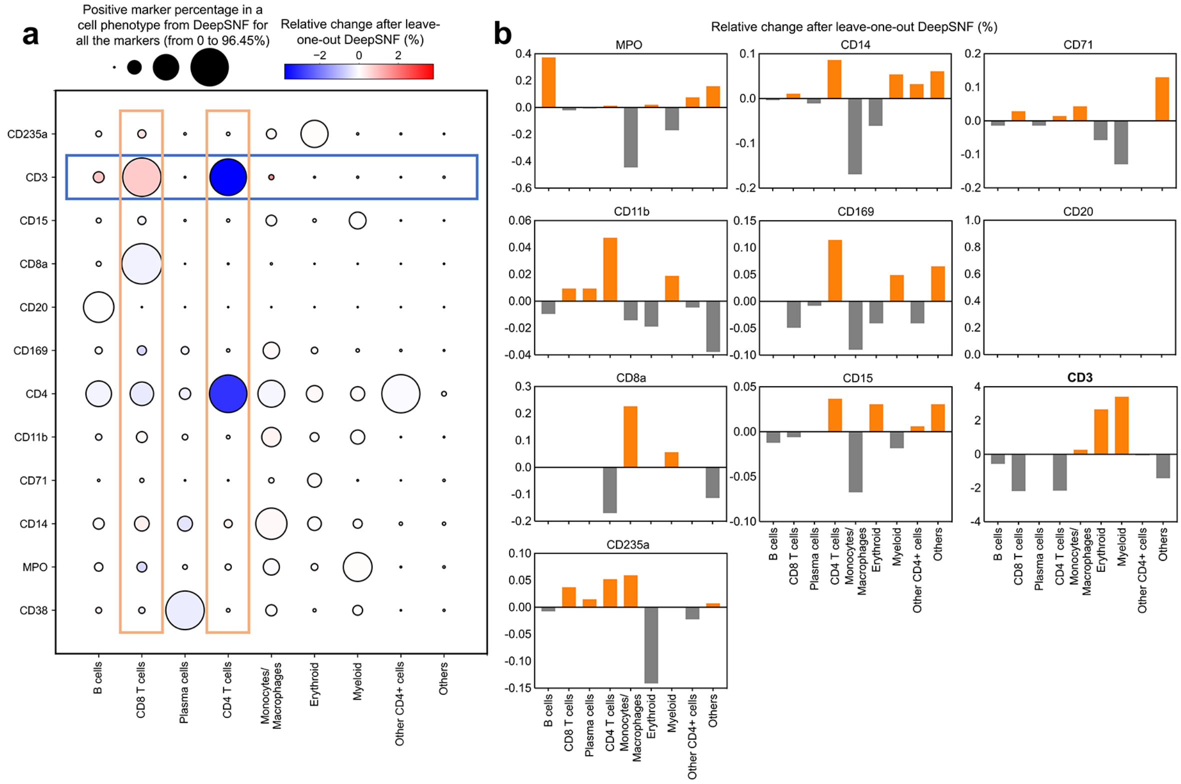

**Supplementary Figure 37.** The impact of CD3 denoising on cell phenotyping. Only CD3 was processed by DIMR while all the other markers were denoised by both DIMR and DeepSNF. Then the leave-one-out DeepSNF phenotyping result was compared to the DeepSNF for all the markers. (a) The sensitivity of cell phenotyping reduces for CD8 and CD4 T cells. The specific marker signals reduce while the non-specific ones enrich in the CD4 T cells, respectively. For CD8 T cells, even though the ratio of CD3+ cells increases, the ratio of CD8a+ cells decreases slightly. Overall, the sensitivity for CD8 T cells also decreases. The impacts on other cell types are limited. The circle size indicates the positive marker percentage in a particular phenotype from DeepSNF for all the markers, and the circle color indicates the relative changes of the positive rate for the particular markers after processed by the leave-one-out DeepSNF. (b) The specificity of cell phenotyping on CD3+ cells reduces. The ratios of CD8 and CD4 T cells decrease while those of non-specific phenotypes increase in the CD3+ cells. The specificity of cell phenotyping on other positive markers, such as MPO and CD8a, also reduces, but the extent is limited.

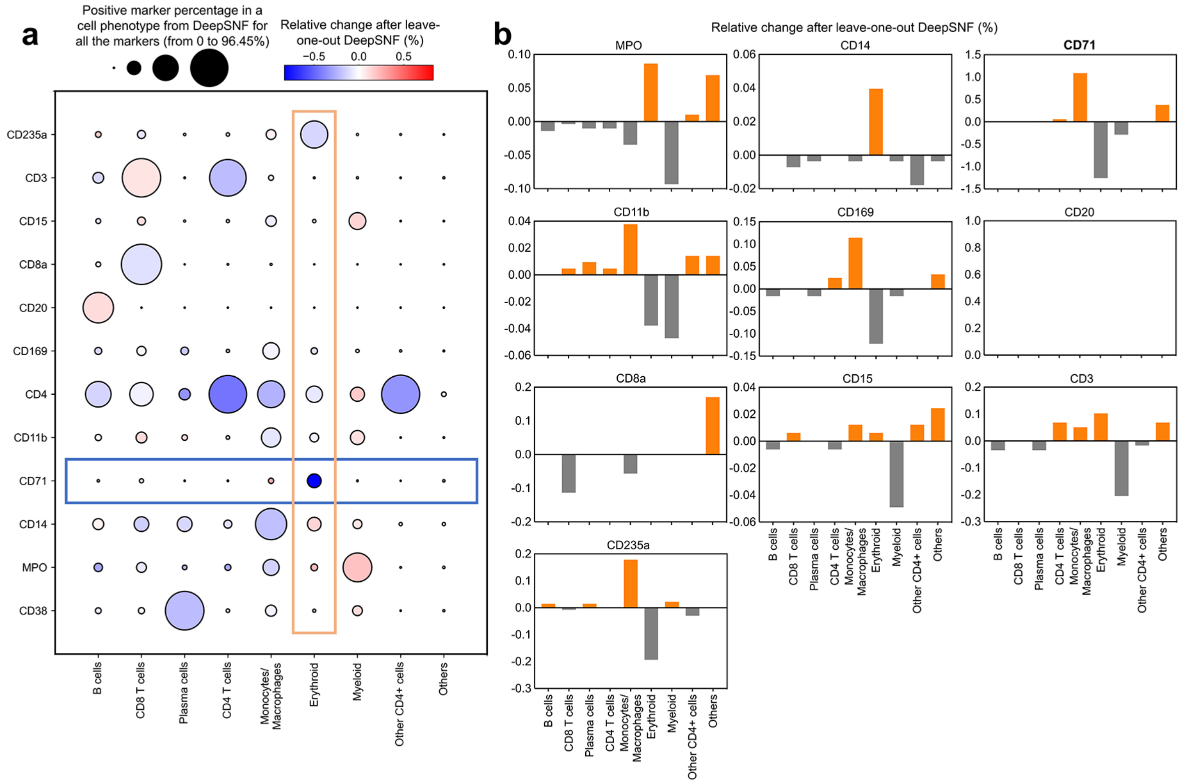

**Supplementary Figure 38.** The impact of CD71 denoising on cell phenotyping. Only CD71 was processed by DIMR while all the other markers were denoised by both DIMR and DeepSNF. Then the leave-one-out DeepSNF phenotyping result was compared to the DeepSNF for all the markers. (a) The sensitivity of cell phenotyping reduces for erythroid cells. The specific marker signals reduce while the non-specific ones enrich in the erythroids, respectively. The impacts on other cell types are limited compared to erythroid cells. The circle size indicates the positive marker percentage in a particular phenotype from DeepSNF for all the markers, and the circle color indicates the relative changes of the positive rate for the particular markers after processed by the leave-one-out DeepSNF. (b) The specificity of cell phenotyping on CD71+ cells reduces. The ratio of erythroids decreases while those of non-specific phenotypes increase in the CD71+ cells. Overall, the impact of CD71 denoising is smaller than those of CD20 and CD3, which corresponds to Supplementary Fig. 29(b).

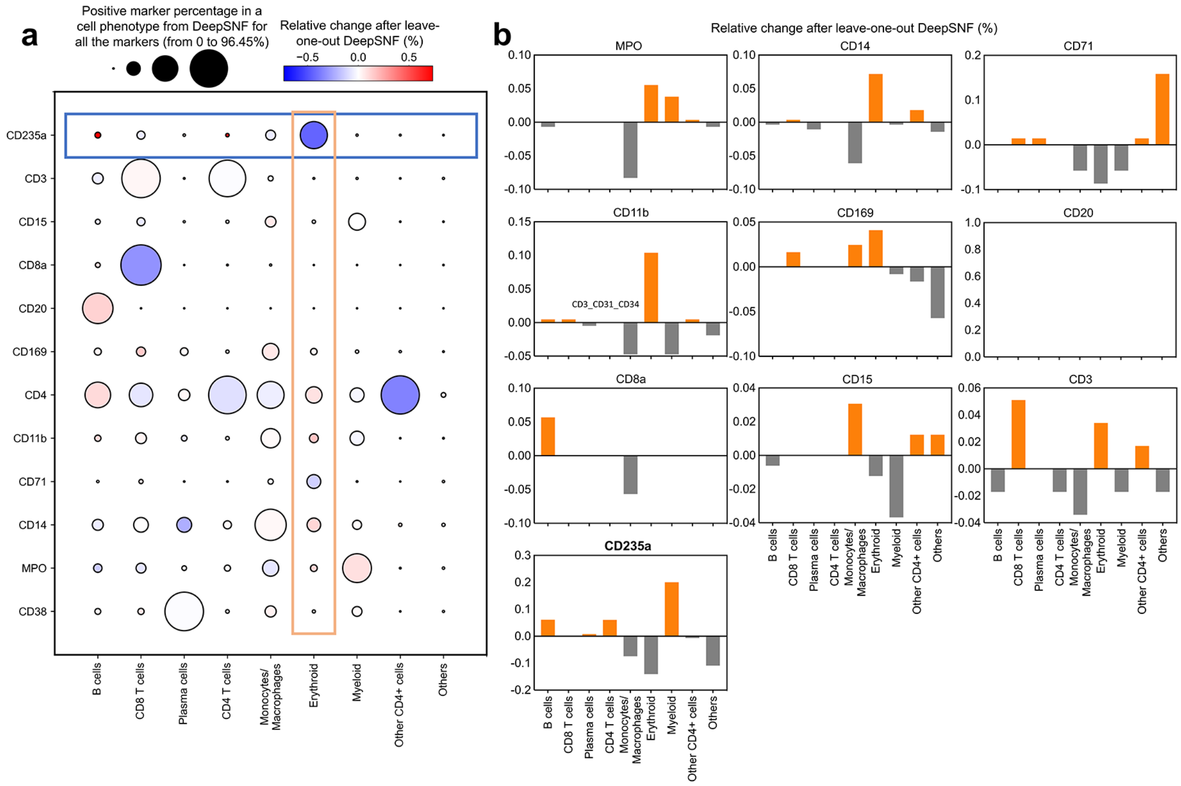

**Supplementary Figure 39.** The impact of CD235a denoising on cell phenotyping. Only CD235a was processed by DIMR while all the other markers were denoised by both DIMR and DeepSNF. Then the leave-one-out DeepSNF phenotyping result was compared to the DeepSNF for all the markers. (a) The sensitivity of cell phenotyping reduces slightly for erythroid cells. The specific marker signals reduce while the non-specific ones enrich in the erythroids, respectively. The circle size indicates the positive marker percentage in a particular phenotype from DeepSNF for all the markers, and the circle color indicates the relative changes of the positive rate for the particular markers after processed by the leave-one-out DeepSNF. (b) The specificity of cell phenotyping on CD235a+ cells reduces slightly. The ratio of erythroid cells decreases while those of non-specific phenotypes increase in the CD235a+ cells. Overall, the impact of CD235a denoising is limited compared to CD3, CD20 and CD71 because of the high SNR of CD235a IMC images, which corresponds to Supplmentary Fig. 29(b).

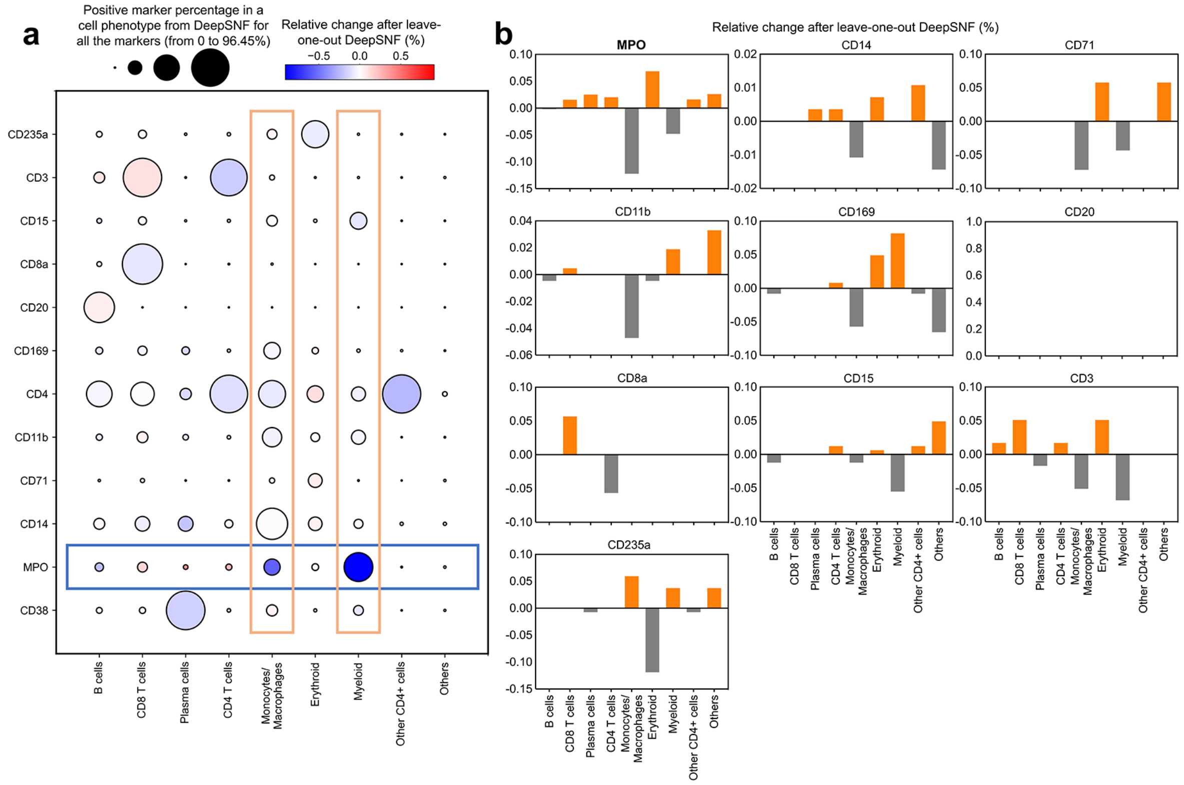

**Supplementary Figure 40.** The impact of MPO denoising on cell phenotyping. Only MPO was processed by DIMR while all the other markers were denoised by both DIMR and DeepSNF. Then the leave-one-out DeepSNF phenotyping result was compared to the DeepSNF for all the markers. (a) The sensitivity of cell phenotyping reduces slightly for myeloid cells and monocytes/macrophages. The specific marker signals reduce while the non-specific ones enrich in the myeloid cells and monocytes/macrophages, respectively. The circle size indicates the positive marker percentage in a particular phenotype from DeepSNF for all the markers, and the circle color indicates the relative changes of the positive rate for the particular markers after processed by the leave-one-out DeepSNF. (b) The specificity of cell phenotyping on MPO+ cells reduces slightly. The ratios of monocytes/macrophages and myeloid cells decrease while those of non-specific phenotypes increase in the MPO+ cells. Overall, the impact of MPO denoising is limited compared to CD3, CD20 and CD71 because of the high SNR of MPO IMC images, which corresponds to Supplmentary Fig. 29(b).

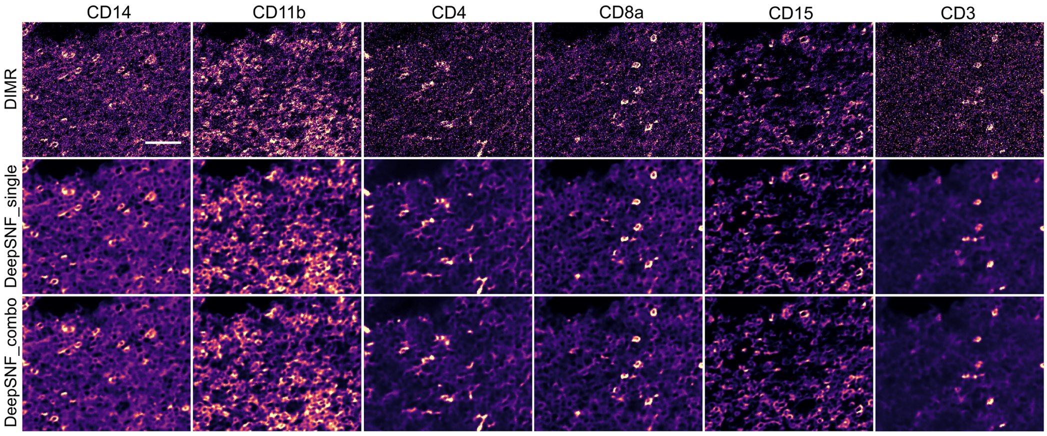

**Supplementary Figure 41.** DeepSNF works on multiple markers training. The DIMR-processed IMC images were trained by DeepSNF with single marker in each network (DeepSNF_single) and all the markers in a single network (DeepSNF_combo), respectively. The denoising results indicate both approaches enables IMC image quality improvement. Scale bar: 48 $\mu$m.

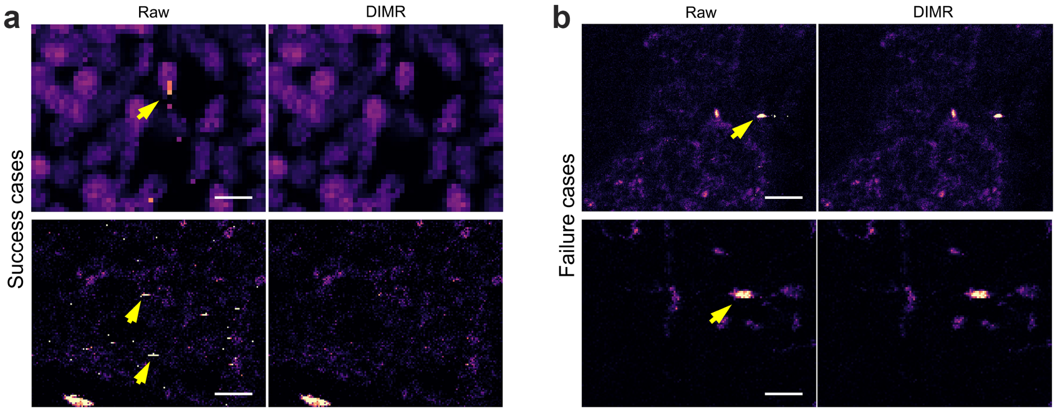

**Supplementary Figure 42.** The limitation of the DIMR algorithm. (a) Success cases of DIMR on challenging hot pixels. (b) Failure cases of DIMR on hot clusters. DIMR is able to remove line-style consecutive hot pixels while fails on hot clusters. Scale bar: (a) Top: 8 $\mu$m, bottom: 24 $\mu$m. (b) Top: 40 $\mu$m, bottom: 20 $\mu$m.

### Supplementary Tables

**Supplementary Table 2.** List of cell markers used for the Collagen III-labeled tissues in Fig. 1d and Fig. 2a

| Isotope | Metal | Epitope | Clone | Source | Catalog # | Dilution |
|---|---|---|---|---|---|---|
| 141 | Pr | CD235 | HIR2 | Fluidigm | 3141001B | 1:200 |
| 142 | Nd | MPO | polyclonal | Dako | A0398 | 1:800 |
| 144 | Nd | CD14 | EPR3653 | Fluidigm | 3144025D | 1:800 |
| 145 | Nd | CD117 | YR145 | Abcam | ab216450 | 1:100 |
| 147 | Sm | CD163 | EDHu-1 | Fluidigm | 3147021D | 1:100 |
| 148 | Nd | CD71 | MRQ-48 | eBiosciences | 14-0718-93 | 1:200 |
| 149 | Sm | CD11b | EPR1344 | Fluidigm | 3149028D | 1:150 |
| 151 | Eu | CD31 | EPR3094 | Fluidigm | 3151025D | 1:100 |
| 152 | Sm | CD34 | QBend/10 | ThermoFisher | MA1-10202 | 1:400 |
| 153 | Eu | pSTAT5 | 47 | BD | custom quote | 1:100 |
| 154 | Sm | TNFa | TNF706 + P/T2 | Abcam | ab212899 | 1:150 |
| 155 | Gd | IL8 | 807 | Abcam | custom quote | 1:200 |
| 156 | Gd | CD4 | EPR6855 | Fluidigm | 3156033D | 1:500 |
| 157 | Gd | IL6 | 1936 | R&D | MAB2061 | 1:100 |
| 158 | Gd | pSTAT3 | 4/P-STAT3 | Fluidigm | 3158030D | 1:100 |
| 159 | Tb | CD90 | 5.00E+10 | Fluidigm | 3159007B | 1:100 |
| 160 | Gd | CD61 | 2f2 | Sigma | custom quote | 1:400 |
| 161 | Dy | CD20 | H1 | Fluidigm | 3161029D | 1:400 |
| 162 | Dy | CD8a | C8/144B | Fluidigm | 3162034D | 1:300 |
| 163 | Dy | TGFb | TB21 | Invitrogen | MA1-21595 | 1:800 |
| 164 | Dy | CD15 | W6D3 | Fluidigm | 3164001B | 1:150 |
| 165 | Ho | pCREB | 87G3 | Fluidigm | 3165009A | 1:400 |
| 166 | Er | NFkBp65 pS529 | K10x | Fluidigm | 3166006A | 1:200 |
| 167 | Er | RELA | 2A12A7 | ThermoFisher | 33-9900 | 1:100 |
| 168 | Er | Ki-67 | B56 | Fluidigm | 3168022D | 1:200 |
| 169 | Tm | Pikk1/2 I | 16A6 | CST | 2697BF | 1:200 |
| 170 | Er | CD3 | polyclonal | Fluidigm | 3170019D | 1:200 |

**Table 2 continued from previous page**

| | | | | | | |
|---|---|---|---|---|---|---|
| 172 | Yb | Cleaved casp 3 | 5A1E | Fluidigm | 3172027D | 1:300 |
| 174 | Yb | pERK1/2 | D13.14.4E | Fluidigm | 3171021D | 1:100 |
| 175 | Lu | pS6 | N7-548 | Fluidigm | 3175009A | 1:400 |
| 176 | Yb | Histone-H3 | D1H2 | Fluidigm | 3176023D | 1:2000 |
| 209 | Bi | Collagen III | polyclonal | Southern Biotech | 1330-01 | 1:100 |
| 191/193 | Ir | intercalator | | | | 1:300 |

**Supplementary Table 3.** List of cell markers for the tissue staining in Fig. 1g

| Isotope | Metal | Epitope | Clone | Source | Catalog # | Dilution |
|---|---|---|---|---|---|---|
| 142 | Nd | MPO | polyclonal | Dako | A0398 | 1:400 |
| 145 | Nd | CD117 | YR145 | Abcam | ab216450 | 1:50 |
| 150 | Nd | CXCL12 | 79018 | Novus | MAB350 | 1:75 |
| 153 | Eu | IFNg | IFNG/466 | Novus | NBP2-54394 | 1:25 |
| 154 | Sm | TNFa | TNF706 + P/T2 | Abcam | ab212899 | 1:75 |
| 155 | Gd | IL8 | 807 | Abcam | custom quote | 1:25 |
| 157 | Gd | IL6 | 1936 | R&D | MAB2061 | 1:25 |
| 158 | Gd | pSmad4 | polyclonal | ThermoFisher | PA5-12695 | 1:150 |
| 159 | Tb | CD169 | SP213 | Abcam | ab245735 | 1:100 |
| 163 | Dy | CD271 | EP1039Y | Abcam | ab256584 | 1:50 |
| 173 | Yb | Collagen III | polyclonal | Southern Biotech | 1330-01 | 1:400 |
| 209 | Bi | Collagen III | polyclonal | Southern Biotech | 1330-01 | 1:200 |
| 191/193 | Ir | intercalator | | | | 1:200 |

**Supplementary Table 4.** List of cell markers for the tissue staining in Supplementary Fig. 20

| Isotope | Metal | Epitope | Clone | Source | Catalog # | Dilution |
|---|---|---|---|---|---|---|
| 142 | Nd | MPO | polyclonal | Dako | A0398 | 1:400 |
| 143 | Nd | TP53 | DO-7 | Fluidigm | 3143026D | 1:50 |
| 146 | Nd | TP53 | DO-7 | Biolegend | 345102 | 1:50 |
| 150 | Nd | CXCL12 | 79018 | Novus | MAB350 | 1:50 |
| 158 | Gd | CD56 | MRQ-42 | CellMarque | custom quote | 1:100 |
| 163 | Dy | CD271 | EP1039Y | Abcam | ab256584 | 1:50 |
| 167 | Er | GranzymeB | EPR20129-217 | Fluidigm | 3167021D | 1:600 |
| 173 | Yb | CD56 | MRQ-42 | CellMarque | custom quote | 1:100 |
| 191/193 | Ir | intercalator | | | | 1:300 |

**Supplementary Table 5.** List of cell markers used for other IMC images from the human bone marrow IMC dataset

| Isotope | Metal | Epitope | Clone | Source | Catalog # | Dilution |
|---|---|---|---|---|---|---|
| 89 | Yb | Alpha-SMA | 1A4 | Bio-Rad | MCA5781GA | 1:100 |
| 115 | In | perilipin | D1D8 | CST | 9349 custom | 1:50 |
| 139 | La | VCAM1 | EPR5047 | Abcam | ab215380 | 1:50 |
| 141 | Pr | CD38 | EPR4106 | Fluidigm | 3141018D | 1:50 |
| 142 | Nd | MPO | polyclonal | Dako | A0398 | 1:400 |
| 143 | Nd | vimentin | RV202 | Fluidigm | 3143029D | 1:200 |
| 144 | Nd | CD14 | EPR3653 | Fluidigm | 3144025D | 1:400 |
| 145 | Nd | CD117 | YR145 | Abcam | ab216450 | 1:50 |
| 146 | Nd | CD16 | EPR16784 | Fluidigm | 3146020D | 1:150 |
| 147 | Sm | CD163 | EDHu-1 | Fluidigm | 3147021D | 1:100 |
| 148 | Nd | CD71 | MRQ-48 | eBiosciences | 14-0718-93 | 1:50 |
| 149 | Sm | CD11b | EPR1344 | Fluidigm | 3149028D | 1:150 |
| 150 | Nd | CXCL12 | 79018 | Novus | MAB350 | 1:25 |
| 151 | Eu | CD31 | EPR3094 | Fluidigm | 3151025D | 1:50 |
| 152 | Sm | CD34 | QBend/10 | ThermoFisher | MA1-10202 | 1:50 |
| 153 | Eu | IFNg | IFNG/466 | Novus | NBP2-54394 | 1:25 |
| 154 | Sm | TNFa | TNF706 + P/T2 | Abcam | ab212899 | 1:100 |
| 156 | Gd | CD4 | EPR6855 | Fluidigm | 3156033D | 1:200 |
| 157 | Gd | IL6 | 1936 | R&D | MAB2061 | 1:25 |
| 158 | Gd | pSmad4 | polyclonal | ThermoFisher | PA5-12695 | 1:150 |
| 159 | Tb | CD169 | SP213 | Abcam | ab245735 | 1:100 |
| 160 | Gd | CD61 | 2f2 | Sigma | custom quote | 1:100 |
| 161 | Dy | CD20 | H1 | Fluidigm | 3161029D | 1:400 |
| 162 | Dy | CD8a | C8/144B | Fluidigm | 3162034D | 1:300 |
| 163 | Dy | CD271 | EP1039Y | Abcam | ab256584 | 1:50 |
| 164 | Dy | CD15 | W6D3 | Fluidigm | 3164001B | 1:150 |
| 165 | Ho | pH2AX | N1-431 | Fluidigm | 3165036D | 1:150 |
| 166 | Er | NFkBp65 pS529 | K10x | Fluidigm | 3166006A | 1:25 |
| 167 | Er | SCF | polyclonal | ThermoFisher | PA5-20746 | 1:25 |

**Table 5 continued from previous page**

| | | | | | | |
|---|---|---|---|---|---|---|
| 168 | Er | Ki-67 | B56 | Fluidigm | 3168022D | 1:100 |
| 169 | Tm | Collagen I | polyclonal | Fluidigm | 3169023D | 1:2000 |
| 170 | Er | CD3 | polyclonal | Fluidigm | 3170019D | 1:100 |
| 171 | Yb | pERK1/2 | D13.14.4E | Fluidigm | 3171021D | 1:50 |
| 172 | Yb | Cleaved casp 3 | 5A1E | Fluidigm | 3172027D | 1:25 |
| 173 | Yb | CD45RO | UCHL1 | Fluidigm | 3173016D | 1:500 |
| 174 | Yb | HLA-DR | YE2/36HLK | Fluidigm | 3174023D | 1:100 |
| 175 | Lu | CD235a | HIR2 | Fluidigm | 3175029D | 1:200 |
| 176 | Yb | Histone-H3 | D1H2 | Fluidigm | 3176023D | 1:2000 |
| 209 | Bi | Collagen III | polyclonal | Southern Biotech | 1330-01 | 1:75 |
| 191/193 | Ir | intercalator | | | | 1:200 |

Note: The tissues with headers of K, L do not have CXCL12.

**Supplementary Table 6.** Training details for the simulation datasets

| #Patches | Normalzed percentile | training time |
|---|---|---|
| 12000 | 99.999 | 89 min |

**Supplementary Table 7.** Training details for the Collagen III-labeled images in Fig. 1d, g and Fig.2a

| Marker | #Patches | Normalzed percentile | Background thresh $\rho$ | training time |
|---|---|---|---|---|
| Collagen III | 1992 | 99.9 | 0.55 | 17 min |

**Supplementary Table 8.** Training details for the other markers-labeled images from the human bone marrow IMC dataset

| Marker | #Patches | Normalzed percentile | Background thresh $\rho$ | training time |
|---|---|---|---|---|
| CD38 | 21768 | 99.999 | 0.9 | 160 min |
| MPO | 20800 | 99.999 | 0.8 | 153 min |
| CD14 | 21784 | 99.999 | 0.9 | 160 min |
| CD71 | 14960 | 99.999 | 0.9 | 110 min |
| CD11b | 20096 | 99.999 | 0.9 | 147 min |
| CD31 | 9040 | 99.99 | 0.75 | 67 min |
| CD34 | 15208 | 99.9 | 0.85 | 114 min |
| CD4 | 20832 | 99.9 | 0.9 | 154 min |
| CD169 | 20800 | 99.999 | 0.9 | 153 min |
| CD61 | 3360 | 99.9 | 0.75 | 27 min |
| CD20 | 12304 | 99.9 | 0.95 | 90 min |
| CD8a | 19360 | 99.999 | 0.9 | 144 min |
| CD15 | 17456 | 99.99 | 0.9 | 127 min |
| Ki-67 | 18032 | 99.999 | 0.9 | 134 min |
| CD3 | 16728 | 99.99 | 0.9 | 124 min |
| CD45RO | 11672 | 99.99 | 0.75 | 87 min |
| CD235a | 21144 | 99.999 | 0.7 | 154 min |
| Histone H3 | 14952 | 99.999 | 0.5 | 108 min |
| DNA2 | 22136 | 99.999 | 0.4 | 161 min |
| Combinations of CD4, CD8a, CD3, CD14, CD11b, CD71 and CD15 | 131216 | 99.9 | | 15.8 h |

**Supplementary Table 9.** Training details for the images from human breast cancer IMC dataset

| Marker | #Patches | Normalzed percentile | Background thresh $\rho$ | training time |
|---|---|---|---|---|
| CD3 | 12592 | 99.9 | 0.95 | 94 min |
| CD20 | 17592 | 99.9 | 0.95 | 133 min |
| CD45 | 9368 | 99.999 | 0.9 | 72 min |
| CD68 | 13024 | 99.999 | 0.85 | 97 min |
| c-Myc | 14056 | 99.999 | 0.85 | 104 min |
| EGFR | 17624 | 99.999 | 0.85 | 135 min |
| EpCAM | 15064 | 99.999 | 0.8 | 112 min |
| Ki-67 | 10824 | 99.999 | 0.9 | 80 min |
| Rabbit IgG H L | 10928 | 99.999 | 0.9 | 84 min |
| Slug | 11856 | 99.999 | 0.9 | 90 min |
| Twist | 14496 | 99.999 | 0.9 | 107 min |
| vWF | 17928 | 99.999 | 0.95 | 136 min |

**Supplementary Table 10.** Training details for the images from the human pancreatic cancer IMC dataset

| Marker | #Patches | Normalzed percentile | Background thresh $\rho$ | training time |
|---|---|---|---|---|
| CD3 | 16304 | 99.99 | 0.75 | 120 min |
| CD4 | 14344 | 99.999 | 0.7 | 107 min |
| CD8 | 9792 | 99.9 | 0.77 | 74 min |
| CD11b | 21120 | 99.99 | 0.3 | 154 min |
| CD14 | 10896 | 99.99 | 0.5 | 82 min |
| CD31 | 17088 | 99.99 | 0.75 | 126 min |
| CD44 | 17392 | 99.99 | 0.5 | 127 min |
| CD45 | 21984 | 99.99 | 0.7 | 161 min |
| CD45RO | 18056 | 99.99 | 0.6 | 134 min |
| CD56 | 11168 | 99.99 | 0.5 | 84 min |
| Foxp3 | 9240 | 99.99 | 0.4 | 70 min |
| pS6 | 19440 | 99.99 | 0.2 | 144 min |

**Supplementary Table 11.** Training details for the images from the IMC dataset

| Marker | #Patches | Normalzed percentile | Background thresh $\rho$ | training time |
|---|---|---|---|---|
| CD3 | 5096 | 99.999 | 0.95 | 43 min |
| CD4 | 4120 | 99.999 | 0.95 | 34 min |
| CD8 | 7472 | 99.999 | 0.98 | 57 min |
| CD11b | 11400 | 99.999 | 0.95 | 87 min |
| CD11c | 6720 | 99.999 | 0.96 | 50 min |
| CD14 | 9768 | 99.999 | 0.95 | 74 min |
| CD20 | 13184 | 99.999 | 0.95 | 97 min |
| CD31 | 7272 | 99.999 | 0.99 | 57 min |
| CD45 | 13792 | 99.999 | 0.95 | 104 min |
| CD68 | 8368 | 99.999 | 0.98 | 64 min |
| CD206 | 10672 | 99.999 | 0.98 | 80 min |
| HLA-DR | 13296 | 99.999 | 0.95 | 100 min |

**Supplementary Table 12.** The estimated thresholds for positive markers

| Marker | DIMR | DeepSNF | Marker | DIMR | DeepSNF |
|---|---|---|---|---|---|
| CD38 | 1.3981 | 1.4037 | CD169 | 2.5625 | 2.5698 |
| MPO | 4.2308 | 4.2200 | CD20 | 0.8000 | 0.7347 |
| CD14 | 2.3949 | 2.4272 | CD8a | 3.3559 | 3.2336 |
| CD71 | 0.9478 | 0.9438 | CD15 | 2.6698 | 2.6461 |
| CD11b | 2.1194 | 2.1504 | CD3 | 0.9180 | 0.8866 |
| CD4 | 0.7830 | 0.8140 | CD235a | 35.7895 | 35.0695 |

### References

[1] S. Chevrier, H. L. Crowell, V. R. Zanotelli, S. Engler, M. D. Robinson, B. Bodenmiller, Compensation of signal spillover in suspension and imaging mass cytometry, Cell Systems 6 (5) (2018) 612–620.

[2] P. Lu, N. Benabdallah, W. Jiang, B. W. Simons, H. Zhang, R. F. Hobbs, D. Ulmert, B. Baumann, R. K. Pachynski, A. K. Jha, et al., Blind image restoration enhances digital autoradiographic imaging of radiopharmaceutical tissue distribution, Journal of Nuclear Medicine (2021).

[3] F. J. Anscombe, The transformation of poisson, binomial and negative-binomial data, Biometrika 35 (3/4) (1948) 246–254.

[4] S. K. Bar-Lev, P. Enis, On the classical choice of variance stabilizing transformations and an application for a poisson variate, Biometrika 75 (4) (1988) 803–804.

[5] M. F. T. B. C. Russell, W. T. Freeman, Exploiting the sparse derivative prior for super-resolution and image demosaicing, in: Proceedings of the Third International Workshop Statistical and Computational Theories of Vision, 2003, pp. 1–28.

[6] W. Rudin, et al., Principles of mathematical analysis, Vol. 3, McGraw-hill New York, 1964.

[7] B. W. Silverman, Density estimation for statistics and data analysis, Routledge, 2018.

[8] M. Makitalo, A. Foi, Optimal inversion of the anscombe transformation in low-count poisson image denoising, IEEE transactions on Image Processing 20 (1) (2010) 99–109.

[9] L. Finesso, P. Spreij, Nonnegative matrix factorization and i-divergence alternating minimization, Linear Algebra and its Applications 416 (2-3) (2006) 270–287.

[10] A. Krull, T.-O. Buchholz, F. Jug, Noise2void-learning denoising from single noisy images, in: Proceedings of the IEEE/CVF Conference on Computer Vision and Pattern Recognition, 2019, pp. 2129–2137.

[11] J. Batson, L. Royer, Noise2self: Blind denoising by self-supervision, in: International Conference on Machine Learning, PMLR, 2019, pp. 524–533.

[12] X. Huang, J. Fan, L. Li, H. Liu, R. Wu, Y. Wu, L. Wei, H. Mao, A. Lal, P. Xi, et al., Fast, long-term, super-resolution imaging with hessian structured illumination microscopy, Nature biotechnology 36 (5) (2018) 451–459.

[13] W. Zhao, S. Zhao, L. Li, X. Huang, S. Xing, Y. Zhang, G. Qiu, Z. Han, Y. Shang, D.-e. Sun, et al., Sparse deconvolution improves the resolution of live-cell super-resolution fluorescence microscopy, Nature biotechnology (2021) 1–12.

[14] V. Zanotelli, B. Bodenmiller, Imc segmentation pipeline: a pixel classification based multiplexed image segmentation pipeline, Zenodo https://doi. org/10.5281/zenodo 3841960 (2017).

[15] A. F. Rendeiro, H. Ravichandran, Y. Bram, V. Chandar, J. Kim, C. Meydan, J. Park, J. Foox, T. Hether, S. Warren, et al., The spatial landscape of lung pathology during covid-19 progression, Nature (2021) 1–6.

[16] M. Wu, M. Y. Lee, V. Bahl, D. Traum, J. Schug, I. Kusmartseva, M. A. Atkinson, G. Fan, K. H. Kaestner, H. Consortium, et al., Single-cell analysis of the human pancreas in type 2 diabetes using multi-spectral imaging mass cytometry, Cell reports 37 (5) (2021) 109919.

[17] Y. J. Wang, D. Traum, J. Schug, L. Gao, C. Liu, M. A. Atkinson, A. C. Powers, M. D. Feldman, A. Naji, K.-M. Chang, et al., Multiplexed in situ imaging mass cytometry analysis of the human endocrine pancreas and immune system in type 1 diabetes, Cell metabolism 29 (3) (2019) 769–783.

[18] B. Xu, N. Wang, T. Chen, M. Li, Empirical evaluation of rectified activations in convolutional network, arXiv preprint arXiv:1505.00853 (2015).

[19] M. Weigert, U. Schmidt, T. Boothe, A. Müller, A. Dibrov, A. Jain, B. Wilhelm, D. Schmidt, C. Broaddus, S. Culley, et al., Content-aware image restoration: pushing the limits of fluorescence microscopy, Nature methods 15 (12) (2018) 1090–1097.

[20] A. Buades, B. Coll, J.-M. Morel, A non-local algorithm for image denoising, in: 2005 IEEE Computer Society Conference on Computer Vision and Pattern Recognition (CVPR'05), Vol. 2, IEEE, 2005, pp. 60–65.

[21] K. Dabov, A. Foi, V. Katkovnik, K. Egiazarian, Image denoising by sparse 3-d transform-domain collaborative filtering, IEEE Transactions on image processing 16 (8) (2007) 2080–2095.

[22] R. Rashid, G. Gaglia, Y.-A. Chen, J.-R. Lin, Z. Du, Z. Maliga, D. Schapiro, C. Yapp, J. Muhlich, A. Sokolov, et al., Highly multiplexed immunofluorescence images and single-cell data of immune markers in tonsil and lung cancer, Scientific data 6 (1) (2019) 1–10.

[23] Z. Wang, A. C. Bovik, H. R. Sheikh, E. P. Simoncelli, Image quality assessment: from error visibility to structural similarity, IEEE transactions on image processing 13 (4) (2004) 600–612.
